# Ultrasensitive single-genome sequencing reveals strong purifying selection in acute HIV- 1 infection

**DOI:** 10.64898/2026.08.21.746200

**Authors:** Adam A. Capoferri, Valerie F. Boltz, Wei Shao, Clarissa Halpern, Rasmi Thomas, Nittaya Phanuphak, Lydie Trautmann, Sandhya Vasan, Carlo Sacdalan, Somchai Sriplienchan, John W. Mellors, John M. Coffin, Jason W. Rausch, Mary F. Kearney, the RV254/SEARCH 010 Study Team

## Abstract

HIV transmission from one individual to another occurs by one or a small number of virions followed by spread and genetic diversification into a complex quasispecies. To understand the early events in this process, we investigated how HIV-1 genomes diversify within the first two to three weeks after transmission by use of ultra-deep single subgenomic sequencing of over 10,000 plasma RNA genomes in each of a cohort of 15 individuals in acute infection. This approach confirmed transmission of one or a few transmitted/founder (TF) viral lineages and very limited early divergence from the founder sequences. Most observed variants that differed from the TF included single nucleotide changes attributable to HIV-1 reverse transcriptase (RT) error or host APOBEC3G/F activity. Comparing the number of expected versus observed changes after transmission indicated that most *de novo* mutations do not persist in the virus population, consistent with strong purifying selection. We found little evidence that early diversification is driven by reversions to subtype consensus or by cytotoxic T lymphocyte pressure, although rare multi-mutation lineages suggest occasional influences. Together, these findings indicate that early HIV-1 evolution is influenced by stochastic and host-mediated mutational processes (*e.g.*, APOBEC3G/F) filtered by strong purifying selection. The strong purifying selection observed in the early weeks of HIV-1 infection may provide an opportunity to investigate the potential of new interventions to induce viremic control, such as combinations of broadly neutralizing antibodies, cellular immunotherapy, or mRNA therapeutic vaccination.

**AUTHOR SUMMARY:** When a person acquires HIV, especially through sexual transmission, infection is usually established by just one or a few viral variants. However, over the early months and years of infection, these viral variants accumulate mutations until almost no two viral genomes are identical in a typical sample. Here, we sequenced tens of thousands of viral variants in the early weeks after transmission to understand the early events that contribute to this vast viral diversification. We found that the accumulation of mutations was slower than expected, implying a selection against HIV-1 diversification in acute infection, potentially leaving a window of low genetic diversity for the study of new interventions towards inducing viremic control, such as immunotherapy or mRNA vaccination. Of the early viral mutations that were observed, many were induced by host enzymes, rather than from errors by the viral enzyme used for replication. Our results provide more context for understanding HIV evolution and provide a deep sampling of viral diversity after transmission.

## INTRODUCTION

In an HIV-1 transmission event from an individual in chronic infection, the recipient is likely exposed to a diverse population of viral quasispecies. However, shortly after exposure, this viral diversity is often dramatically reduced resulting in a severe genetic bottleneck, thought to be imposed, in part, from anatomical barriers (*1–7*). The variants inferred to establish a new infection are referred to as the transmitted/founder (TF) virus(es), and they dominate the early stages of viral spread (*1, 2, 6–10*). In most sexually transmitted cases (75-80%) (*2, 9, 11–13*), infection is established by a single TF variant as transmission fluids enter a restrictive mucosal environment, compounded by local immune pressures (*14–16*). Infections founded by multiple TF variants may reflect a less stringent bottleneck where multiple variants transverse mucosal barriers (*2–4, 17–19*) or enter through other exposure routes (*e.g.*, 60% of transmission via injected drugs) (*20*).

HIV-1 acute infection and disease progression is staged by the Fiebig classification system based on the sequential appearance of viral RNA, capsid (p24) antigen, antibodies, and protein detected by western blot (WB) (*21*). Acute infection is typically defined as the first 30 days following transmission (*i.e.*, before a full WB^+^ profile) and represents a critical window for potential curative interventions, owing to the small reservoir size and generally limited viral diversity. During the course of HIV-1 infection, within-host viral populations undergo extensive genetic diversification (*6, 7, 10*) driven by large population sizes, high mutation rates, recombination, and immune selection (*1, 22–24*). Longitudinal analyses of *env* sequences show a steady accumulation of diversity with periodic replacement of dominant lineages as the virus repeatedly escapes host immunity (*6, 7, 10, 17*), often accompanied by phenotypic shifts such as changes in neutralization sensitivity or coreceptor usage (*1, 25*). This trajectory often manifests as acute infection harboring predominantly CCR5-tropic homogeneous sequences, with chronic infection sometimes including CXCR4-tropic variants and increased heterogeneity (*17, 25*). Correspondingly, diversity among early viral genomic sequences is low relative to the heterogeneous quasispecies characteristic of chronic infection (*6, 7, 10*).

The genetic composition of the viral population during acute infection has been characterized using single-genome sequencing (SGS) and next-generation sequencing (NGS). SGS typically involves sequencing 10–100 individual viral genomes (*2, 5, 9*), while NGS can be used to sequence subgenomic or full-length gene regions from hundreds or thousands of viral templates (*26–29*). The latter approach can be used to generate consensus HIV sequences from bulk populations, as well as estimates of the genetic diversity and allele frequencies in these samples. These methods have been instrumental for identifying transmission pairs and recency of source partners (*8, 16, 30–32*), inferring TF virus sequences (*2, 5, 9, 33, 34*), and evaluating baseline resistance to broadly neutralizing antibodies (*35–40*) or antiretroviral therapy (ART) (*41–45*). However, these approaches can have limited capacity for detection of low-frequency variants (*e.g.*, SGS) and establishing linkage between mutations due to the obfuscating effects of PCR recombination (*e.g.*, NGS) and resampling.

To address these deficiencies, we applied the ultrasensitive SGS (uSGS) technique we develped (*46*) to plasma samples collected from a cohort of individuals in acute infection with previously determined TF virus sequences from near-full length SGS of 8-10 viral genomes. uSGS is an NGS-based method that eliminates the problems of PCR recombination, PCR error, and sequencing artifacts through stringent amplification conditions and bioinformatic filtering, and delivers very large, high-fidelity datasets that accurately reflect the true composition of the viral populations. In the first step of this workflow, viral RNA is reverse transcribed using a gene-specific primer that contains both a primer ID (*i.e.*, a random sequence that serves as unique molecular identifier, UMI) and a constant region for selective PCR amplification. This procedure enables unique labeling and amplification of individual cDNA molecules. Hence, unlike conventional NGS methods where sequencing may reflect PCR resampling and recombinants rather than distinct genomes, uSGS ensures that each high-quality consensus sequence accurately corresponds to a unique viral genome. Moreover, by collapsing reads to UMI-defined templates and filtering artifacts, uSGS reduces resampling bias and improves minority-variant fidelity.

Here, we applied uSGS to plasma samples from people with HIV (PWH) in acute infection with previously defined TF designation. This approach enabled deep sampling of thousands of viral genomes to define the earliest diversification of HIV-1 after transmission. We reasoned that if mutation accumulation during acute infection was driven largely as a consequence of HIV-1 reverse-transcriptase error alone, then low-frequency variants would emerge in a predictable manner. Conversely, deviation from that expectation would suggest strong pressure constraints on early viral evolution. Through this framework, we evaluated TF designation, and quantified and characterized the earliest mutations arising post-transmission. Together, these analyses enabled us to determine the extent to which early HIV-1 diversification is generated by mutations, yet restricted under purifying selection pressure.

## RESULTS

### Donor characteristics

To genetically characterize single and multiple TF viruses during the acute stage of HIV-1 infection, frozen plasma samples were obtained from 15 participants enrolled in the RV254/SEARCH 010 study (NCT00796146) (*47*) (**Table 1**). The 15 samples were selected based on those that had previously undergone near full-length standard SGS of ∼10 genomes (*48*). Participants were predominately male with a median age of 29 years and recently acquired HIV- 1 CRF01_AE. All samples were collected during acute infection: Fiebig stage II (n=4), III (n=10), and IV (n=1). Participants had a median level of plasma viremia of 6.9 log_10_ copies HIV RNA/mL with CD4 counts of 289 cells/mm^3^. MHC-I HLA alleles were previously determined (*49, 50*).

**Table 1.** Participant characteristics.

| Participant ID (PID) | Sex<sup>a</sup> | Age at sampling (years) | Fiebig Stage at sampling<sup>b</sup> | Viral load at sampling (copies/mL)<sup>c</sup> | CD4 count at sampling (cells/mm<sup>3</sup>) | MHC-I HLA alleles<sup>d</sup> |  |  |
| --- | --- | --- | --- | --- | --- | --- | --- | --- |
|  |  |  |  |  |  | A1*/ A2* | B1*/ B2* | C1*/ C2* |
| 7279 | M | 27 | II | 7,249,767 | 234 | 02:01/ 02:07 | 40:01/ 46:01 | 01:02/ 07:02 |
| 6340 | M | 29 | II | 23,317,000 | 338 | 03:01/ 24:02 | 07:02/ 35:03 | 04:01/ 07:02 |
| 7905 | M | 29 | IV | 7,263,860 | 213 | 02:07/ 11:01 | 15:25/ 40:01 | 04:03/ 07:02 |
| 9813 | M | 22 | II | 5,163,823 | 165 | 11:01/ 11:01 | 15:02/ 38:02 | 07:02/ 08:01 |
| 7268 | M | 23 | III | 8,914,961 | 334 | 11:01/ 11:01 | 15:02/ 15:45 | 08:01/ 12:02 |
| 3928 | M | 46 | III | 5,517,440 | 525 | 11:01/ 11:01 | 15:02/ 15:02 | 08:01/ 08:01 |
| 3832 | M | 36 | II | 36,694,000 | 269 | 02:07/ 33:03 | 44:03/ 46:01 | 01:02/ 07:01 |
| 5436 | M | 32 | III | 30,811,000 | 621 | 11:03/ 33:03 | 44:03/ 52:01 | 07:01/ 07:02 |
| 3698 | M | 30 | III | 4,939,160 | 352 | 02:03/ 11:01 | 18:02/ 35:05 | 04:01/ 07:04 |
| 6609 | M | 29 | III | 4,112,500 | 206 | 24:02/ 33:03 | 07:05/ 51:01 | 07:02/ 14:02 |
| 8123 | F | 45 | III | 25,579,700 | 132 | 02:03/ 11:01 | 13:01/ 35:03 | 03:04/ 04:01 |
| 3513 | M | 28 | III | 13,557,900 | 359 | 11:01/ 11:01 | 18:01/ 40:06 | 07:01/ 15:02 |
| 9114 | M | 22 | III | 2,656,900 | 350 | 02:07/ 24:10 | 46:01/ 52:01 | 01:02/ 12:02 |
| 3193 | M | 24 | III | 2,412,840 | 289 | 33:03/ 33:03 | 44:03/ 58:01 | 03:02/ 07:01 |
| 8522 | M | 29 | III | 10,000,000 | 265 | 24:07/ 33:03 | 51:02/ 58:01 | 03:02/ 14:02 |

<sup>a</sup> Sex: M (male), F (female)<sup>b</sup> Fiebig stage was determined according to Fiebig *et al.* (2003)<sup>c</sup> Level of plasma HIV-1 RNA was determined by assays approved for clinical care<sup>d</sup> MHC-I HLA alleles were determined by Ehrenberg *et al.* (2014) and Ehrenberg *et al.* (2018)

### Determination of single vs. multiple TF

As part of the selection criteria, all participants were previously characterized by standard near full-length SGS (10–12 sequences, spanning ∼8.8kb from *gag* to *nef*) as having acquiring a single TF (n=10) or multiple TFs (n=5) (**Table 2**). Here, we asked whether sequencing thousands of plasma viral RNAs by uSGS would lead us to the same conclusions, hypothesizing that this much more sensitive assay might facilitate detection of previously unidentified minority founder viruses. For this approach, we targeted two ∼234bp segments (after primer trimming) in the RT coding region of *pol* and in the gp120/V3 loop region of *env*. In total, we obtained sequences where each viral genome had a unique barcode, from a median of 11,252 viral genomes in *pol* and a median of 10,329 viral genomes in *env* from each participant (**Table 2**). Aggregate phylogenetic analysis confirmed an absence of sequence mixing of variants across participant samples (single TF samples shown in **Figure S1**). The phylogenetic topology of all participant-gene datasets exhibited ‘star-like’ structures around one or more central nodes reflective of TF virus sequences, the numbers of which were used to assign single or multiple TF designations in this study (**Figure 1**; **Figure S2 & S3**).

**Figure 1.**
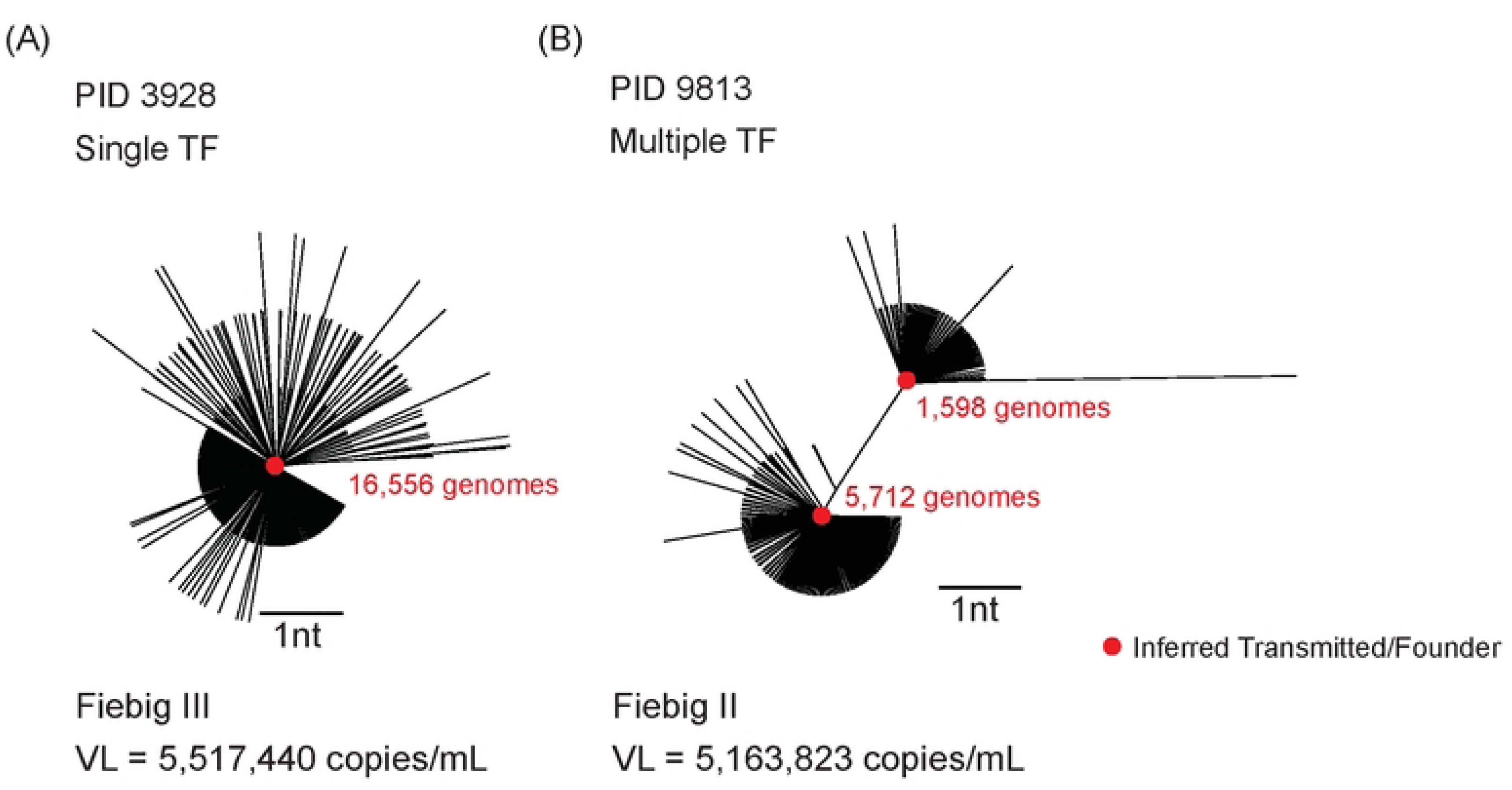
Single versus Multiple transmitted/founder phylogenies. (**A**) Example of an inferred single transmitted/founder (TF) virus. (**B**) Example of an inferred multiple TF virus. The red dots from which the diverging viral populations radiate indicate the inferred TF genome(s) with the number of viral genomes matching the TF found by uSGS shown in red. The Fiebig stage and level of plasma viremia (viral load, VL) of the donor are denoted below. The scale bar is set to 1 nt.

**Table 2.** Number of HIV-1 genomes sequenced, percent matching a transmitted founder, and number of possible transmitted founders for *pol* and *env*.

| PID | Fiebig Stage | # of TF by standard NFL-SGS<sup>a</sup> | *pol* |  |  | *env* |  |  |
| --- | --- | --- | --- | --- | --- | --- | --- | --- |
|  |  |  | # total genomes sequenced by uSGS<sup>b</sup><br>(# unique genomes) | % of total genomes matching TF<sup>c</sup> | # of TFs by uSGS | # total genomes sequenced by uSGS<sup>b</sup><br>(# unique genomes) | % of total genomes matching TF<sup>c</sup> | # of TFs by uSGS |
| 7279 | II | 2 | 11,252 (279) | 87, 8 | 2 | 9,611 (215) | 95 | 1 |
| 6340 | II | 3 | 10,326 (848) | 90, 6, 2 | 3 | 9,889 (338) | 74, 10, 2 | 3 |
| 7905 | IV | 3 | 20,924 (531) | 50, 17, 15, 10 | 4 | 10,211 (284) | 52, 32, 12 | 3 |
| 9813 | II | 2 | 7,835 (216) | 73, 20 | 2 | 8,868 (184) | 79, 17 | 2 |
| 7268 | III | 3 | 2,264 (139) | 71, 10, 6 | 3 | 1,929 (74) | 75, 17 | 2 |
| 3928 | III | 1 | 19,251 (472) | 86 | 1 | 17,242 (351) | 94 | 1 |
| 3832 | II | 1 | 12,518 (219) | 95 | 1 | 12,993 (215) | 96 | 1 |
| 5436 | III | 1 | 7,766 (206) | 91 | 1 | 10,329 (209) | 85 | 1 |
| 3698 | III | 1 | 8,148 (289) | 69, 18 | 1 or 2<sup>d</sup> | 10,120 (238) | 95 | 1 |
| 6609<sup>e</sup> | III | 1 | 10,386 (269) | 92 | 1 | 13,476 (228) | 91 | 1 |
| 8123 | III | 1 | 11,339 (238) | 94 | 1 | 13,867 (241) | 93 | 1 |
| 3513 | III | 1 | 12,913 (230) | 96 | 1 | 12,459 (209) | 92 | 1 |
| 9114 | III | 1 | 27,884 (438) | 87 | 1 | 10,957 (192) | 96 | 1 |
| 3193 | III | 1 | 3,704 (96) | 94 | 1 | 6,914 (144) | 96 | 1 |
| 8522 | III | 1 | 22,675 (351) | 94 | 1 | 12,602 (138) | 95 | 1 |

<sup>a</sup> Near Full-Length Single-Genome Sequencing (NFL-SGS), ~8.8kb from *gag* to *nef*; 10-12 genomes obtained from each sample (48)
<sup>b</sup> Ultrasensitive single-genome sequencing (uSGS)
<sup>c</sup> TF = Transmitted founder - exhibit a Poisson distribution of mutations and a radial (star-like) phylogeny (2); if otherwise and recombinants were evident between viral populations, was considered multiple TFs
<sup>d</sup> Two viral populations separated by a Hamming distance of 1 nt such that a classification could not be confidently assigned
<sup>e</sup> Based on initial near full-length sequencing (*gag* to *nef*) PID 6609 appears as a 'Single' TF, however, *env* sequencing of others found a co-infection with a highly divergent variant (48)

Using our uSGS method compared to the prior standard near full-length SGS analysis, concordant scoring of single or multiple TF was achieved in 14/15 participants (**Table 2**), illustrating that, in most cases, low-level sampling of much longer viral sequence segments during acute infection is sufficient for accurate TF identification. In the uSGS datasets, approximately 94% of sequences matched the previously reported TF sequences in both *pol* and *env*. Some minor deviations from the ‘star-like’ phylogenies were due to recombination among multiple TF lineages (**Figure S3**). Overall, these results demonstrate general concordance between TF designations made using standard near-full length SGS of 10-12 genomes and uSGS of thousands of genomes.

### Observed vs. expected mutation rates in acute HIV infection

To better understand the dynamics of early HIV sequence diversification, we used several analyses to assess the observed vs. expected number of mutations in the uSGS datasets: 1) A simple Monte Carlo mutation-only model, 2) Observed mutation counting, 3) Monte Carlo mutation accumulation with viral generation time sensitivity analysis, and 4) TF-match Poisson estimate (details in **Text S1**). First, we used a simple mutation-only model to establish a null expectation of mutations if diversity is driven mainly by HIV-1 RT error. Given the HIV-1 single-step replication error rate (*ca* 10^-5^ mutations/nt/cycle (*51*)), and assuming a replication rate of 1 cycle/day, we expected a mean of 70.2 mutations to occur each day in a region of 234bp in a population of 10,000 viral genomes (**Text S1 figure 1a**). We used Monte Carlo simulation to find the expected number of mutations over 30 days post-transmission following the HIV-1 RT single-step error corrected for *in vitro* assay error (*51, 52*) (**Text S1 figure 1b**). This approach allowed us to plot representative independent lineages with the expected Hamming distance over the first 30 days post-transmission. The accumulation of mutations should follow a Poisson distribution assuming mutations during this period are discrete and independent events (**Text S2 figure 1c**). Based on simple modeling, assuming no significant exertive selection pressures, the expected percent of viral genomes to have a Hamming distance of zero (*i.e.*, matching the inferred single TF) was 87% (Fiebig II, calculated as 19 days post-transmission) and 85% (Fiebig III, 23 days post-transmission). In contrast, among the participants in the study, we observed a median of 95% of *pol* and 93% of *env* (**Table 2**) sequences matching the inferred TF, suggesting strong purifying selection *in vivo*.

The second approach used counting observed mutations. As will be discussed in more detail later, a preponderance of G to A mutations in genomes with a Hamming distance ≥2 implied a significant influence of APOBEC3G/F on the observed diversity, leading us to analyze data in two separate ways: with and without the inclusion of potential APOBEC3G/F-induced (GR to AR) mutations. Data with all observed mutations are denoted as “unfiltered” (*i.e.*, including all GR>AR); whereas data with potential APOBEC3G/F-induced mutations removed (*i.e.*, excluding ≥2 GR- AR) are denoted as “filtered.” It is important to note that in the filtered data, a single GR-AR mutation, which may still have resulted from APOBEC3G/F, was allowed (discussed further in **Methods**). While this choice affected the total number of genomes analyzed, the difference was not significant with respect to the number of genomes with either zero or 1 nt difference to the TF.

In the unfiltered dataset, the inferred minimal single-step mutation rates in both *pol* and *env* were found to be significantly less than the minimal inferred HIV-1 RT single-step error (*51*), with *pol* having a median of 1.46x10^-5^ mut/site/day (2.05-fold lower than expected; *p*=0.0007, log_10_- transformed one-sample t-test) and *env* a median of 1.10x10^-5^ mut/site/day (2.73-fold lower; *p*<0.0001, log_10_-transformed one-sample t-test) (**Table S1**). The same was also true when accounting for GR-AR filtered viral genomes for *pol* and *env* (*p*=0.001 and *p*<0.0001, respectively; **Table S1**). We also compared these inferred rates to an *in vitro* tissue culture HIV-1 RT rate of 1.4x10^-5^ mut/site/cycle (*52*) in both *pol* and *env* for the unfiltered (*p*=0.26 and *p*=0.051, respectively) and GR-AR filtered datasets (*p*=0.89 and *p*=0.04, respectively) (**Table S1**).

Thus far, we have explored a simple theoretical baseline and empirical estimate based on observed mutations. However, the viral generation time may vary and the exact sampling date post-transmission is unknown. In order to better establish a null expected number of mutations under an HIV-1 RT error alone and test the effects of slower viral replication, we applied a third approach. While these data were initially derived with a viral generation time of 1 cycle per day, 1.5 to 2.0 days may be more consistent in biological systems (*51–54*). Despite the lengthened viral generation time, for unfiltered data, the observed mutations comprised only a median of ∼26– 54% (3.0x10^-5^ mut/site/day applied) or 40-75% (1.4x10^-5^ mut/site/day applied) of the number expected mutations (*p*=0.08, Wilcoxon signed-rank test) (**Text S1 tables 1-6**). Consequently, the data suggest a strong purifying selection, including at APOBEC3G/F-targeted sites (40 sites in *pol* vs. 24 sites in *env*), and/or the existence of sub-populations with different replication and mutation rates.

Finally, we used the proportion of unfiltered genomes matching the inferred TF to estimate a Poisson-informed mean (**Table S2**; details in **Text S1**). The median rates were 1.35x10^-5^ mut/site/day for *pol* (2.2-fold lower than the expected rate from HIV-1 RT; *p*=0.0009, log_10_- transformed one-sample t-test) and 1.05x10^-5^ mut/site/day for *env* (2.9-fold lower than expected from HIV-1 RT; *p*<0.0001). This difference between the HIV-1 RT Poisson-inferred and observed mutation-informed rates was consistent with previous analyses that suggest a strong purifying selection in early infection substantially constrains viral diversification. However, these rates were more similar when compared to the the *in vitro* tissue culture rate of 1.4x10^-5^ mut/site/day (*52*) in both *pol* and *env* (*p*=0.57 and *p*=0.28, respectively) (**Table S3**). Of note, in one participant (PID 3698) there was a potential ‘Founder Effect’ present (noted in **Table 2**) in which 18% of the sequences differed from the TF by a single C to T mutation from the TF (**Figure S4**). In examining these as two populations, using a TF-match Poisson estimate, two rates could be estimated at 3.47x10^-6^ mut/site/day and 2.95x10^-5^ mut/site/day, respectively.

### Accumulation of mutations after single TF follows a Poisson-like distribution

To visualize the accumulation of mutations following transmission of a single TF, the Hamming distances (HD) relative to inferred single TF viral sequences were calculated and plotted with unfiltered and GR-AR filtered data (**Figure 2**). Following transmission of a single HIV-1 variant, the initial accumulation of mutations due to replication error and in the absence of selection is expected to follow a Poisson distribution. We assessed this fit using both the χ^2^-goodness of fit test and Kolmogorov-Smirnov test for distribution comparison. Since the large χ^2^-statistic with small p-value is likely driven by the Law of Large Numbers, while the Kolmogorov-Smirnov test is comparatively less sensitive to large sample sizes, we deemed the latter test to be generally superior for this application. Our analysis showed that HIV-1 populations after a single variant transmission reflected a Poisson/Poisson-like distribution in both *pol* and *env* (further discussion in the **Methods**). However, we consistently observed Hamming distances of >3nt even though the probability of observing Hamming distances >3nt within 12–25 days post-transmission was far <10^-6^ in a simplified model (**Text S1 figure 1c**) (*2*).

**Figure 2.**
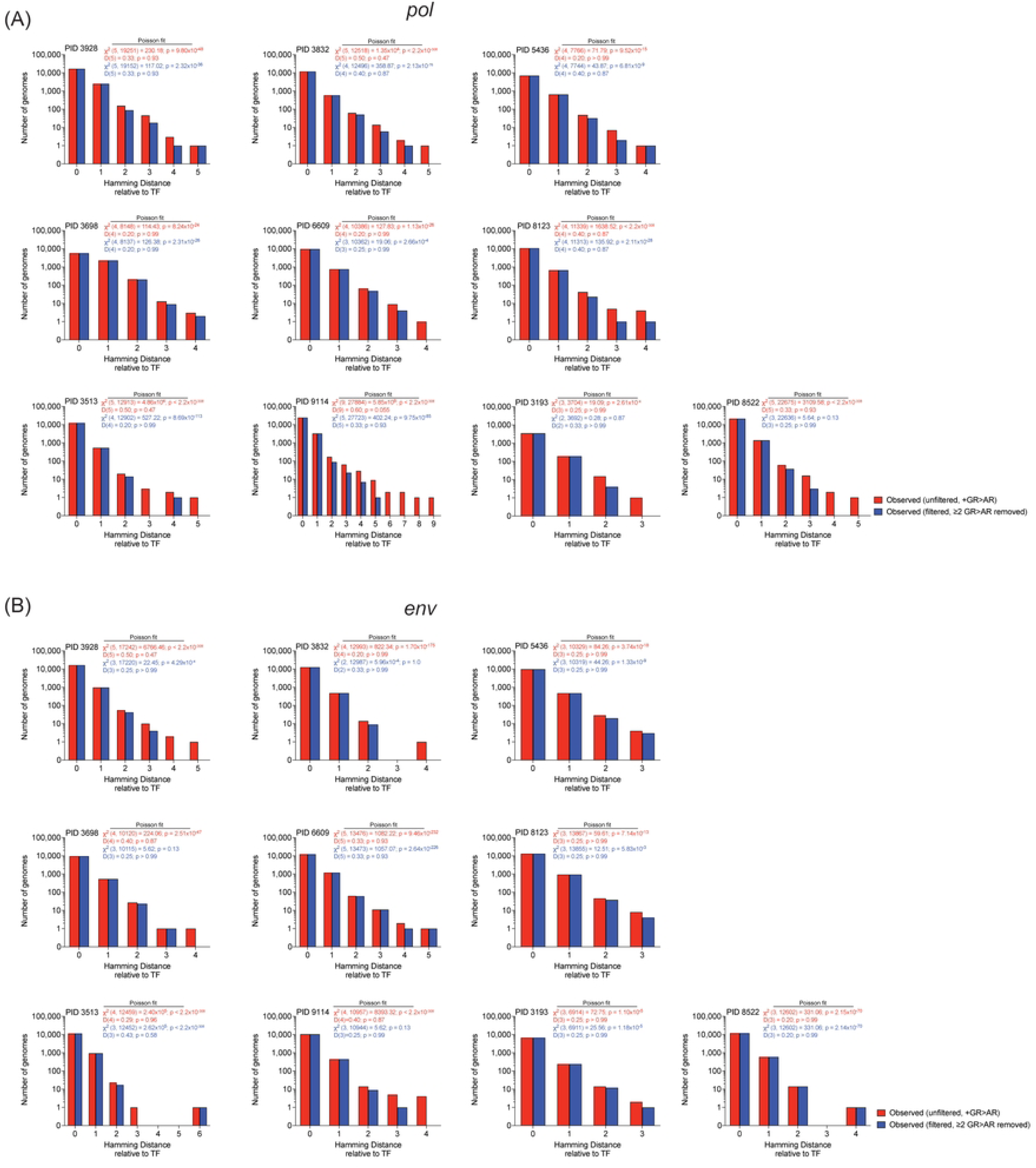
Hamming distance plots for inferred single TF virus participants. Hamming distances calculated relative to the respective inferred single TF virus are shown (**A**) for the *pol* region and (**B**) for the *env* region. Bars indicate the number of viral genomes sequenced in unfiltered (red) and GR-AR filtered (blue) datasets. The expected number of viral genomes was determined as a probability based on the Poisson fit of each respective observed dataset. The χ^2^-goodness of fit test and Kolmogorov-Smirnov test are reported for both the unfiltered and GR- AR filtered datasets.

A major potential driver of higher than expected Hamming distances (HD) relative to the TF is likely to be hypermutation due to APOBEC3G/F, which deaminates dC-(dC/dT) residues in nascent reverse transcripts. Consistent with a role for these mutagenic host factors, in most samples, sequences with the greatest HD were removed by GR>AR filtering (**Figure 2**). This analysis provided evidence that APOBEC3G/F is a major factor driving mutations in genomes that have a HD >3-4nt in acute HIV infection. An example can be seen in PID 9114 *pol* where filtering removed 16/17 viral genome sequences with a Hamming Distance >4nt. In a few cases (*e.g.*, PID 3193 *pol*), after filtering, the χ^2^-goodness of fit test was in agreement with the Kolmogorov-Smirnov test for the data following a Poisson distribution (**Figure 2**). Thus, APOBEC3G/F activity has a large effect on sequences with the furthest Hamming distance from the TF.

Finally, in merging our modeling analyses and the observed accumulation of mutations in a Poisson-like distribution, we turned to sequence-level viral evolution simulation. Here, we modeled expected Hamming distance distributions for comparison with observed viral populations using empirically-defined nucleotide substitution rates across replication cycles (**Text S1**). Primarily, we compared the observed GR-AR filtered datasets to the model using three rates: 3.0x10^-5^ mut/site/day (*51*), 1.4x10^-5^ mut/site/day (*52*), and the PID-specific inferred TF-match Poisson estimate rate (**Text S1 figure 2**). While anectdotal in nature, as the inferred TF-match

Poisson estimate rate was near or less than 1.4x10^-5^ mut/site/day, the two predictive models became indistinguishable to the observed data.

### Genetic characterization of descendants of single TF viruses

To further characterize the very early stages of HIV-1 evolution, genetic diversity in *pol* and *env* in those participants with inferred transmission of a single TF was compared using multiple measurements (**Table 3**). First, we measured the ratio of transitions versus transversions (Ti/Tv) and found, as expected, that transitions (*i.e.*, A<->G and C<->T) were strongly favored in both *pol* (Ti/Tv=5.1) and *env* (Ti/Tv=3.8). We examined biases towards specific nucleotide substitutions by analyzing rates of all possible mono-nucleotide combinations of mutations from the TF variants (**Figure 3A-D**; **Figure S4C**). When comparing mono-nucleotide exchange rate frequencies, consistent with the Ti/Tv ratio, transition mutations were highly prevalent in both genes. The mono-nucleotide substitution of G>A was most common (consistent with our results above showing the impact of APOBEC3G/F on HIV diversity in acute infection) greatly exceeding A>G, with C>T and T>C being second and approximately equal. Transversions were rare with G>T and T>G being the least frequent mutations in the datasets. Since both strands are initially synthesized by HIV-1 RT, one would expect for G>A vs. C>T and A>G vs. T>C rates to be respectively about equal. However, this was not the case when comparing the rate of G>A/C>T in *pol* (∼3.7) and *env* (∼3.0); yet A>G/T>C were closer to expected in *pol* (∼1.3) and *env* (∼1.4) (**Figure 3E-F**). While the overall bias toward transitions is consistent with properties of HIV-1 RT (*51, 55*), there was an unexpected skew with G>A/C>T. Despite the transition bias, the global transition/transversion ratio was not significantly different between the two subgenomic regions (*p*=0.24, Wilcoxon matched-pairs signed-rank test). The median average pairwise distances (APD) in *pol* (0.07%) and in *env* (0.05%) were similar for the unfiltered data and the GR-AR filtered data (0.06% in *pol*) and (0.05% in *env*) (*p*=0.12, paired Wilcoxon test). Finally, the mean ratio of non-synonymous to synonymous mutations (dN/dS) in the two genes, relative to the inferred TF, was different, 0.38 in *pol* (implying strong purifying selection) and 1.0 in *env* (indicating that an equal number of sites were under positive and negative pressure) (*p*=0.03, paired Wilcoxon test) (**Figure S5**). By contrast, when compared to the consensus CRF01_AE sequence, both *pol* and *env* were under purifying selection (**Table 3**; 0.12 for *pol* and 0.57 for *env*, *p*=0.002, paired Wilcoxon test). The dN/dS difference when comparing to the inferred TF vs. the consensus CRF01_AE was significant in *pol* but not in *env* (*p*=0.02 and *p*=0.25, respectively; paired Wilcoxon test). This difference was not appreciably affected with the differential effects of APOBEC3G/F on *pol* vs. *env* (40 targetable site in *pol* and only 24 in *env*).

**Figure 3.**
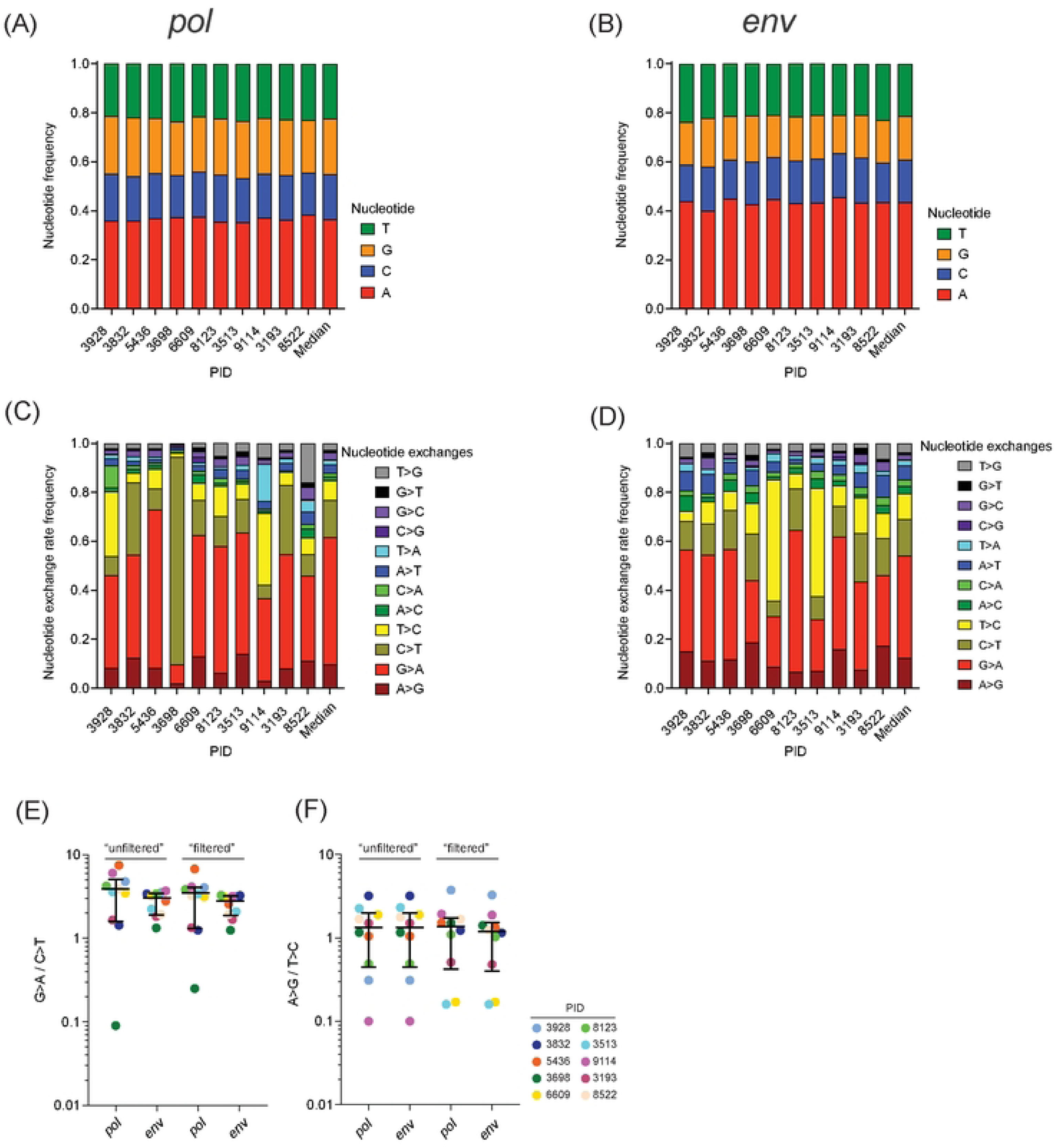
Nucleotide frequency and substitution rates of single TF. Nucleotide frequencies observed in (**A**) *pol* and (**B**) *env* across all viral genomes were compared to the correspondingTF. Median Individual nucleotide exchange rates relative to the single TF virus in *pol* (**C**) and *env* (**D**) are shown for each participant with a single TF. (**E**) The ratio of G>A/C>T mutations in *pol* and *env* for the unfiltered and GR-AR filtered datasets. Median and IQR are shown. (**F**) The ratio of A>G/T>C mutations in *pol* and *env* for the unfiltered and GR-AR filtered datasets. Median and interquartile ratio are shown.

**Table 3.**
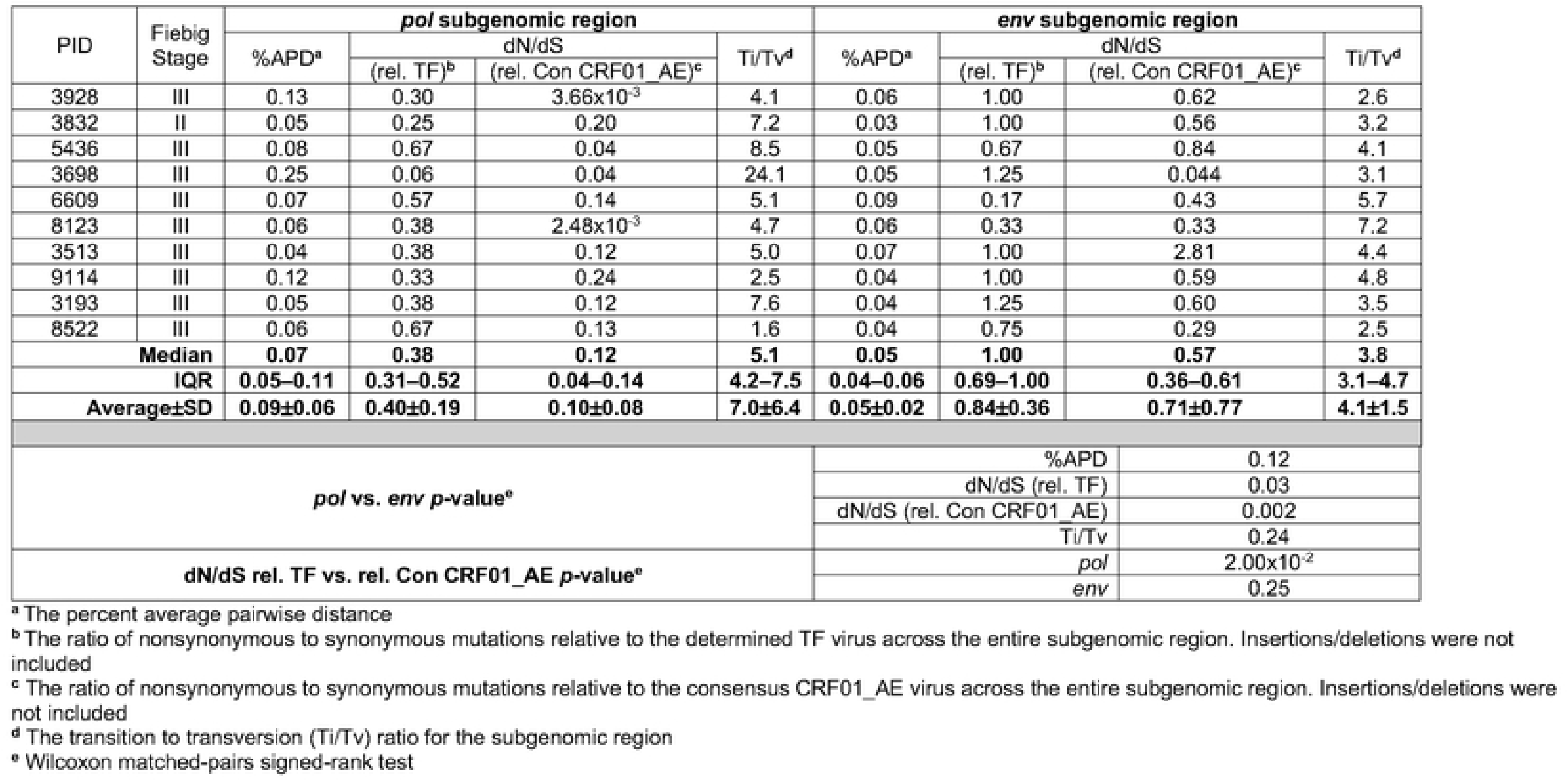
Genetic diversity after transmission of single founders for *pol* and *env* regions.

### Impact of APOBEC3G/F activity on HIV-1 diversity in acute infection

We asked what fraction of G>A mutations were likely to have been caused by APOBEC3G/F activity (using the dinucleotide change GR>AR) in the unfiltered data. For all observed G>A mutations, a median of 82.4% [IQR 74.9–87.3%] in *pol* and 78.1% [IQR 72.8–82.5%] in *env* could be attributed to APOBEC3G/F (**Figure 4A**; **Table S4**). Additional parsing found that the median contribution of APOBEC3F was greater than APOBEC3G by 3.4-fold for *pol* (60.4% vs 18.0% ; *p*=0.001, paired Wilcoxon test) and 4.1-fold for *env* (60.8% vs. 14.8%; *p*=0.003, paired Wilcoxon test). Next, we determined the fraction of all observed mutations that likely resulted from APOBEC3G/F. Across all observed mono-nucleotide mutations (not only G>A mutations), a median of 37.1% [IQR 29.5–41.6%] for *pol* and 26.9% [IQR 19.0–34.2%] for *env* could be attributed to APOBEC3G/F (**Figure 4B**). As expected from above, the median contributions of APOBEC3F were greater than those of APOBEC3G by 3.7-fold for *pol* (26.8% vs. 7.2%; *p*=0.002, paired Wilcoxon test) and 3.4-fold for *env* (18.3% vs. 5.4%), respectively. Thus, consistent with previous analyses, APOBEC3G/F-induced mutations contribute an important role to early HIV diversification.

**Figure 4.**
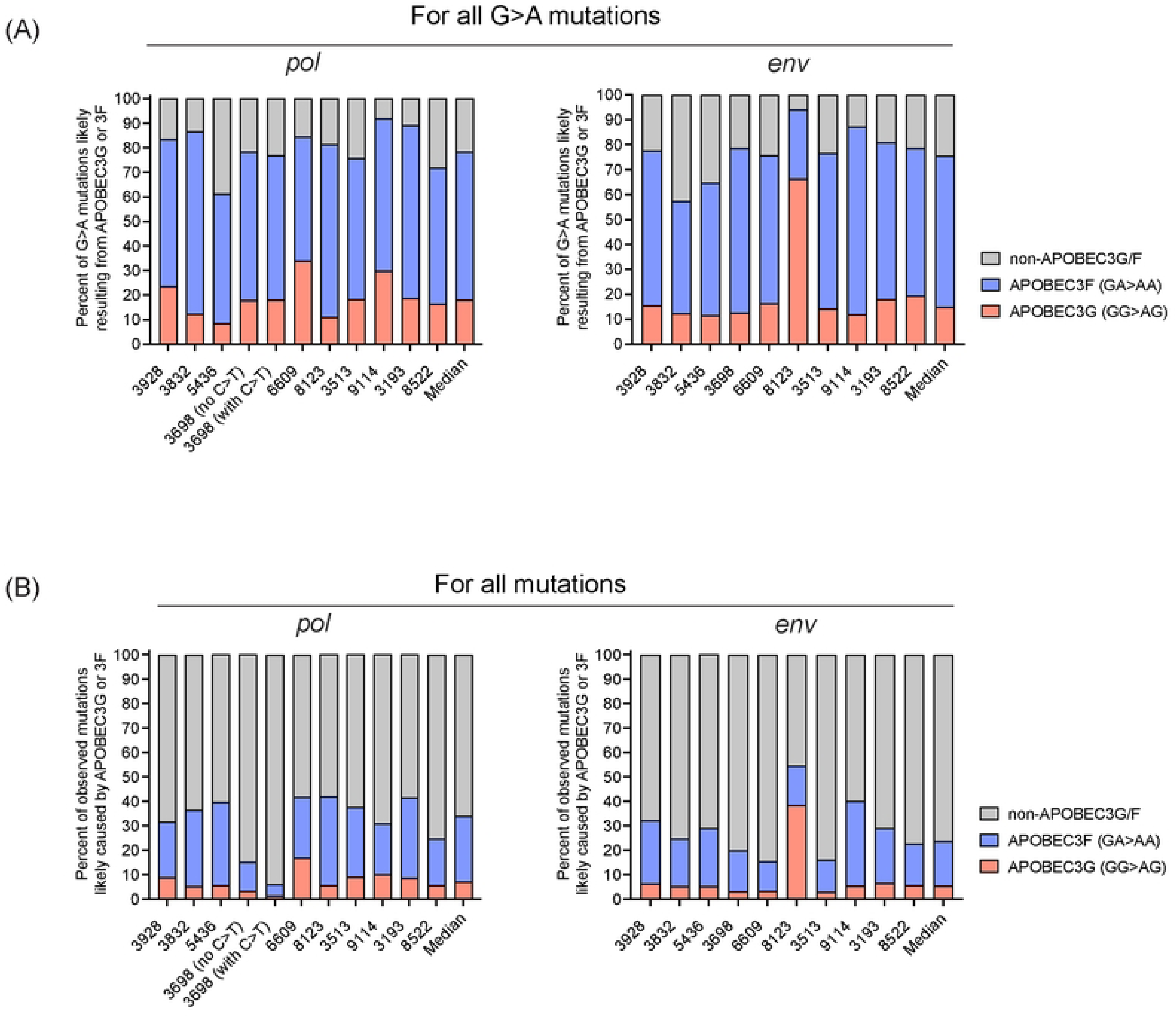
Contribution of observed APOBEC3G/F-mediated mutations among single TF participants. (**A**) For all observed mutations that resulted in a G>A transition relative to the TF, the fraction that could be attributed to APOBEC3G/F (GR>AR) by either APOBEC3G (GG>GA) or APOBEC3F (GA>AA) was determined for *pol* and *env*. The median across all participants is reported. Since PID 3698 *pol* has a potential ‘Founder Effect’ (a C>T mutation that likely occurred immediately after transmission), the data were analyed for both all viral genomes (denoted “with C>T”) and those with the C>T founder population removed (denoted “no C>T”). (**B**) For all observed mutations for a given participant relative to the TF, the fraction that could be attributed to APOBEC3G/F (GR>AR) by either APOBEC3G (GG>GA) or APOBEC3F (GA>AA) were determined for *pol* and *env*. The median across all participants is reported. Data are a representation of **Table S4**.

### Direct inference of early evolution during acute infection

Considering the significant effect of APOBEC3G/F on early HIV-1 diversification and the evidence for strong purifying selection overall, we asked if we could identify any mutations that appeared to have been positively selected in the first 30 days after transmission. To identify some possibilities, we reconstructed neighbor-joining trees of all sequences obtained from donors with single TF after GR-AR filtering (**Figure S6**) and without filtering (**Figure S7**). Of particular interest, artifacts originating from hypermutation due to APOBEC3G/F produce “pseudo-trees” that do not reflect proper phylogenetic relationships (*56*) (**Figure S8**). For example, in PID 9114 *pol*, there is a variant with a nonsynomous E270K_G>A_ mutation linked to nine other mutations (E292K_G>A_), (E292K_G>A_ + R316K_G>A_), and (E292K_G>A_ + R316K_G>A_ + G325R_G>A_) (**Figure S8A; >Figure S7** PID 9114 *pol* tree with red * at node), all of which are GR>AR changes. In another example, for PID 3928 *env*, there is a variant with a nonsynonymous G324R_G>A_ mutation linked to seven other mutations: (R327K_G>A_), (R327K_G>A_ + R324K_G>A_ + G319E_G>A_), (G301R_G>A_), and (G310R_G>A_ + R327K_G>A_ + Q305Q_G>A_ + V254V_G>A_) (**Figure S8B; >Figure S7** PID3928 env tree with blue * at node). All of these mutations were GR>AR changes, however, two were synonymous. The presence of pseudo-topologies were pervasive in all participants for both *pol* and *env* subgenomic regions. An illustrious example can be seen in PID 3193 *pol* which contains complex pseudo-topology resulting in multiple nodes (**Figure S9A**). After filtering out singlet leaves and branches (**Figure S9B**), we further masked potential APOBEC3G/F sites (**Figure S9C**). The majority (50/55 viral genomes or 16 unique genomes) collapsed identically into the TF; thus demonstrating that all 20 GR positions directly contributed to the topologic differentiation of the pseudo-tree. Further, within the pseudo-tree, an internal branch (5/55 or 3 unique genomes) deconvoluted as a single collapsed leaf/branch (**Figure S9C**). Interestingly, only one population (bright blue branch) maintained evidence of independent mutation accumulation (**Figure S9**). Thus, in subtrees with extensive G>A changes, haplotypes that might appear to arise from sequential accumulation of mutations are more likely artifacts resulting from APOBEC3G/F activity occurring during the same replication cycle, rather than independently accumulating during successive cycles.

Consequently, we focused on phylogenetic relationships within the GR-AR filtered data, and found evidence for what could be selection for rare mutations. For example, in PID 3928 *pol*, we observed mutations in RT on a backbone with a non-synonymous change (S252R_C>A_) linked to 3 different lineages (T216T_T>C_, E194D_A>C_, and Y188H_T>C_ + M231I_G>A_) (**Figure 5A**). It is possible that the M231I mutation arose from APOBEC3G/F activity since the filtering permitted the allowance of 1 GR>AR mutation. In PID 6609 *pol*, we observed linkage of three mutations (E194E_A>G_ + Q197E_C>G_ + L209P_T>C_). Similar patterns were observed in *env* (**Figure 5B**). In the selection of representative sub-trees, there were several key branching structures that were observed which we aimed to highlight here. The observation that for the most part, the maximum number of mutations occurring on a single genetic backbone was 3nt, which is consistent with what would be expected based on the Poisson distribution for HIV-1 RT replication error given the amount of time post-transmission. Although rare cases of linked mutations were seen, overall, the filtered data yielded very little evidence for positive selection in acute HIV-1 infection.

**Figure 5.**
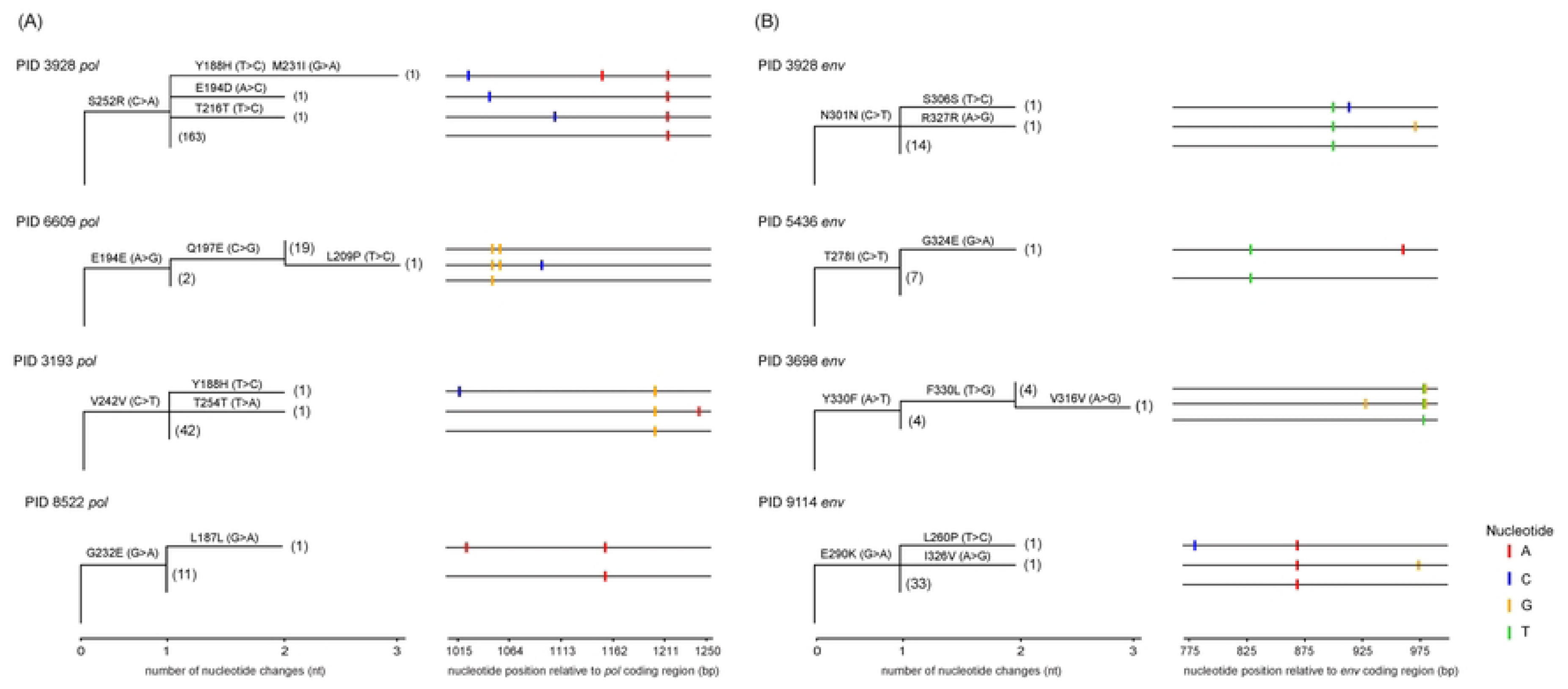
Phylogenetic reconstruction of early evolution during Fiebig stages II-III. Neighbor-joining phylogenetic trees were constructed for filtered and unfiltered sequence data from inferred single TF virus participants for (**A**) *pol* and (**B**) *env* and selected branches that represented the most common topologies were excised for further analysis The excised branches are plotted according to the Hamming distance rooted on that participant’s inferred single TF sequence. Labeled on each branch is the corresponding amino acid changes with the associated nucleotide change. The leaves of the trees show the number of viral genomes (in parentheses). In alignment to the phylogenies, an SNP matrix (highlighter plot) is provided where changes to the inferred single TF virus are noted as: A (red), G (orange), C (blue), and T (green). The nucleotide position is relative to the coding region sequenced in HXB2 coordinates. The listed mutations on the branches are relative to the amino acid change in HXB2 coordinates.

### Only a small fraction of mutations are reversions to the consensus CRF01_AE in acute infection

Based on our earlier studies (*57*), we hypothesized that a significant fraction of the amino acid changes observed would be reversions from the TF back to the consensus CRF01_AE. However, in *pol*, we found very few positions in the TF viruses that were different than the consensus (median of 2 positions per PID) and, surprisingly, only 0.33% of these amino acid mutations in the datasets were reversions to the consensus CRF01_AE (**Table 4**). However, in *env*, there were more positions in the TF that were divergent from the consensus CRF01_AE (median 8 positions per participant) (*p*=0.002, paired Wilcoxon test), possibly reflecting antibody escape in the individual who transmittrd the virus to the current donor, whereas 2.8% were reversions to the consensus CRF01_AE (**Table 4**; *p*=0.008, paired Wilcoxon test). These data show that reversions to the consensus CRF01_AE are not likely to be a significant driver of HIV-1 diversity in acute infection.

**Table 4.**
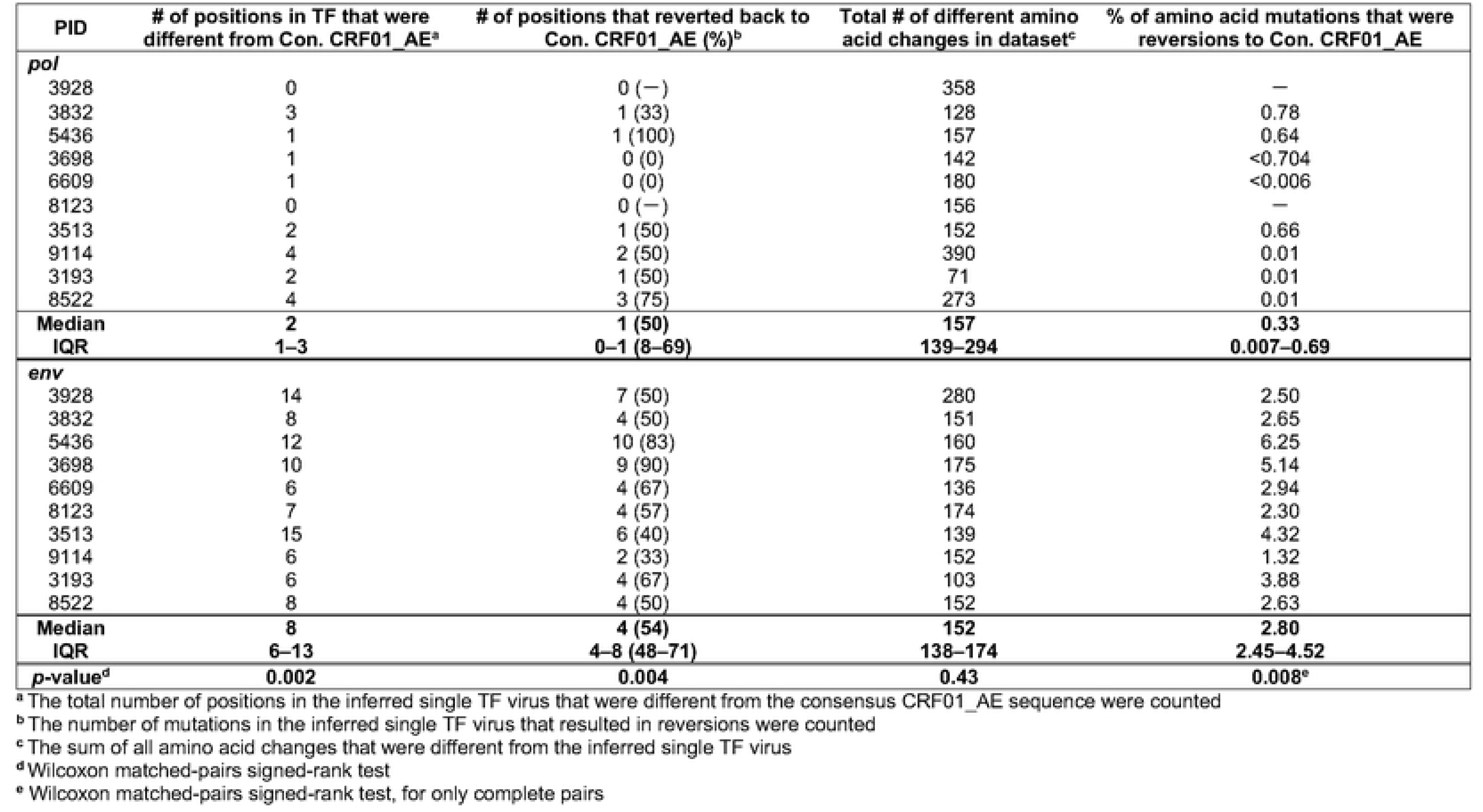
Percent of positions that are reversions from TF to the consensus CRF01_AE Table 5. Percent of mutations that mapped to CTL epitopes.

| PID | # of positions in TF that were different from Con. CRF01_AE<sup>a</sup> | # of positions that reverted back to Con. CRF01_AE (%)<sup>b</sup> | Total # of different amino acid changes in dataset<sup>c</sup> | % of amino acid mutations that were reversions to Con. CRF01_AE |
| --- | --- | --- | --- | --- |
| *pol* |  |  |  |  |
| 3928 | 0 | 0 (—) | 358 | — |
| 3832 | 3 | 1 (33) | 128 | 0.78 |
| 5436 | 1 | 1 (100) | 157 | 0.64 |
| 3698 | 1 | 0 (0) | 142 | <0.704 |
| 6609 | 1 | 0 (0) | 180 | <0.006 |
| 8123 | 0 | 0 (—) | 156 | — |
| 3513 | 2 | 1 (50) | 152 | 0.66 |
| 9114 | 4 | 2 (50) | 390 | 0.01 |
| 3193 | 2 | 1 (50) | 71 | 0.01 |
| 8522 | 4 | 3 (75) | 273 | 0.01 |
| **Median** | **2** | **1 (50)** | **157** | **0.33** |
| **IQR** | **1–3** | **0–1 (8–69)** | **139–294** | **0.007–0.69** |
| *env* |  |  |  |  |
| 3928 | 14 | 7 (50) | 280 | 2.50 |
| 3832 | 8 | 4 (50) | 151 | 2.65 |
| 5436 | 12 | 10 (83) | 160 | 6.25 |
| 3698 | 10 | 9 (90) | 175 | 5.14 |
| 6609 | 6 | 4 (67) | 136 | 2.94 |
| 8123 | 7 | 4 (57) | 174 | 2.30 |
| 3513 | 15 | 6 (40) | 139 | 4.32 |
| 9114 | 6 | 2 (33) | 152 | 1.32 |
| 3193 | 6 | 4 (67) | 103 | 3.88 |
| 8522 | 8 | 4 (50) | 152 | 2.63 |
| **Median** | **8** | **4 (54)** | **152** | **2.80** |
| **IQR** | **6–13** | **4–8 (48–71)** | **138–174** | **2.45–4.52** |
| **p-value<sup>d</sup>** | **0.002** | **0.004** | **0.43** | **0.008<sup>e</sup>** |

<sup>a</sup> The total number of positions in the inferred single TF virus that were different from the consensus CRF01\_AE sequence were counted<sup>b</sup> The number of mutations in the inferred single TF virus that resulted in reversions were counted<sup>c</sup> The sum of all amino acid changes that were different from the inferred single TF virus<sup>d</sup> Wilcoxon matched-pairs signed-rank test<sup>e</sup> Wilcoxon matched-pairs signed-rank test, for only complete pairs

### Nearly half of all amino acid changes fall within CTL epitopes

HIV replicating *in vivo* is subject to strong negative selection by the host immune response. To assess the extent of possible early escape from CTL pressure, we mapped epitopes matching each participant’s MHC-I HLA alleles to identify mutations that fell within CTL epitopes. While most CTL epitopes were initially identified in studies performed on HIV-1 subtype B, we were able to identifiy their genomic location in analogous regions in CRF01_AE (**Figure 6** & **Figure S10**). We found no significant differences in the total number of amino acid changes in *pol* (median 157 mutations) and *env* sites (median 152 mutations) (**Table 5**; *p*=0.43, paired Wilcoxon test). The difference was also insignificant when we calculated the number of amino acid changes that fell within CTL epitopes (compared to the consensus CRF01_AE), a median of 41.8% in *pol* and 51.3% in *env* (**Table 5**; *p*=0.56, paired Wilcoxon test). While it is known that some mutations in CTL epitopes eventually undergo selection in the early months of infection (*57*), we did not observe direct evidence of such selection in this acute HIV-1 infection cohort.

**Table 5.** Percent of mutations that mapped to CTL epitopes.

| **PID** | **Total # of different amino acid changes in dataset** | **Number of amino acid changes in CTL epitopes from Con. CRF01_AE** | **% of all amino acid changes in the dataset fall within CTL epitope** |
| --- | --- | --- | --- |
| *pol* |  |  |  |
| 3928 | 358 | 65 | 18.2 |
| 3832 | 128 | 83 | 64.8 |
| 5436 | 157 | 63 | 40.1 |
| 3698 | 142 | 95 | 66.9 |
| 6609 | 180 | 49 | 27.2 |
| 8123 | 156 | 77 | 49.4 |
| 3513 | 152 | 66 | 43.4 |
| 9114 | 390 | 76 | 19.5 |
| 3193 | 71 | 47 | 66.2 |
| 8522 | 273 | 54 | 19.8 |
| **Median** | **157** | **66** | **41.8** |
| **IQR** | **139–294** | **53–79** | **19.7–65.2** |
| *env* |  |  |  |
| 3928 | 280 | 59 | 21.1 |
| 3832 | 151 | 82 | 54.3 |
| 5436 | 160 | 54 | 33.8 |
| 3698 | 175 | 105 | 60.0 |
| 6609 | 136 | 81 | 59.6 |
| 8123 | 174 | 102 | 58.6 |
| 3513 | 139 | 67 | 48.2 |
| 9114 | 152 | 91 | 59.9 |
| 3193 | 103 | 42 | 40.8 |
| 8522 | 152 | 66 | 43.4 |
| **Median** | **152** | **74** | **51.3** |
| **IQR** | **138–174** | **58–94** | **39.1–59.7** |
| ***p*-value<sup>a</sup>** | **0.43** | **0.17** | **0.56** |

<sup>a</sup> Wilcoxon matched-pairs signed-rank test

**Figure 6.**
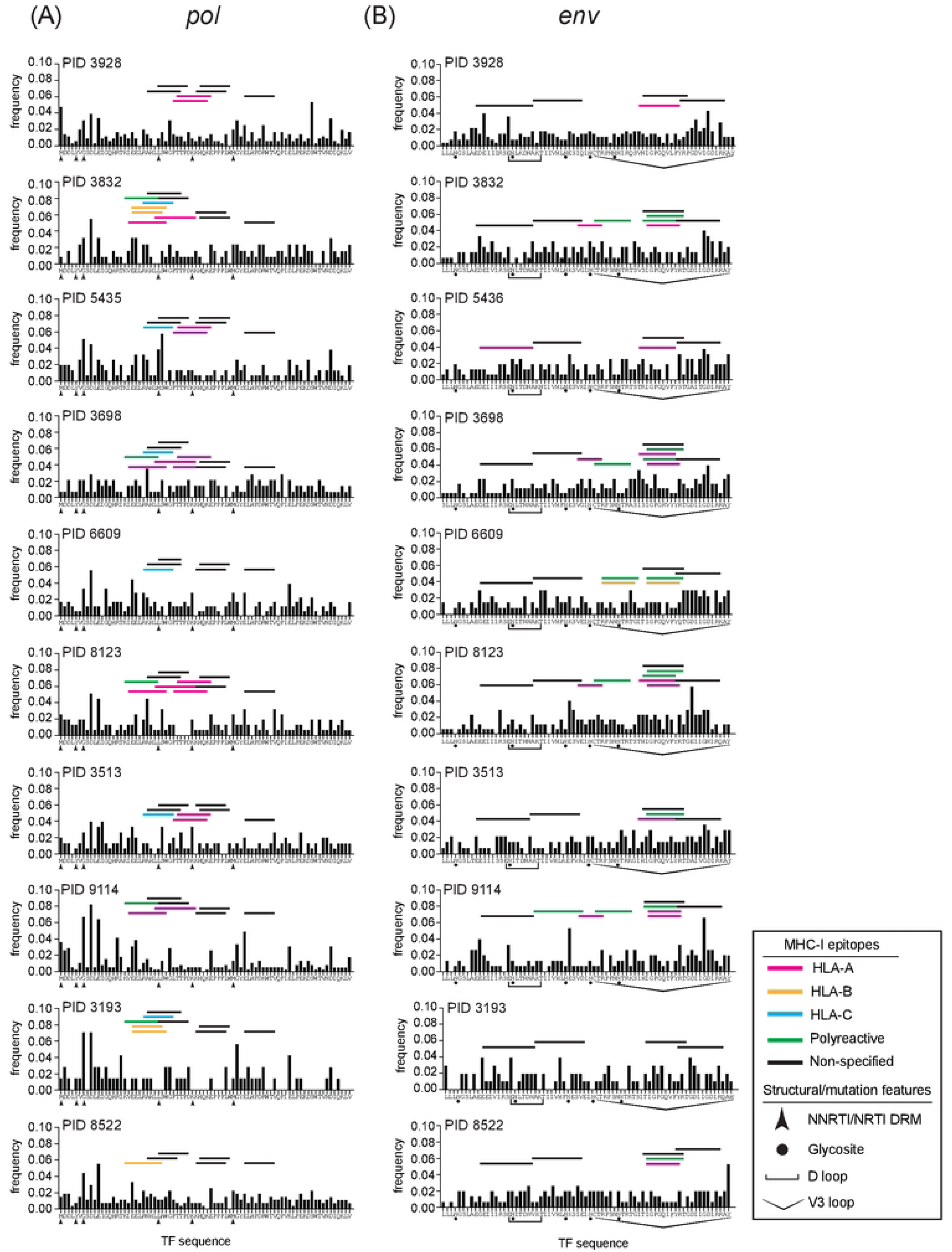
Frequencies of amino acid changes in *pol* and *env* compared to TF. Percent of amino acid changes across the sequenced genomic regions in (**A**) *pol* and (**B**) *env.* The participant identifier is noted next to each distribution. CTL epitope mapping is included based on the participant’s respective MHC-I class HLA alleles. Epitopes are mapped to within a region of 14 amino acids or less. Epitopes were designated as either specific to HLA alleles (*e.g.*, A*02; A* pink/ B* orange/ C* blue bars), polyreactive (*e.g.*, A*02/B*58; green bars), or when no MHC presenting molecule is defined the host species is note (*e.g.*, human; black bars). Structural and/or mutation features are denoted underneath each sequence: NNRTI or NRTI DRM, glycosite, D loop for CD4 binding, and V3 loop.

### Other potential drivers of genetic diversity and early evolution

We asked if viral coreceptor tropism might have influenced early HIV diversification by examining the V3 loop of sequences in the uSGS dataset (**Table S5**). To predict viral coreceptor tropism, we aligned each participant’s inferred TF V3 loop sequence (HXB2 coordinate codons 296 to 331) to a consensus CRF01_AE reference and extracted biologically informed features (*e.g.*, net charge, glycosylation motif, position 25, and X4-associated markers; further detailed in the **Text S2**) (*58–61*). We applied a machine learning model to predict coreceptor tropism of founder V3 loop sequences from 10 single TF designates and found that 9 were predicted to be R5- and 1 X4- tropic (**Text S2; Text S2 table 1**). In the intra-participant viral population (including variants that did not match the TF), most participant samples contained predominantly R5-tropic variants (≥90%), though low-frequency X4 subpopulations (8–12% of unique viral genomes or 0.43–0.90% total viral population) were genetically predicted to be detected in 3 participants (PIDs 3832, 8123, and 3513), possibly reflecting early transitional or dual-tropic states (**Table S5**). In contrast, PID 9114 exhibited a predominant predicted X4-tropic population (∼92% of unique viral genomes or 99.7% of the total viral population). Net charge analysis of the V3 loop further supported these predictions with samples harboring predominantly R5 variants having unimodal or bimodal distributions centered around +3 and +4 (mean ≈ +3.5, SD ≈ 0.6), while PID 9114 showed a broader, right-skewed distribution (mean ≈ +4.4) with many viral variants having a +5 or +6 V3 loop charge. Very low frequency mutations were observed at nearly every position in the V3 loop, whereas in PIDs 3698 and 3513, no sites were absolutely conserved (**Figure S11**). The 18^th^ and/or 21^st^ positions in V3 were conserved in 5 of 10 participants. The GPGR V3 apex motif was identified for only PID 3698 whereas 9 of 10 TF had a GPGQ motif which previously has been shown to induce broader cross-neutralizing anti-V3 antibodies compared to a GPGR V3 apex (*62*).

Finally, we looked for drug resistance mutations (DRMs) in the sequenced segment of *pol* that may have arisen in the absence of ART within the first 17–24 days post-transmission. We detected several nucleoside reverse-transcriptase inhibitor (NRTI) and non-nucleoside reverse-transcriptase inhibitor (NNRTI) resistance mutations (**Figure 7**; **Table S6**). Overall drug resistance mutation frequencies ranged from 0.21% to 4.03%. The two most common mutations observed were M184I (range: 0.35%–4.03%) and G190E (range 0.69%–3.65%). Both mutations (M184I and G190E) can result by GR>AR mutations, largely or entirely the result of APOBEC3G/F activity. This is especially likely in the case of 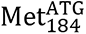 which is part of the YMDD active site motif (in the “palm” subdomain) of HIV-1 RT, and is highly conserved across HIV-1 Group M, where, immediately following, is 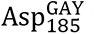 which establishes the GG motif for APOBEC3G targeting.

**Figure 7.**
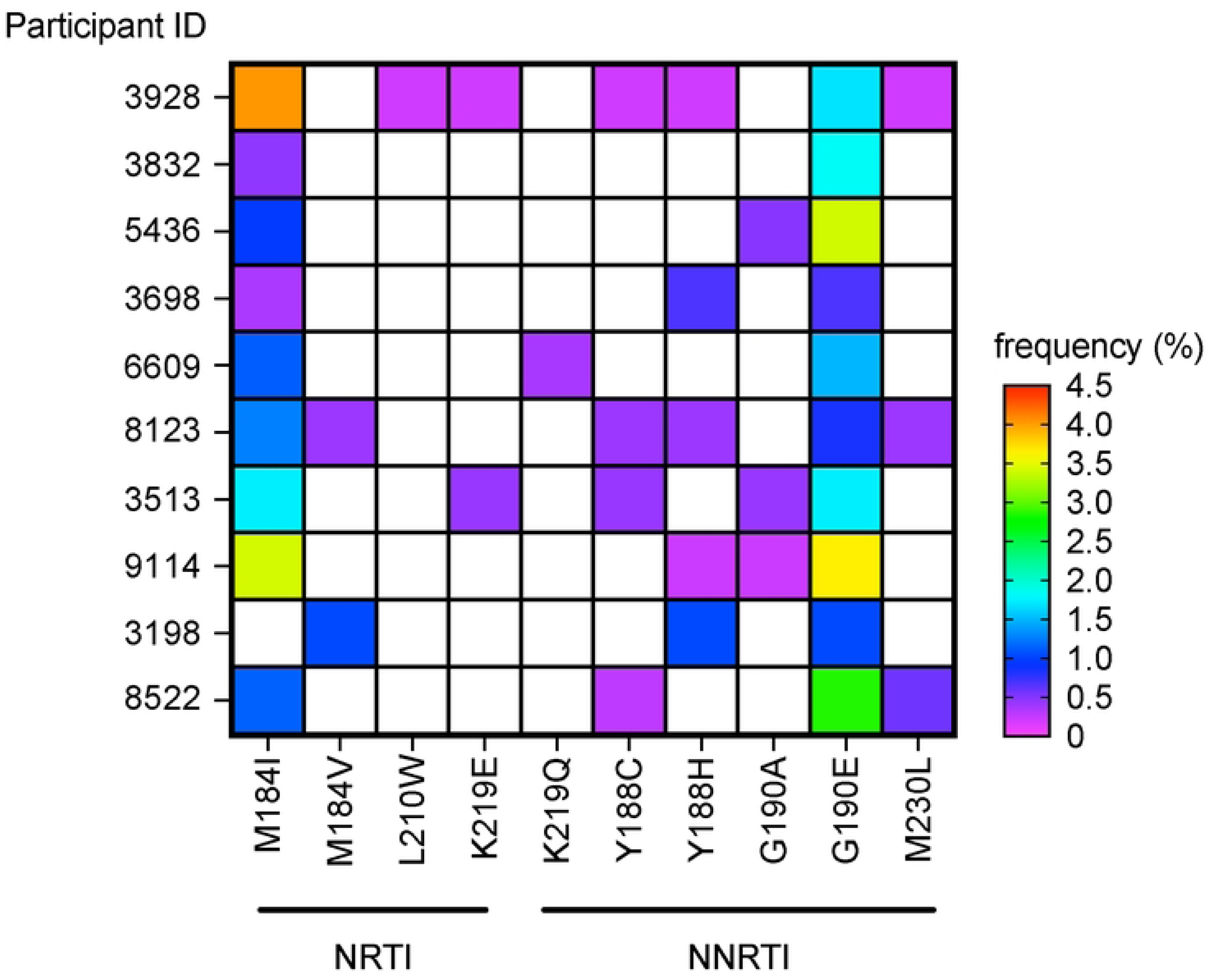
Drug resistance mutations in inferred single TF virus participants. The frequency of each predicted NRTI or NNRTI resistance mutation was determined based on the number of viral genomes in each PID that contained it. The frequency of each mutation is indicated by color gradient. Based on composite data shown in **Table S6**.

## DISCUSSION

In this study, we used uSGS to examine the earliest diversification of HIV-1 during acute infection. Our findings indicate that, although low-frequency variants emerge soon after transmission, they are observed at only a fraction of what is expected from previously reported HIV-1 RT error rates (*51, 52*), and a significant fraction of these are likely the result of APOBEC3G/F activity, and not HIV-1 RT error. By sequencing subgenomic regions of over 10,000 virions in each of 15 donors, we were able to observe early genetic variation at a level of resolution not previously reported, to our knowledge. Analysis of these large sequence datasets consistently revealed ‘star-like’ phylogenies characterized by numerous minor variants centered around one or more central nodes representing the likely TF. With one possible exception, the nodal sequences perfectly matched previous single vs. multiple TF classifications obtained from a few sequences of much larger segments of the viral genome (*40, 48*). Collectively, our observations demonstrate that both stochastic HIV-1 RT misincorporation error and APOBEC3G/F-mediated mutations contribute to early viral diversification, perhaps restricted by purifying selection, in acute infection. We show that mutation accumulation during acute infection is not primarily driven by reversion to the subtype consensus or by very early CTL selection, and we highlight a few possible mutations that may have undergone positive selection driving the earliest stages of viral evolution. These results provide new insights into the early stages of HIV-1 evolution and underscore the role of intrinsic mutation processes, offering a more nuanced understanding of viral diversification in the absence of significant immune or drug pressure.

Our deep dive into the genetics of acute infection showed that more than 90% of the plasma viral genomes matched the TF(s), with most variants differing from the TF(s) by one nucleotide in the subgenomic regions sequenced. Hamming distances among variants in datasets classified as being derived from single TF mostly followed a Poisson distribution for distances <3 nt. Observed mutation frequencies were only about one-third of what was expected from published estimates of HIV-1 RT mutation rates (*51, 52*), estimated *in vivo* replication rate (*63–66*), and time since transmission estimated by Fiebig stage (*40, 67*), suggesting that more than half of mutations that occur in each round of viral replication in acute infection are not sustained in the viral population. Alternatively, it is possible that the *in vivo* replication rate assumed from *in vitro* analyses do not accurately reflect the true biological system. Adjusting viral generation times from 1.0 day to 1.5 or 2.0 days (*53, 54*) did not change the outcome significantly, raising the fraction of observed vs. expected to ∼53-76%, all of which is consistent with early virus populations being shaped by strong purifying selection.

In a complementary analysis, we estimated a Poisson-informed mutation rate that was 2- fold lower than expected compared to previously reported rates (*51, 52*). This finding highlighted the natural variability in direct measurements, where stochastic mutation accumulation may lead to the underestimation of an estimated single-round error rate. While the Poisson fit provided a more refined (*i.e.*, informed) estimate, the adjusted rate still failed to fully account for the observed mutation patterns, suggesting that additional biological factors beyond HIV-1 RT, such as host antiviral factors (*e.g.*, APBOEC3G/F), purifying selection, viral fitness, and potentially immune pressures occurring less than 30 days post-transmission, may constrain early viral evolution. Our findings suggest that the accumulation of mutations in HIV-1 populations after the transmission of a single TF are largely stochastic, independent mutational events which have been observed by others (*2, 9, 68*), but also underscore the role of APOBEC3G/F among observed deviations. The calculation of Hamming distances relative to the inferred TF sequence provided a quantitative framework for assessing early viral diversification. This analysis supported the notion that, early post-transmission, HIV-1 evolution is mostly shaped by both the Poisson distributed accumulation of mutations by HIV-1 RT and APOBEC3G/F-mediated mutations, rather than by selection for specific mutations.

The prevalence of apparent APOPEC3G/F mutation deserves particular attention in these analyses, since it violates the assumptions that underlie the application of Poisson statistics. APOBECs are incorporated during virion assembly and act on newly synthesized minus-strand viral DNA to deaminate cytosine residues, converting them to uracil, leading to G>A mutation in the daughter genomes. Unlike HIV-1 RT error, the frequency of mutation is strongly context-dependent, with APOBEC3G preferring CC and APOBEC3F TC, with additional contribution from other nearby residues (*69–71*). Rather than occurring independently, the mutations are also strongly clustered, in violation of the Poisson requirement of independence and implying that multiple mutations occur during the same replication cycle. The result of this process is the independent generation of genome haplotypes containing multiple related mutations whose pattern is determined by their relative APOBEC3G/F target frequency, rather than by accumulation of independent mutations during sequential replication. The subsequent effects can be visualized as pseudo-trees with otherwise indistinguishable topology from trees arising by RT-generated mutations only. In these cases, large sub-trees at common nodes within a pseudo-tree possess sequences that all contain GR>AR mutations. A good example of this artifact is the *pol* tree from PID 9114 which shows a strong predominance of nonsynonymous mutations, often resulting in a different charged residue (particularly glutamic acid to lysine). It is highly unlikely that such mutations would be well-tolerated in reverse transcriptase.

We also characterized the nature of mutations in acute infection and found that nucleotide substitutions relative to the TF in all donors were primarily transitions, reflective of the tendencies of both HIV-1 RT and APOBEC3G/F. As expected, G>A mutations were the most frequent, followed by C>T and T>C, consistent with HIV-1 RT error and additional effects by APOBEC3G/F-mediated mutation (*51, 72–75*). These findings reveal that HIV-1 diversification in acute infection is shaped by intrinsic HIV-1 RT error under a Poisson distribution and nonuniformly distributed APOBEC3G/F-mediated mutations. With respect to APOBEC3G/F, Armitage and colleagues previously showed that during early infection, the majority of G>A changes (when including and excluding hypermutated sequences), were likely caused by APOBEC3F and considered ‘sublethal’ (*i.e.*, non-lethal in terms of not causing stop codons) (*76, 77*). Additionally, *in vitro* studies have recovered sequences from replication-competent viruses with an accumulation of ‘sublethal’ mutations caused by APOBEC3G/F (*78*).

In the setting of a limited early selection, one important potential factor in shaping acute infection is viral tropism for coreceptor usage. To assess whether early viral diversification contributed to coreceptor tropism, we analyzed the V3 loop of the TF viruses using a CRF01_AE subtype-specific XGBoost model trained on biologically informed features (*e.g.*, net charge, glycosylation motif, and other previously described X4-tropic features). This approach predicted 9 of 10 TF viruses in our study as R5-tropic, and one as X4-tropic. Minor X4-tropic variants detected in some R5 TF participants suggest early transitional or dual-tropic potential within the viral population. X4-tropic variants may be under-detected due to compartmentalization in naïve or central memory CD4+ T cells, which may contribute less to viremia (*79–84*). CRF01_AE isolates have been linked to higher CXCR4 usage, faster CD4^+^ T cell decline, and accelerated immune damage via productive or abortive infection and pyroptosis (*79, 85, 86*). The appearance of X4-tropic variants as early as Fiebig stage III suggests that phenotype divergence can occur remarkably early, although conditions favoring their selection may not occur until much later stages of disease progression. Together, our findings highlight the clinical and evolutionary importance of monitoring early tropism shifts, even when X4-tropic variants remain subdominant in plasma.

Further, we explored the potential emergence of *de novo* drug resistance mutations (DRMs) within the first 17–24 days post-transmission. Since all inferred TF viruses were wild-type, any observed DRMs were likely attributable to HIV-1 RT error or APOBEC3G/F activity. Despite the absence of selective drug pressure, we detected several NRTI and NNRTI resistance mutations, with M184I/V and G190A/E being the most prevalent (0.21%–4.03%). These data are consistent with previous work demonstrating low levels of DRMs despite the lack of ART experience (*87–89*) as well as the possibility of transmission of minor variants with DRMs (*90–94*). There was an initial indication that some rare mutations may have undergone positive selection in acute infection where the Hamming distances were greater than 3nt. However, most of these variants contained probable APOBEC3G/F-induced mutations and were omitted from the APOBEC3-filtered datasets. However, ocassional mutations were reversions back to the consensus CRF01_AE, which may indicate a viral fitness benefit. In general, the TF were largely similar to consensus, particularly in *pol*, where only a handful of positions differed. Even in *env*, where greater divergence was observed, reversions remained infrequent, suggesting that they do not represent a dominant pathway of very early evolution. We also found no significant enrichment of mutations within mapped epitopes during acute infection. Amino acid changes were equally likely to occur inside or outside predicted CTL targets, suggesting that T cell mediated immune-driven escape had not yet exerted a dominant or perceivable effect. This finding does not mean that selection was entirely absent as some mutations did fall within known epitopes; but overall, mutations appeared to be emerging in a largely neutral manner.

The frequency of amino acid changes further supported this notion. While mutations spanned nearly every position in *pol* and *env*, they remained at low frequencies, with most falling below 0.04% and only a few positions reaching higher levels (∼0.1%) perhaps entirely due to APOBEC3G/F. In some cases, regions with multiple overlapping CTL epitopes showed greater mutational activity, but in others, peaks emerged outside of known immune targets, further reinforcing the idea that T cell immune selection had not yet become a primary force shaping viral evolution. Furthermore, shared rapid diversification early in infection at specific codons under purifying selection across multiple PWH highlights the repeated selection on these sites during host adaptation; whereby the possibility that APOBEC3G/F-mediated mutations in CTL epitopes may help lay a foundational pathway for potential future immune escape (*95*). Though this may not be universally true in cases of multiple APOBEC3G/F targeted sites. Thus, while early mutations appeared to follow a stochastic pattern, with occasional ‘Founder Effects’ or selection pressures consistent with intrinsic viral mutation processes rather than widespread immune adaptive pressures, as with DRM, they also serve as escape mutations preexisting the appearance of the adaptive immune response.

While our study provides valuable insights into the early diversification of HIV-1 populations post-transmission, there are several limitations that must be acknowledged. First, the cross-sectional nature of the data restricts our ability to track longitudinal changes in the viral population over time. Although we were able to capture initial stages of viral evolution, a more comprehensive understanding of the dynamics of mutation accumulation would require repeated sampling across multiple timepoints during the early phase of infection. Additionally, our use of uSGS, while allowing us to dramatically increase the number of viral genomes analyzed, inherently sacrifices sequence length, which limits the resolution of certain genetic features. This is a particular issue with APOBEC3G/F since there are very likely to be many more mutations outside of those that were identified within these subgenomic regions. Further, while the use of PID-specific primers improves participant specificity, it may limit the detection of additional viral populations (*e.g.*, via co-infection) that contain polymorphisms. An example of this limitation was shown in a recent report where they identified an additional *env* variant in PID 6609 that was highly divergent from the TF initially observed (*48*). Also, although the *pol* and *env* segments we chose to sequence both provided useful information, the segments were separated and so could only be evaluated independently, without correlation or linkage analysis. Lastly, the model assumptions used in our Monte Carlo simulations, such as no immune pressure, likely do not fully reflect the complexity of host-virus interactions in real-world infections, where immune responses are likely to exert selective pressures.

Despite these limitations, our findings highlight the profound impact of stochastic HIV-1 RT error- and APOBEC3G/F-mediated processes on early HIV-1 evolution. The increased sensitivity provided by uSGS allowed us to gain a more comprehensive view of viral diversity, revealing a relatively homogenous population consistent with successful transmission of one or a few viral variants. Our analysis also confirmed that early viral evolution is strongly shaped by intrinsic mutation processes, with only limited evidence of immune-driven selection in acute infection. These results suggest that, while purifying selection may constrain the fixation of deleterious mutations, the early evolution of HIV-1 is primarily governed by random mutation, founder effects, and APOBEC3G/F mutations, and, perhaps, rare positive selection for some mutations. As such, while not necessarily being able to directly eliminate an established HIV reservoir, these data offers the possibility to explore the development of a ‘personalized medicine approach’ during this stage of acute infection to target the genetically homogenous viral population of an individual that persists for weeks after transmission.

## METHODS

### Study design

Plasma samples were from participants enrolled in the RV254/Southeast Asia Research Collaboration with Hawaii (SEARCH) 010 Trial (NCT00796146). The participant population has been previously reported in detail by de Souza *et al*. (*47*). In brief, the study was designed to identify acute HIV-1 infection in high-risk populations in Thailand and offer immediate initiation of ART. The trial was conducted by the Thai Red Cross AIDS Research Centre and the Department of Retrovirology, US Army Medical Component, Armed Forces Research Institute of Medical Sciences. All participants signed written informed consent and participated in protocols approved by Thai and US (Walter Reed Army Institute of Research) Institutional Review Boards. The investigators have adhered to the policies for protection of human subjects as prescribed in AR 70–25. The samples of this current study were initially collected between 2009-2015 at participants’s respective first visit; and were subsequently shared in September 2021 by the Henry M. Jackson Foundation component of the Military HIV Research Program. Samples were properly de-identified without the ability to identify individual participants during or after data collection.

### Plasma samples

Our laboratory obtained 250 μL of peripheral blood plasma from 15 donors with acutely acquired HIV-1 enrolled in the RV254/SEARCH 010 Trial. Plasma was collected from the participants prior to administration of ART. HIV-1 RNA levels in the plasma ranged from 2.5-36 million copies/mL. Ten of the baseline plasma samples were previously determined as single transmitted founder viruses by standard single-genome sequencing (SGS) of 8-12 full-length single viral genomes (*48*). The SGS of the full-length single genomes were provided to us by the SEARCH study team. During clinical work-up, the 15 participants had baseline plasma collected during Fiebig stages II-IV. Study participants were comprised of both men and women, of different ages, economic status and education level. The greatest risk factor for HIV-1 acquisition was through commercial sex work and same-sex relations between men.

### Ultrasensitive single-genome sequencing (uSGS)

Viral RNA was extracted from each sample using 10-30 μL of plasma per extraction for each subgenomic region of *pol* (HXB2 nt: 2,723-3,332; RT aa: 59-261) and *env* (HXB2 nt: 6,897-7,215; *env* aa: 125-331) (*96*) (**Figure S12**). cDNA libraries were generated with primer IDs (*i.e.*, unique molecular identifier, UMI) to label each unique cDNA molecule for next-generation uSGS as previously described in detail (*46*). By using uSGS, we eliminated PCR induced mutations and PCR recombination as well as resampling of the of the same templates to recover thousands of HIV-1 single genomes. The full-length SGSs provided from the SEARCH 010 study team provided a template from which to design donor-specific primers for subgenomic regions (**File S1**).

Primers for cDNA were designed to be donor and segment-specific with a 5′-terminal randomized (hand mixed) ID segment and PCR priming region, enabling unique tagging of each cDNA molecule during *in vitro* reverse transcription. We performed multiple extractions, 3-5 times for each sample and subgenomic region, which allowed us to obtain ∼10,000 single genomes. The cDNA synthesis reactions were purified and amplified with deoxyuridine-containing residues in donor-specific primers for each region. The deoxyuridine residues were enzymatically removed with the uracil specific reagent, USER II (NEB #M5508S) thereby leaving long single-stranded overhangs available for efficient directional ligation of Illumina adaptors and library construction.

Samples were prepared for MiSeq Illumina sequencing as directed in the protocol for the 600-cycle MiSeq v3 kit (MS-102-3003 Illumina Inc, San Diego, CA). After MiSeq runs were complete, FASTQ files were exported for bioinformatic analyses. Raw reads were first separated by donor-specific indexes and then binned according to their common primer IDs. The MiSeq paired-end reads were concatenated with the reverse complement sequence used for Read 2. Lower quality reads were removed if they did not satisfy the parameters of -Q20 -P90 (http://hannonlab.cshl.edu/fastx_toolkit). The unique primer ID/sample, with thousands of reads per run, were processed through a rigorous bioinformatics pipeline designed to eliminate artifacts from the dataset by satisfying the “supermajority rule” of 80% (*46*). A consensus sequence was then constructed where all PCR recombination, sequencing and PCR errors were eliminated. In addition, the stringent pipeline also eliminated sequences where the primer IDs had been mutated (*97*). Perl scripts used in the above analyses are available at the GitHub code repository at https://github.com/ShaoFred/MiSeq_consensus_builder.git.

Though our original experimental design was to sequence 500-600bp segments of *pol* and *env*, preliminary sequence analysis revealed much, skewed, and consistently lower variation among paired-end Illumina read 1 sequences relative to read 2 that could not be corrected by varying the duration or temperature of cDNA synthesis reactions, bioinformatic pipeline adjustments, or shifting the amplicon regions. Since read 2 covered sequence proximal to the cDNA synthesis primers and the affected read 1 was distal, we attributed this observation to viral RNA fragmentation (probably due to long-term plasma storage) and consequent truncated cDNAs that recombined with amplicons derived from mostly TF-derived amplicons during PCR. Fortunately, variation in read 2 sequences was almost always evenly distributed throughout the respective reads, indicating that cDNA primer-proximal sequences were spared from these effects, as our explanatory hypothesis would predict. Therefore, we discarded all read 1 sequences and performed our analyses exclusively on the cDNA primer-proximal 234bp segments of *pol* and *env* covered by read 2 using in-house Perl scripts.

### APOBEC3G/F data filtering

Each alignment of viral genomes, which included potential APOBEC3G/F mutations, was termed unfiltered. Separately, viral genome sequences with ≥2 GR>AR mutations were removed and termed GR-AR filtered data. One GR>AR mutation was permitted to remain as it was possible that the mutation was not APOBEC3G/F dervived, but rather, an HIV-1 RT error.

### Phylogenetic reconstruction

Collapsed sequences were aligned with MAFFT v7.490 (FFT-NS-1 200PAM/k=2 algorithm) (*98, 99*) in Geneious Prime ® 2025.1.2. Minor adjustments were performed manually. Unrooted radial p-distance neighbor-joining trees were reconstructed using MEGA 11 (*100*) and visually observed in radial format using FigTree v1.4.4 (https://tree.bio.ed.ac.uk/software/figtree/). Phylogenetic trees with SNP matrix and pruned branches for detailed analysis were rooted on the PID TF sequence.

### Genetic diversity analyses

The average pairwise distance (APD) was calculated by the pairwise deletion, p-distance method. In Hamming distance calculations, the input file was the collapsed (*i.e.,* unique genomes included). However, the calculations were based on the non-collapsed sequence number. The TF was set as the reference, meaning that, for each unique genome, Hamming distances were calculated relative to the TF, and not relative to one another. Transition vs. transversion (Ti/Tv) ratios were also calculated on the collapsed datasets, using the total sequence number as the multiplier. Again, each sequence was compared to the TF. The ratios of weighted non-synonymous to synonymous (d_N_/d_S_) were also determined from uncollapsed datasets. The references were the TF and the consensus/ancestral CRF01_AE sequence (2021 complete genome, https://www.hiv.lanl.gov/). HIV-1 drug resistance mutations were identified using HIV- DRLink with the Stanford HIV Database (https://hivdb.stanford.edu) (*101*).

### Reversion to consensus CRF01_AE

To quantify amino acid diversity and identify mutations relative to the TF reference sequence, we implemented a custom R script to process aligned amino acid FASTA files. For each sample, aligned sequences were read and converted to character vectors. Identical sequences were collapsed into unique representatives, preserving information about their abundance and annotation. Specifically, original collapsed sequence names were parsed to extract numeric identifiers when present, which were summed across identical sequences. Each collapsed sequence was renamed to concatenate the original name, the total number of identical sequences, the summed numeric value, and the number of distinct sequence identifiers contributing to that group.

The most frequently observed sequence in each alignment (the TF) was designated as the consensus reference. All other sequences were compared to this consensus to identify positional differences. A match at a given site was represented by a (“.”), while mismatches retained the original amino acid character. Gaps and stop codons were preserved but not included in mutation tallies. A summary reported the number of mismatches (mutations) per position across all sequences, and the total number of non-consensus amino acid substitutions observed. This provided the total number of different amino acid changes in the dataset. The number of positions in the TF that were different than the ancestral/consensus CRF01_AE were identified and any sequences with reversions back to consensus CRF01_AE were noted. The percentage of amino acid mutations that were reversions to the consensus CRF01_AE could be calculated by the number of positions that had a reversion to the total number of different amino acid changes in the dataset.

### Amino acid mutation frequency and mapping to CTL epitopes

Using the data matrix generated for the reversion analysis, the frequency of amino acid mutations at each position could be determined. Since the MHC-I HLA alleles were previously determined for each participant, epitopes mapped to within a region of 14 amino acids or less were considered. The types of epitopes were specific to HLA alleles (*e.g.*, A*02), polyreactive (*e.g.*, A*02/B*58), or host species when no MHC-I presenting molecule was defined (*e.g.*, human). The number of amino acid changes within CTL epitopes from the consensus CRF01_AE were tallied. Importantly, since many of the epitopes were not experimentally tested for CRF01_AE but rather for subtype B, we assumed that the epitope was present, regardless of the subtype used for testing. Epitopes were initially mapped to HXB2 and then pairwise aligned to the consensus CRF01_AE. In some cases, epitopes were excluded in the V3 *env* region since the presence of insertions/deletions skewed the epitope length between subtype B and CRF01_AE. In this way, the frequency of amino acid changes in the dataset that fell within a CTL epitope could be quantified relative to the total number of different amino acid changes in the dataset.

### Viral mutation analysis modeling

Several methods were used to explore simple viral evolution mutation modeling. An estimated inferred mutation rate based on the observed mutations was determined in addition to a TF-match Poisson estimate rate. We utilized Monte Carlo simulations to establish a null model of expected mutation accumulation based on HIV-1 RT error alone while adjusting for varying viral generation times. Finally, a simplified viral evolution sequence simulation was performed to account for viral generation time, HIV-1 RT error rate, and time post-transmission. Full details are reported (**Text S1**).

### Viral tropism of CRF01_AE

We developed a predictive XGBoost classifier machine learning model for CRF01_AE viral tropism based on the V3 loop sequence. Features that were included in the logic were position 25, net-charge, glycosylation motif, and CRF01_AE specific X4-tropic features. Global feature importance of the XGBoost model were performed for Gain, Coverage, and Frequency as well as SHAP with bootstrapping of model predictions. Evaluation of model performance was compared to Gene2Pheno (*102*) and PhenoSeq (*60*). Full details are reported (**Text S2**).

### Statistical analyses

When appropriate, descriptive statistics were reported for mean and standard deviation and median and interquartile range. Wilcoxon matched-pairs signed rank test (paired Wilcoxon test) was used when comparing *pol* and *env* values such as Ti/Tv ratios or %APD, and dN/dS. One-sample t-test was used when comparing inferred mutation rates to the HIV-1 RT single-step rate. Wilcoxon signed-rank test was used when comparing the frequency of observed vs. expected mutations to the hypothetical 100%. Examining the Poisson fit of Hamming distance data is described below. Specific statistical tests or modeling have been described elsewhere in the methods. The *Viral mutation analysis modeling* R scripts are available at https://github.com/aacapoferri/Viral-dynamic-modeling. Statistical tests are indicated either in the text or in the table/figure legends. Statistical tests were performed using Prism GraphPad 10.3.1 with additional analyses using R (version R.4.3.1).

We tested whether Hamming distance counts in acute infection follow a Poisson process, as expected under random, independent mutation accumulation, estimating the Poisson mean (*λ*) from each participant and region sequenced. The χ^2^-goodness of fit test rejected the Poisson model for the full data and again after removing genomes with APOBEC3G/F mutation signatures, as potential detectable departures from the Poisson distribution. Because our sample sizes were very large, the χ^2^-goodness of fit test gains high power and can render small, biologically minor deviations statistically significant (a consequence of the Law of Large Numbers). To assess overall distributional shape, we additionally applied a Kolmogorov–Smirnov test to the empirical cumulative Hamming distance distribution. Although the Kolmogorov–Smirnov test formally assumes continuity while Hamming distances are discrete, it provides a conservative investigation and did not reject the Poisson distribution. Taken together, the Hamming distance distributions were determined as ‘Poisson-like’. In other words, the global pattern is consistent with a Poisson process, but subtle deviations are detectable at our sample sizes, even after removing APOBEC3G/F-associated sequences. Since the Kolmogorov–Smirnov test assumes a continuous distribution while our data are discrete, its acceptance of the Poisson distribution should be interpreted with this caveat. For transparency, we report both χ^2^-goodness of fit and Kolmogorov–Smirnov statistics for each plot in **Figure 2**.

## SUPPLEMENTAL MATERIAL

**Text S1.** Viral mutation analysis modeling

**Text S2.** Viral tropism of CRF01_AE

**File S1.** Primer sequences

**File S2.** GenBank accessions

## AUTHOR CONTRIBUTIONS

Conceptualization: AAC, VFB, JWM, JMC, JWR, MFK Data Curation: AAC, VFB, WS,

Formal Analysis: AAC, VFB, WS Funding Acquisition: MFK Investigation: AAC, VFB, WS, CH Methodology: AAC, VFB, WS

Project Administration: RT, NP, LT, SV, CS, SS, MFK Supervision: MFK

Visualization: AAC, VFB, WS, JWM, JCM, JWR, MFK Writing—Original Draft Preparation: AAC, VFB, WS, JMC, MFK

Writing—Review & Editing: AAC, VFB, WS, CH, RT, NP, LT, SV, CS, SS, JWM, JMC, JWR, MFK

## DATA AVAILABILITY

Data are available in the article itself and its supplementary materials. All sequence data are available in a public repository at GenBank. Previous near full-length plasma single genome sequences and uSGS sequences were deposited under the following accessions (**File S2**): (MG989627, 2543928P000L_Sb; MG989502, MK272357- MK272366; MG989521, MK272383- MK272391; MG989626, 2543698P000L_Sa(254031P00La); MG989608, MG989658, MK272520-MK272528, MK272550-MK272558; MG989527, MK272672-MK272679; MG989495, 2543513P000FL_Sj_(254056P00j); MG989542, 2547905P000FL_Sb(254060P00b); MG989561, 254114P00A; MG989492, 2543193P00C; MG989538; MG989568, MK272432- MK272440; MG989537, MK272395-MK272403; MG989554, MK272423-MK272431. Newly generated sequences from the ultrasensitive single genome sequencing assay can be found as (**in the process of submission**). Any additional data is available upon request.

## ACKNOWLEDGEMENTS

We would like to thank the study participants who committed so much of their time for this study. The participants were from the RV254/SEARCH 010, which is supported by cooperative agreements (WW81XWH-18-2-0040) between the Henry M. Jackson Foundation for the Advancement of Military Medicine, Inc., and the US Department of Defense (DOD) and in part, by the, National Institute of Health (DAIDS, NIAID, NIH grant AAI21058-001-01000). Antiretroviral therpay for RV254/SEARCH 010 participants was supported by the Thai Government Pharmaceutical Organization, Gilead Sciences, Merck, and ViiV Healthcare. We would also like to thank Drs. Brian T. Luke, Brandon F. Keele, Taina T. Immonen, and Morgane Rolland for invaluable discussions. We would also like to thank Meera Bose, AnneMarie O’Sullivan, and Eric Sanders-Buell for their efforts in curating and submitting RV254/SEARCH 010 near full-length sequences.

## DISCLAIMER

The views expressed here reflect the results of research conducted by the author(s) and do not necessarily reflect or construed to represent the official policy or be construed of the National Cancer Institute, National Institutes of Health, Department of Health and Human Services, Defense Health Agency, Department of War, US Government, or Henry Jackson Foundation. The study protocol was approved by the relevant Institutional Review Board(s) in compliance with all applicable Federal regulations governing the protection of human participants. The investigators have adhered to the policies for protection of human subjects as prescribed in AR-70-25.

---

**Figure S1.**
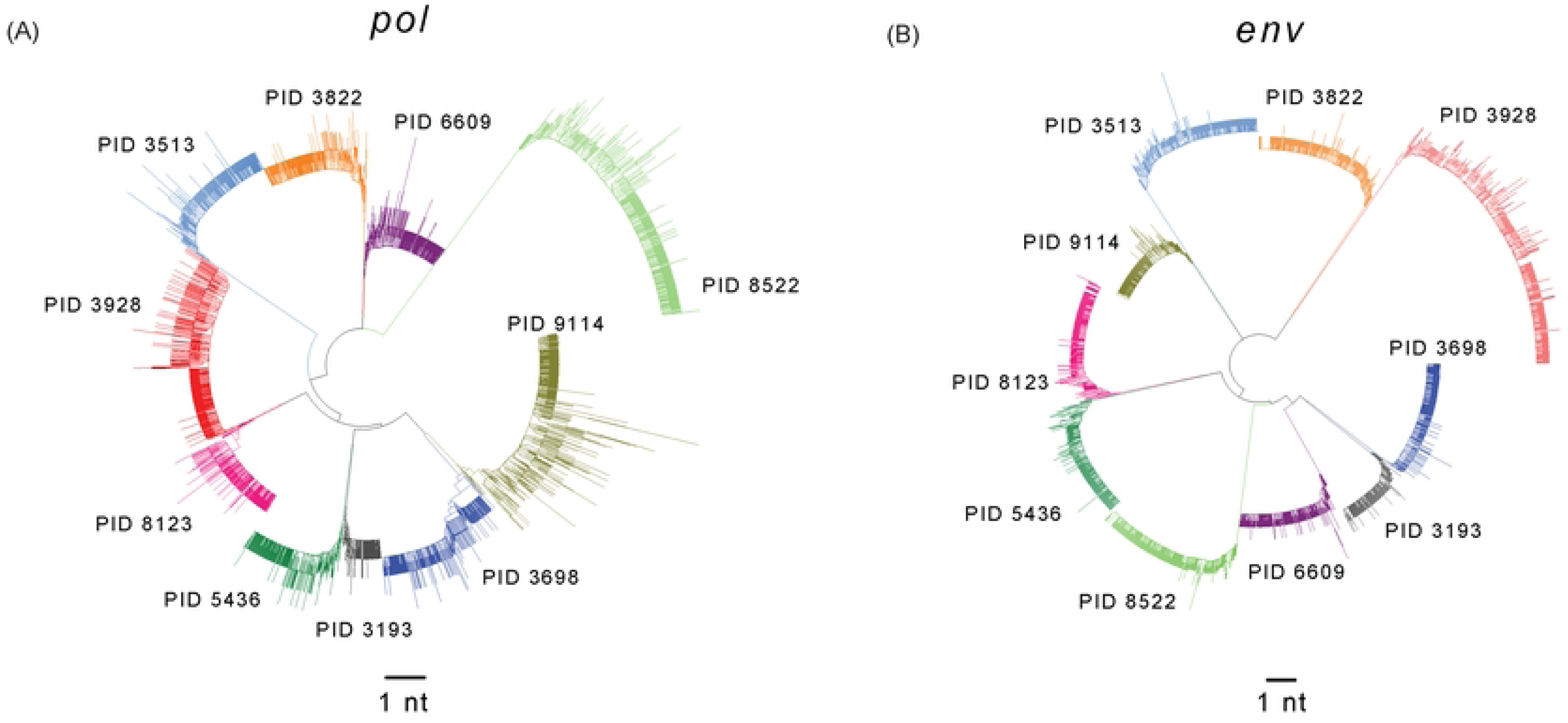
Sequence validation phylogenies for inferred single TF virus participants. Neighbor-joining phylogenetic reconstructed trees for (**A**) *pol* and (**B**) *env*. Each participant identifier is indicated and branches colored, respectively. The scale is set to 1 nt.

**Figure S2.**
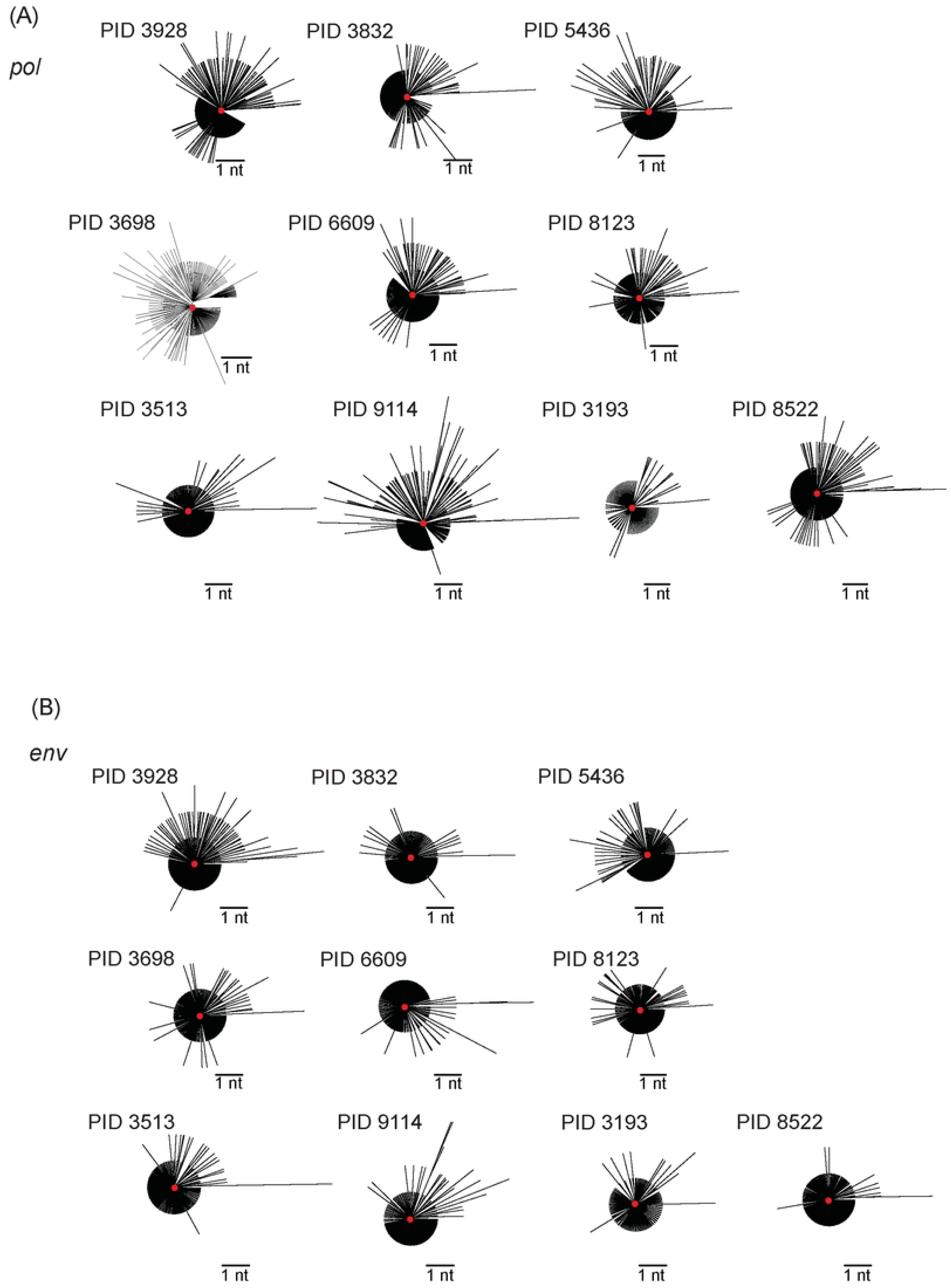
Phylogenies in inferred single TF participants. (**A**) Reconstructed phylogenies for the *pol* region. (**B**) Reconstructed phylogenies for the *env* region. The red dot in the center of the viral populations indicate the inferred TF. The participant identifier is noted next to the respective phylogeny. The scale is set to 1 nt.

**Figure S3.**
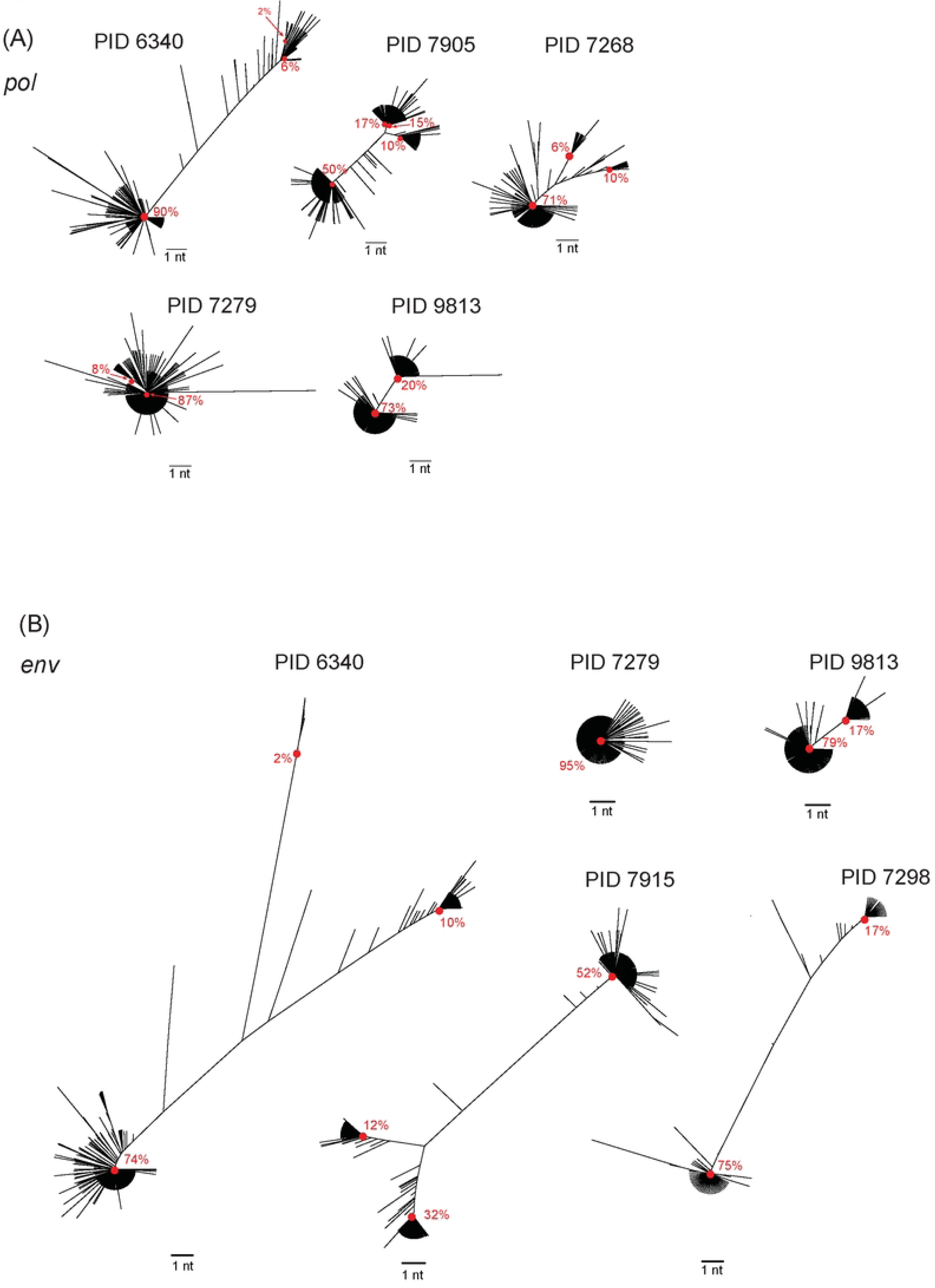
Phylogenies in inferred multiple TF participants. (**A**) Reconstructed phylogenies for the *pol* region. (**B**) Reconstructed phylogenies for the *env* region. The red dots in the center of the viral populations indicate the inferred TF. The proportion of viral genomes that comprise of the TF virus is noted as a percent. The participant identifier is noted next to the respective phylogeny. Long branches have been removed. The branches intermediate between the clusters reflect inferred recombinants between two TFs. The scale is set to 1 nt.

**Figure S4.**
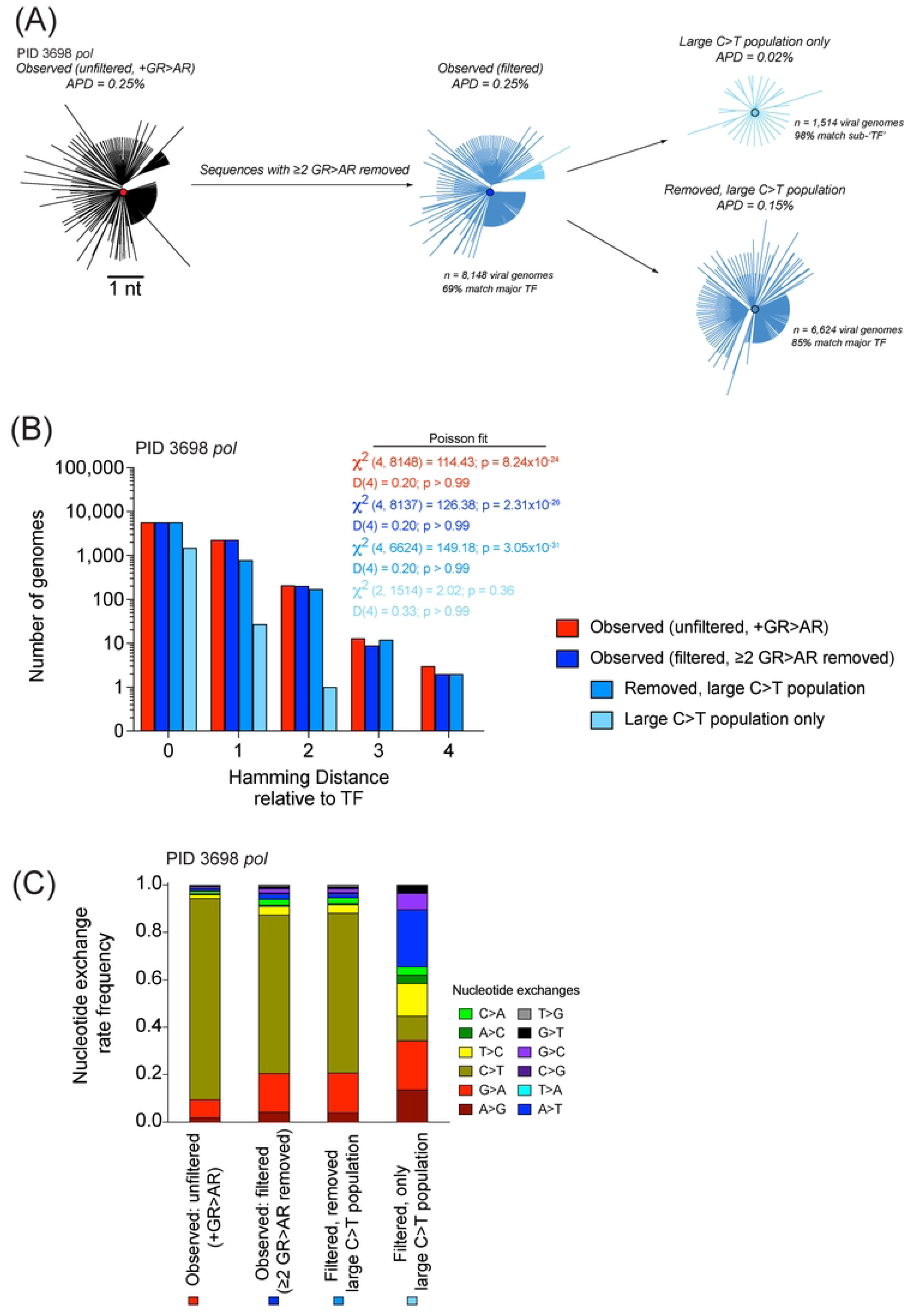
Examination of subpopulation in PID 3698 *pol*. (**A**) The star phylogeny as found in Supplemental Figure 2A. The dataset underwent GR-AR filtered with the two populations noted as two shades of blue. The two populations were split for each respective star phylogeny. The average pairwise distance is reported for each tree. The number of viral genomes with their respective percent that match the ‘main’ TF (dark blue) or subpopulation TF (light blue). The large subpopulation are a set of viral genomes that all contain a synonymous C>T change. (**B**) Hamming distance plots for the parsed datasets. Hamming distances were calculated relative to the respective inferred single TF virus. Bars indicate the number of viral genomes sequenced in unfiltered (red), GR-AR filtered (bright blue), without the large C>T population in GR-AR filtered (sky blue), and large C>T population only in GR-AR filtered (pale blue) data. The expected number of viral genomes was determined as a probability based on the Poisson fit of each respective observed dataset. The χ^2^-goodness of fit test and Kolmogorov-Smirnov test are reported for all respective datasets. (**C**) Nucleotide exchange rate frequencies of populations for PID 3698. Bars colors next to the names match the data filtering as above. Data from the unfiltered observed is from **Figure 3A**.

**Figure S5.**
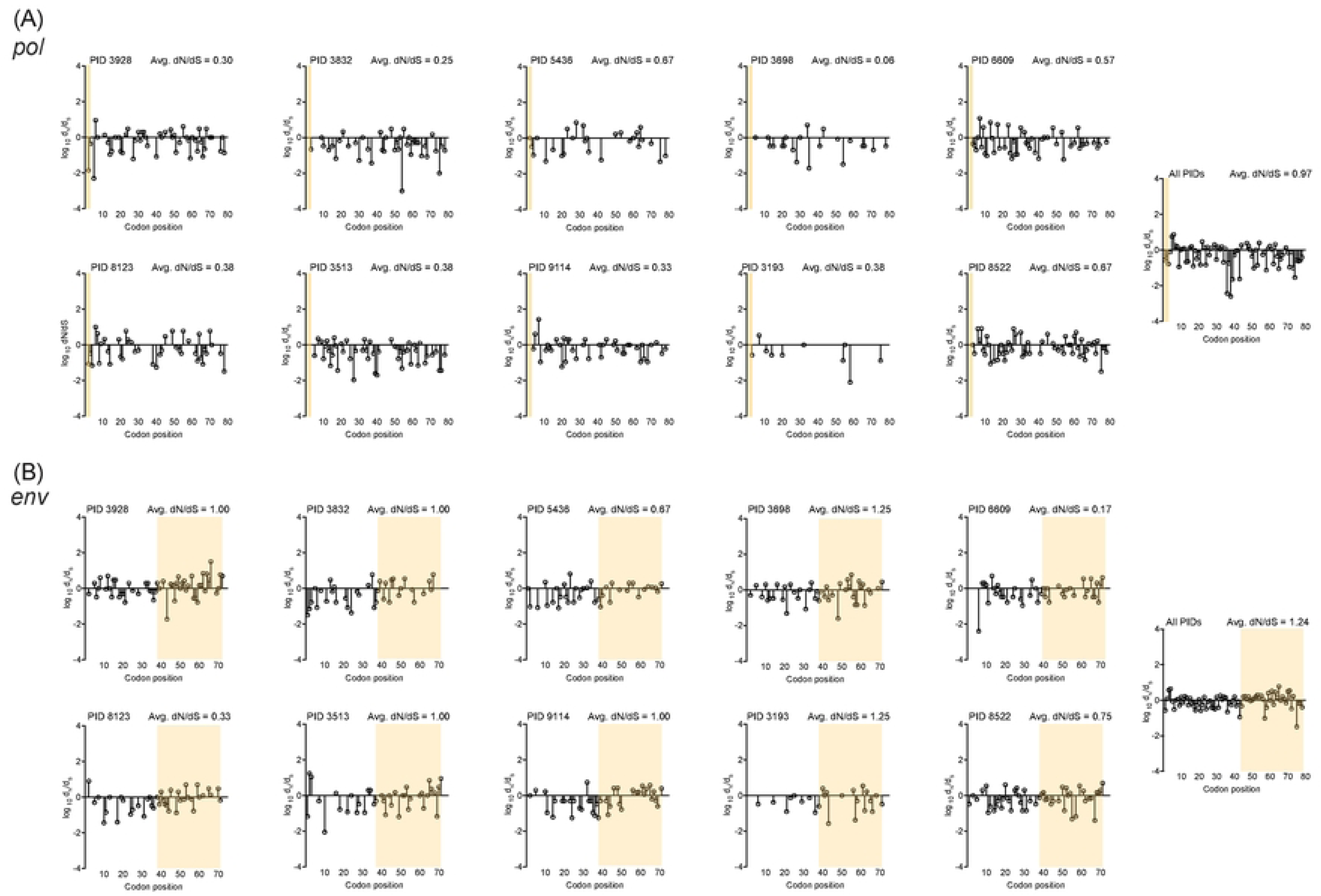
Selection of non-synonymous to synonymous (d_N_/d_S_) mutations across sequenced HIV-1 genomes. (**A**) d_N_/d_S_ lollipop plots for the *pol* region where the shaded region highlights the (Y)MDD codons 183-186, the highly conserved tyrosine residue was not captured in all the sequencing read. Since there is only one codon for methionine, the dS is always 0. (**B**) d_N_/d_S_ lollipop plots for the *env* region where the shaded area highlights the V3 region. Each PID is indicated on each plot. Calculated d_N_/d_S_ values were log_10_-transformed with positive (> 1), neutral (= 1), and purifying (< 1) values indicated. Codon position is relative to the sequence length. The average dN/dS across all participants for *pol* and *env* are reported.

**Figure S6.**
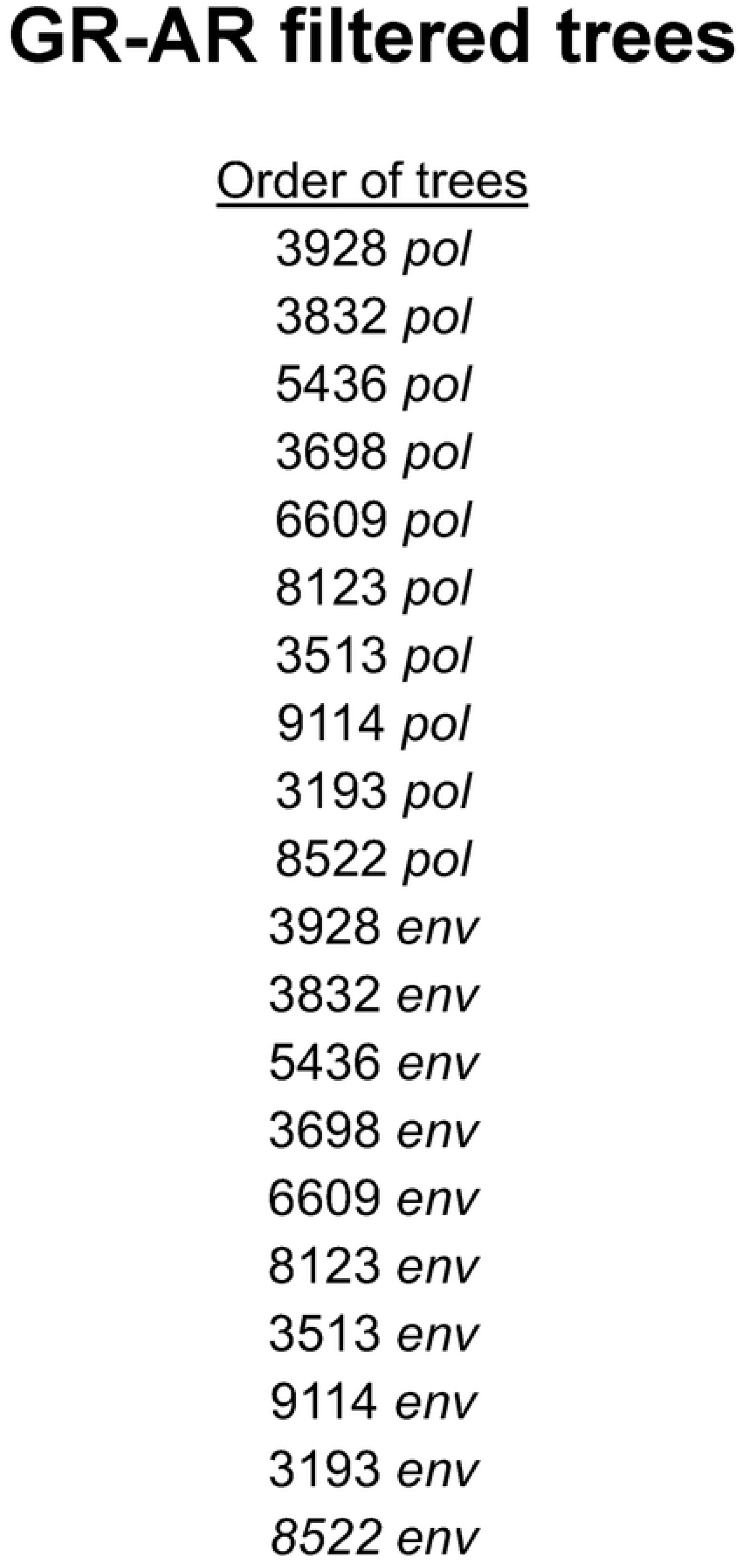
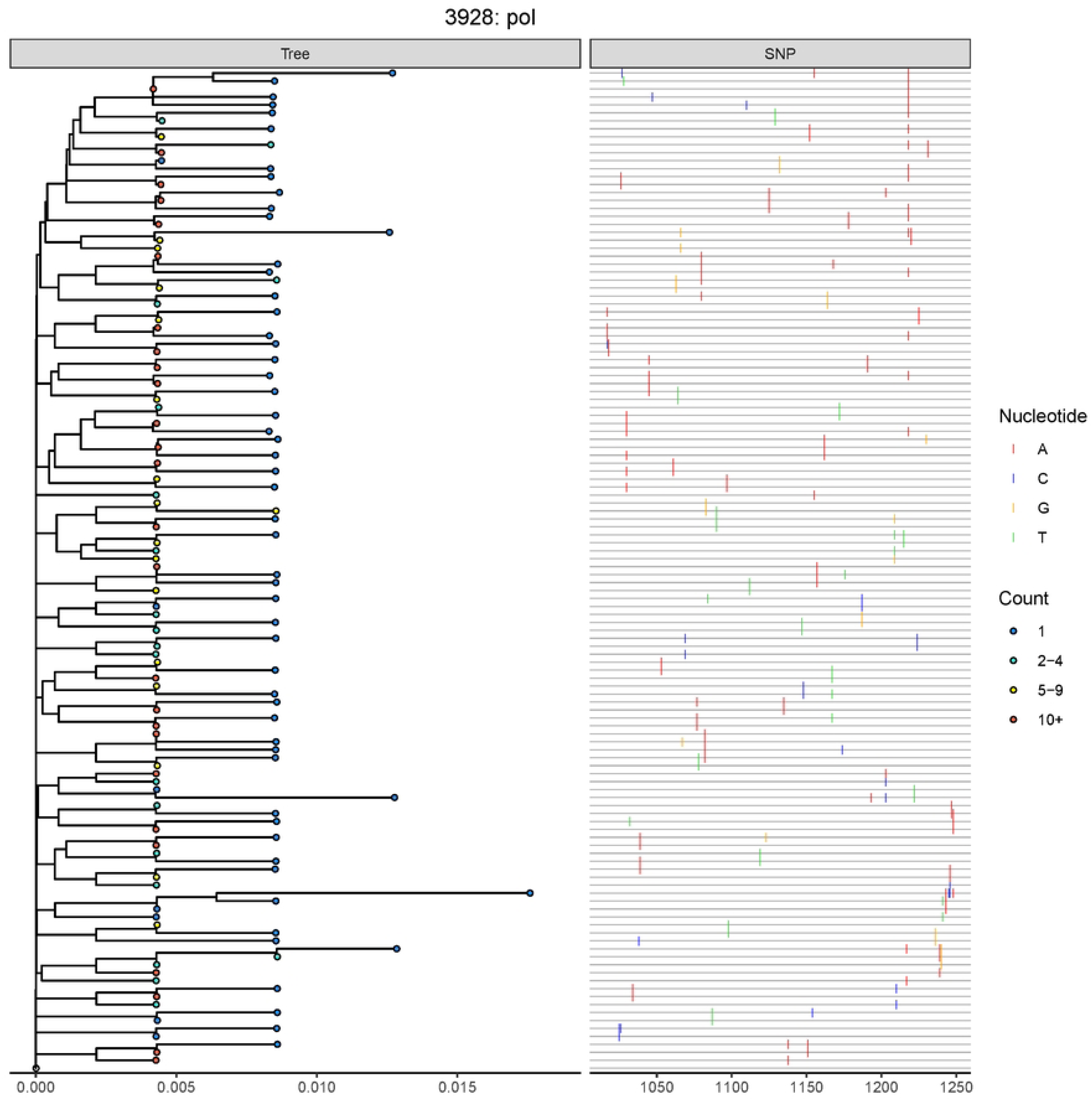
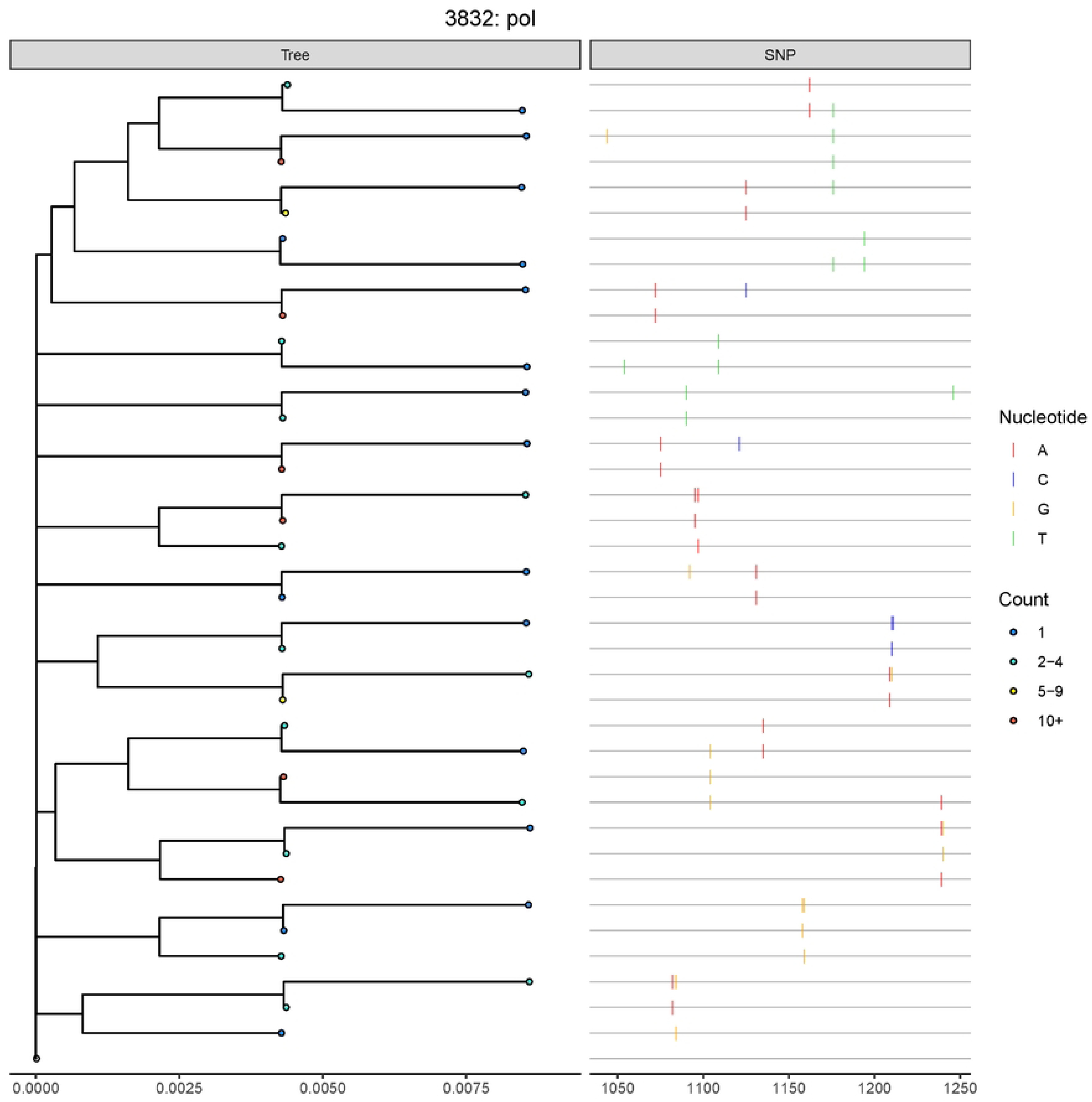
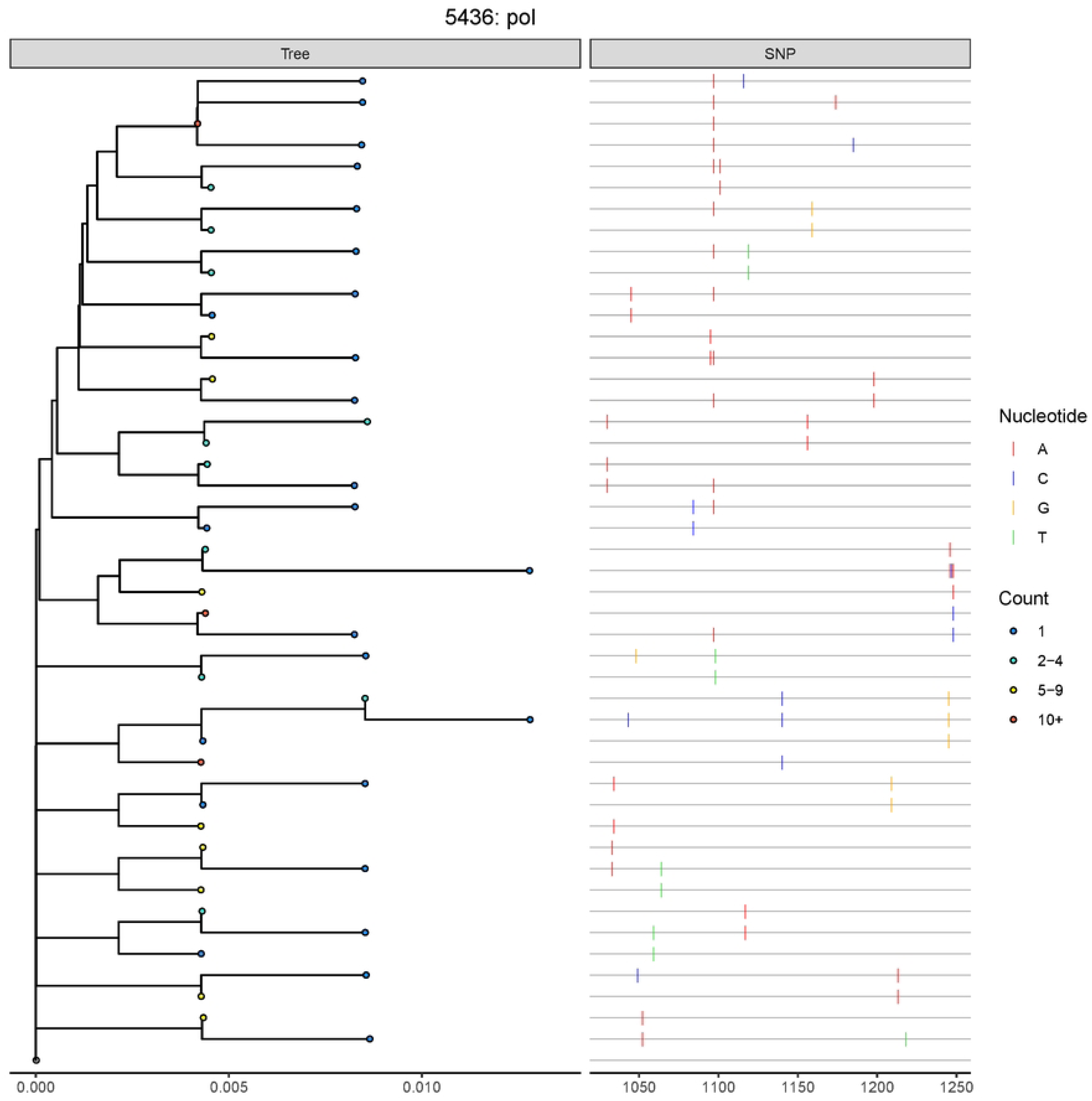
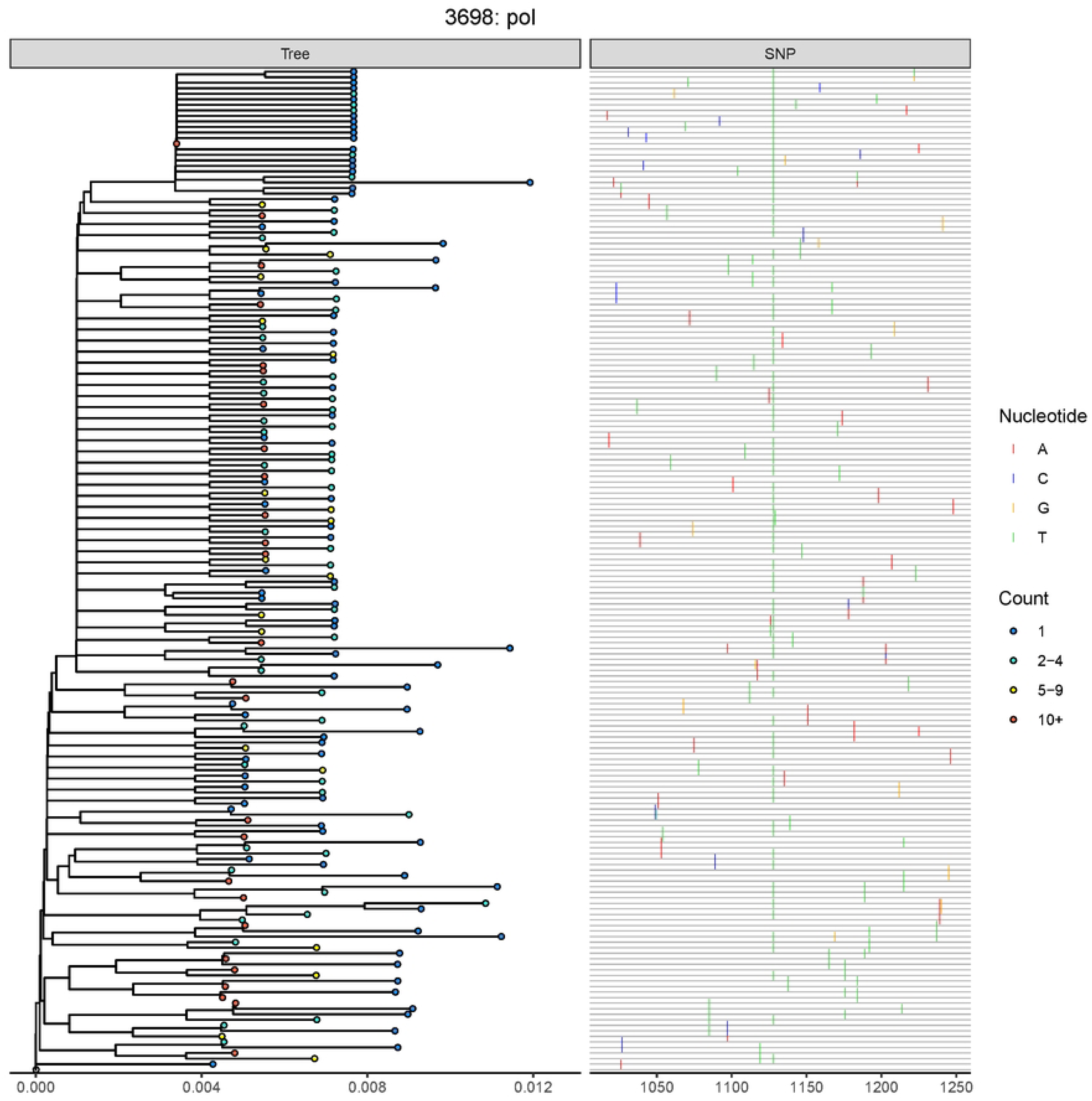
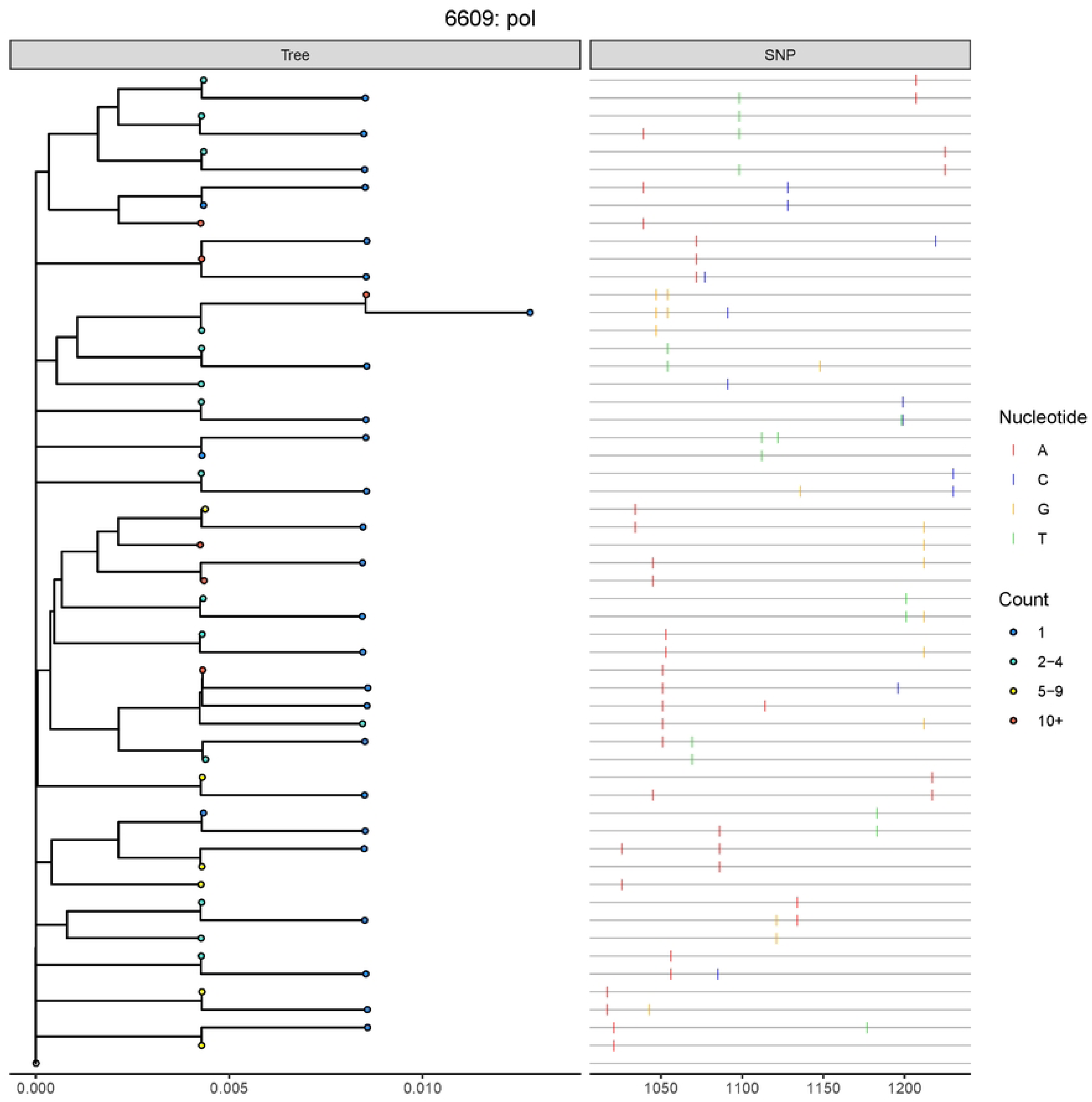
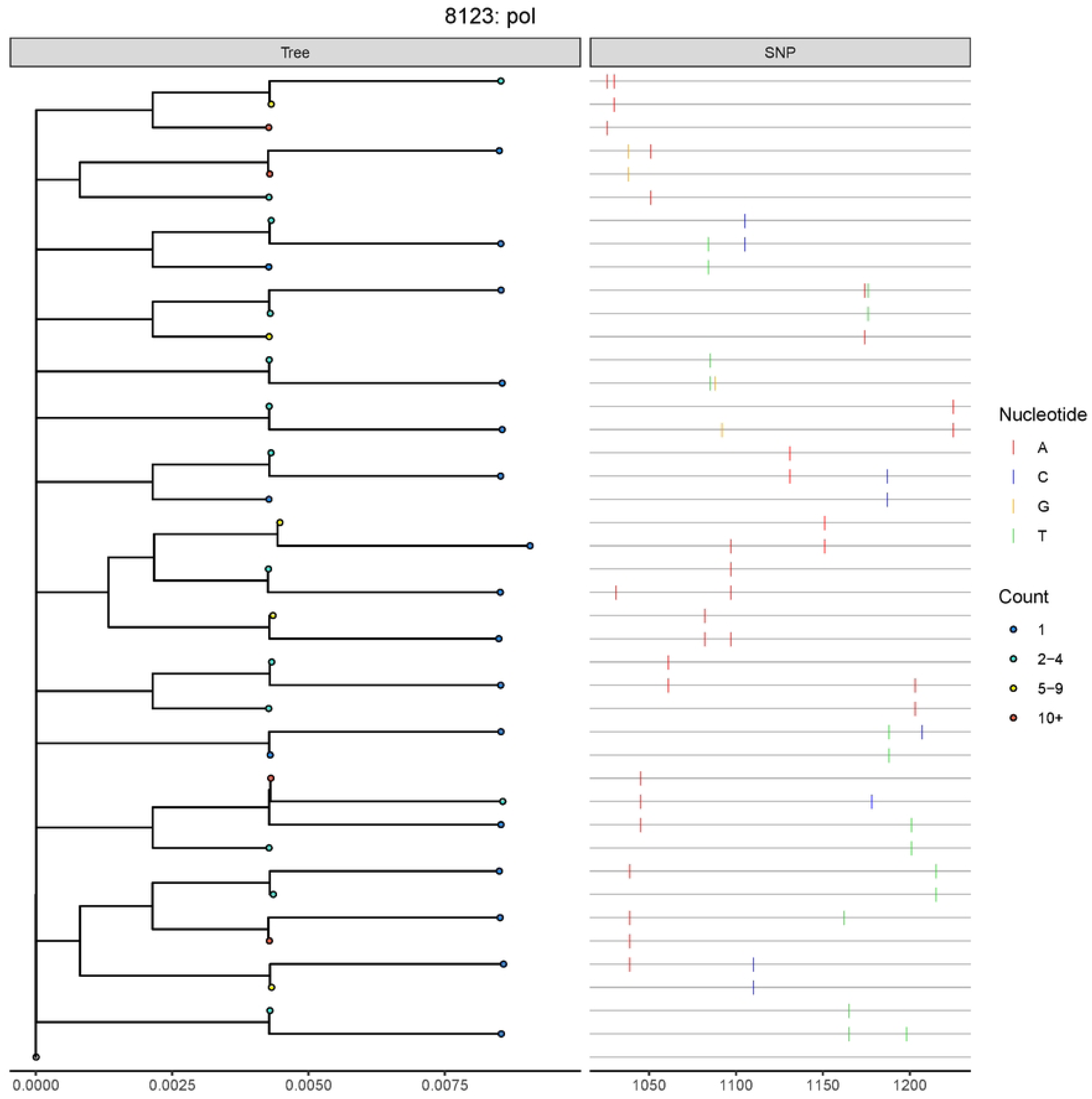
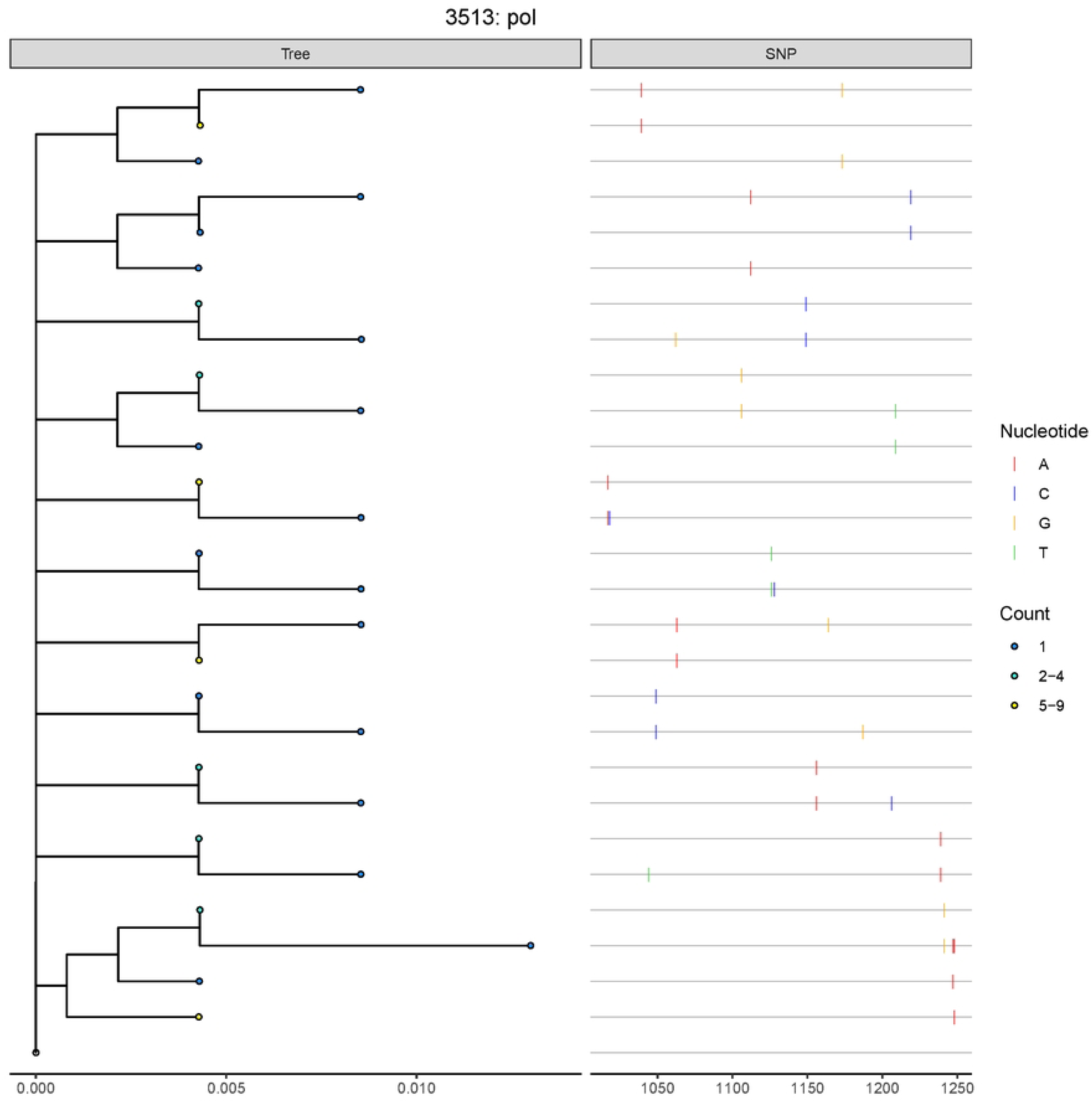
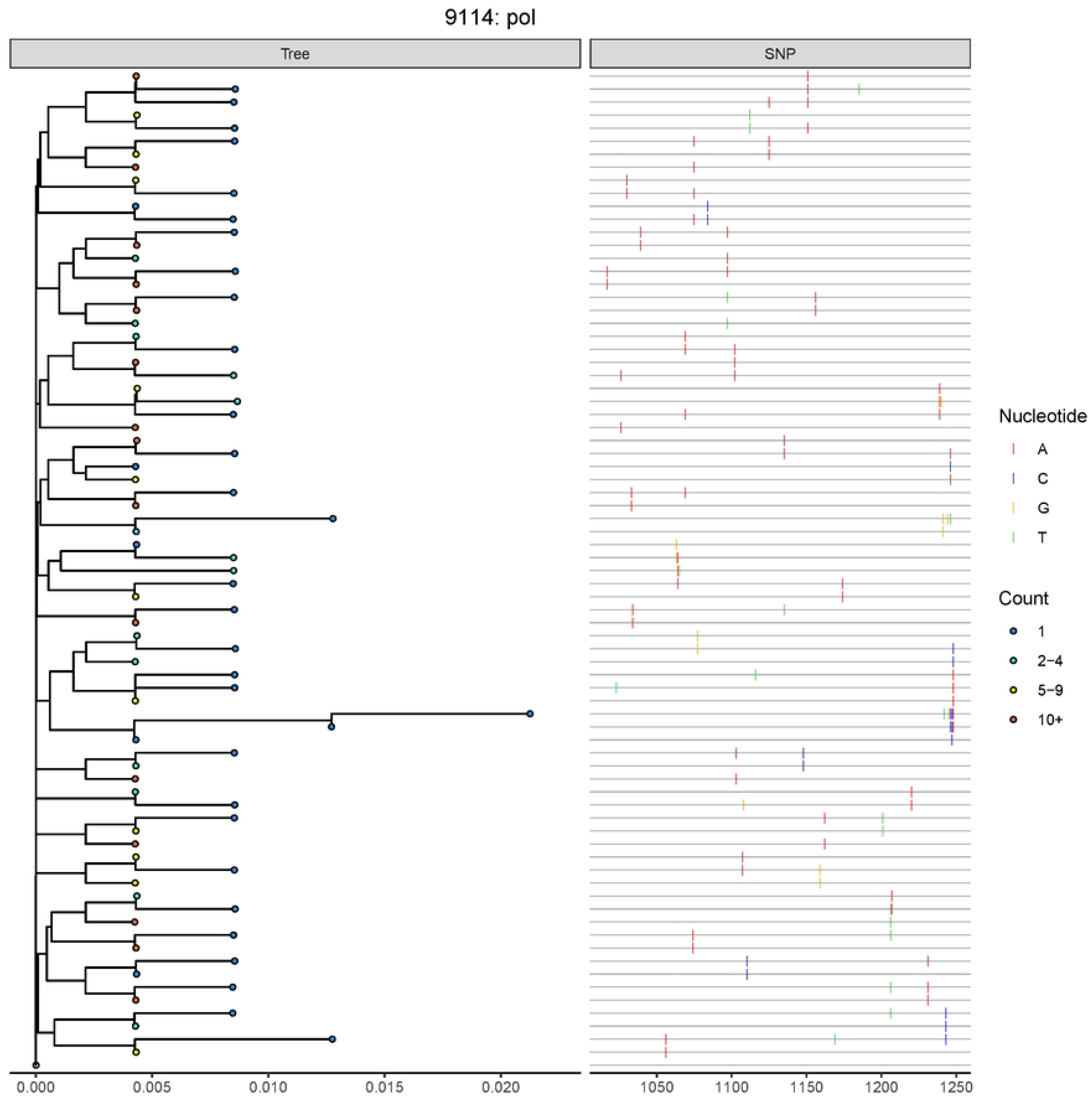
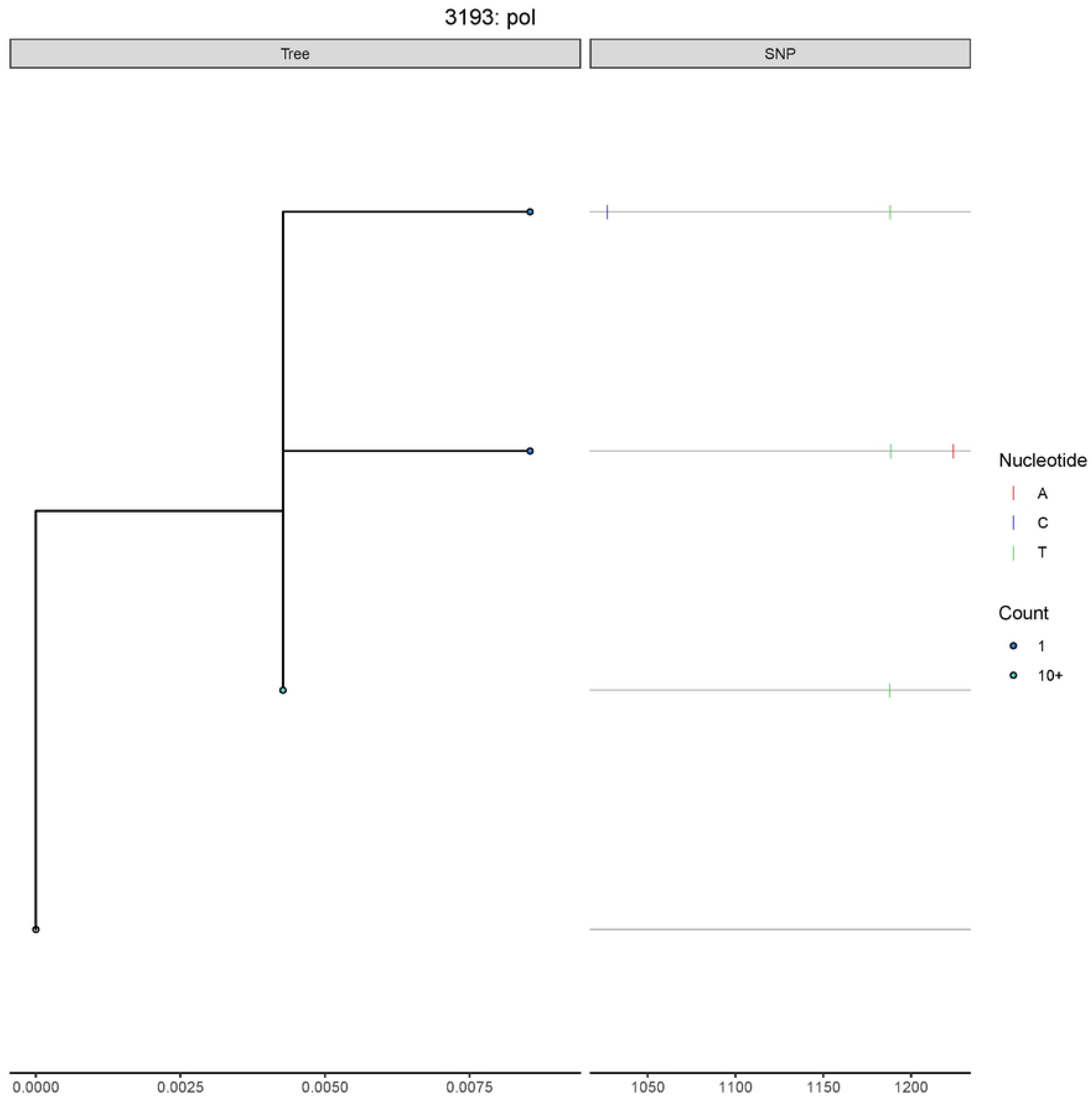
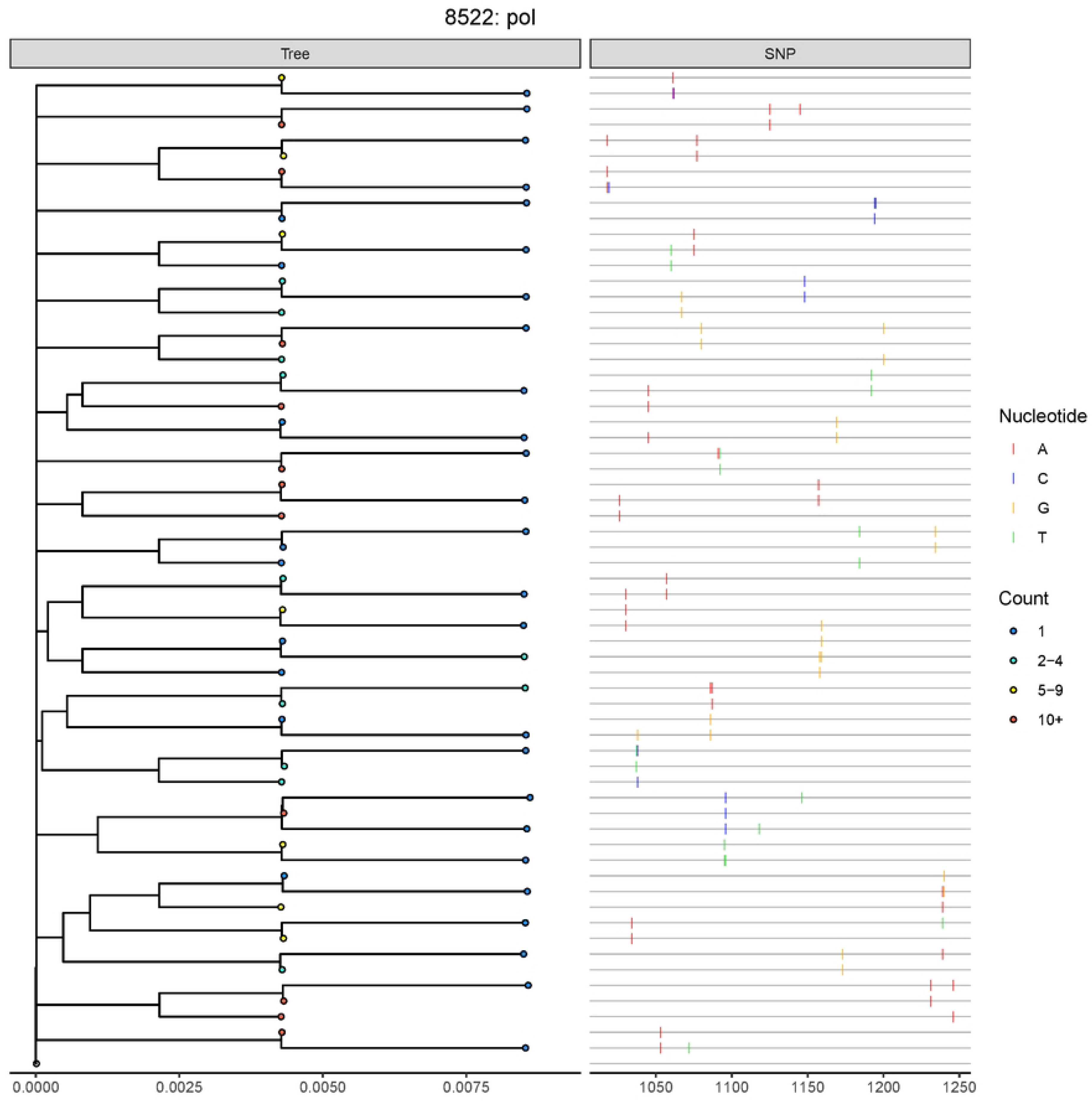
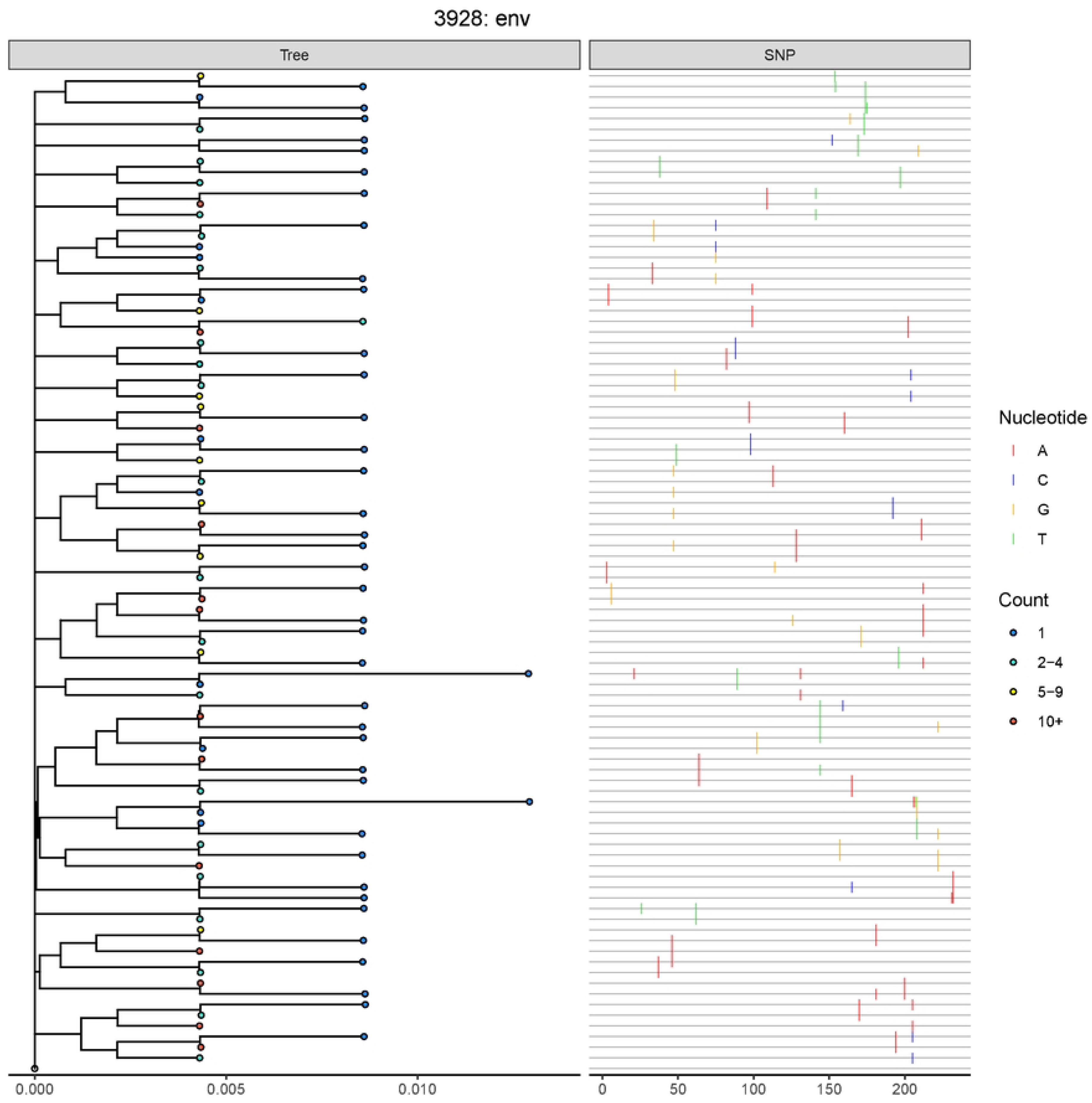
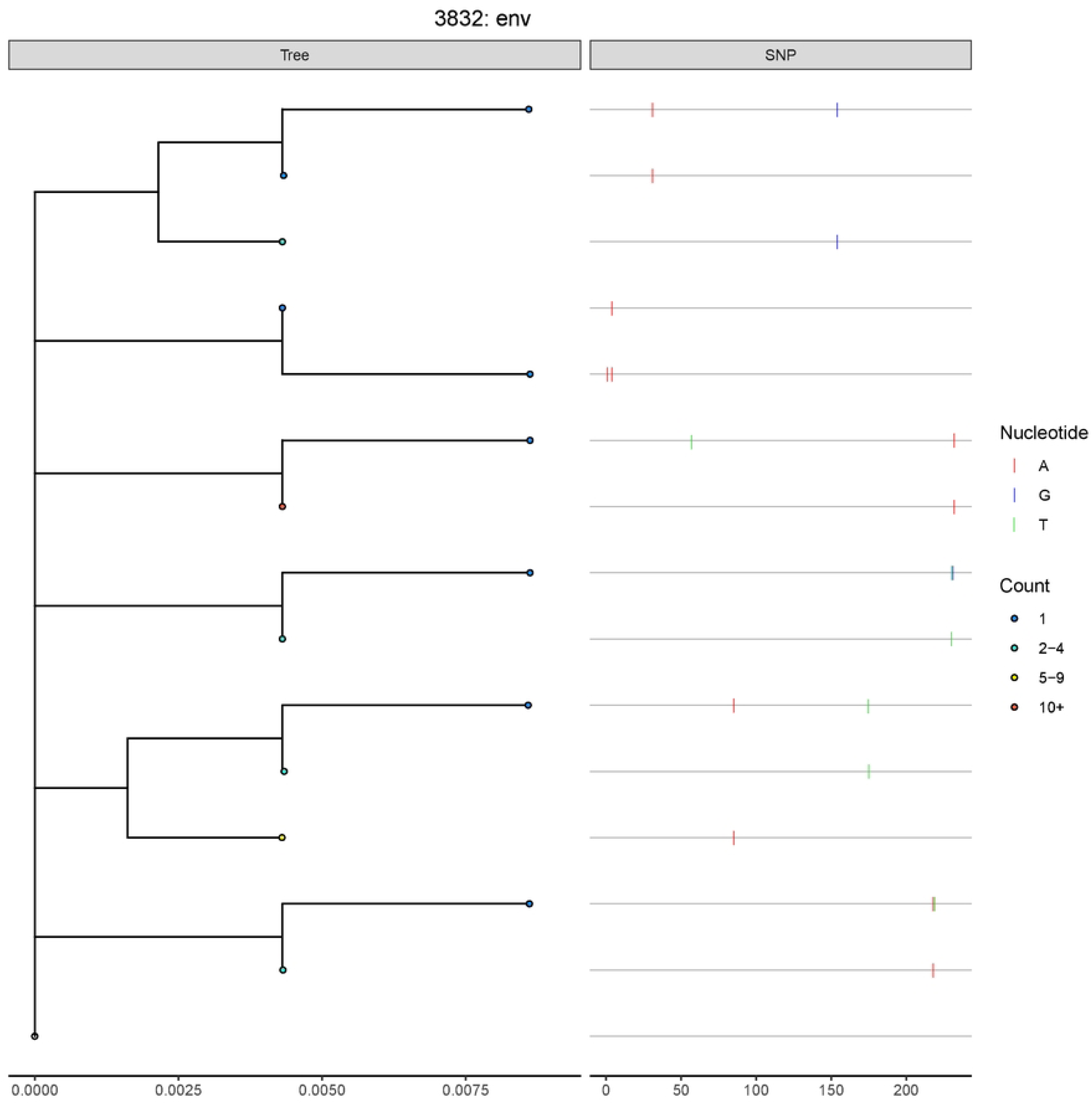
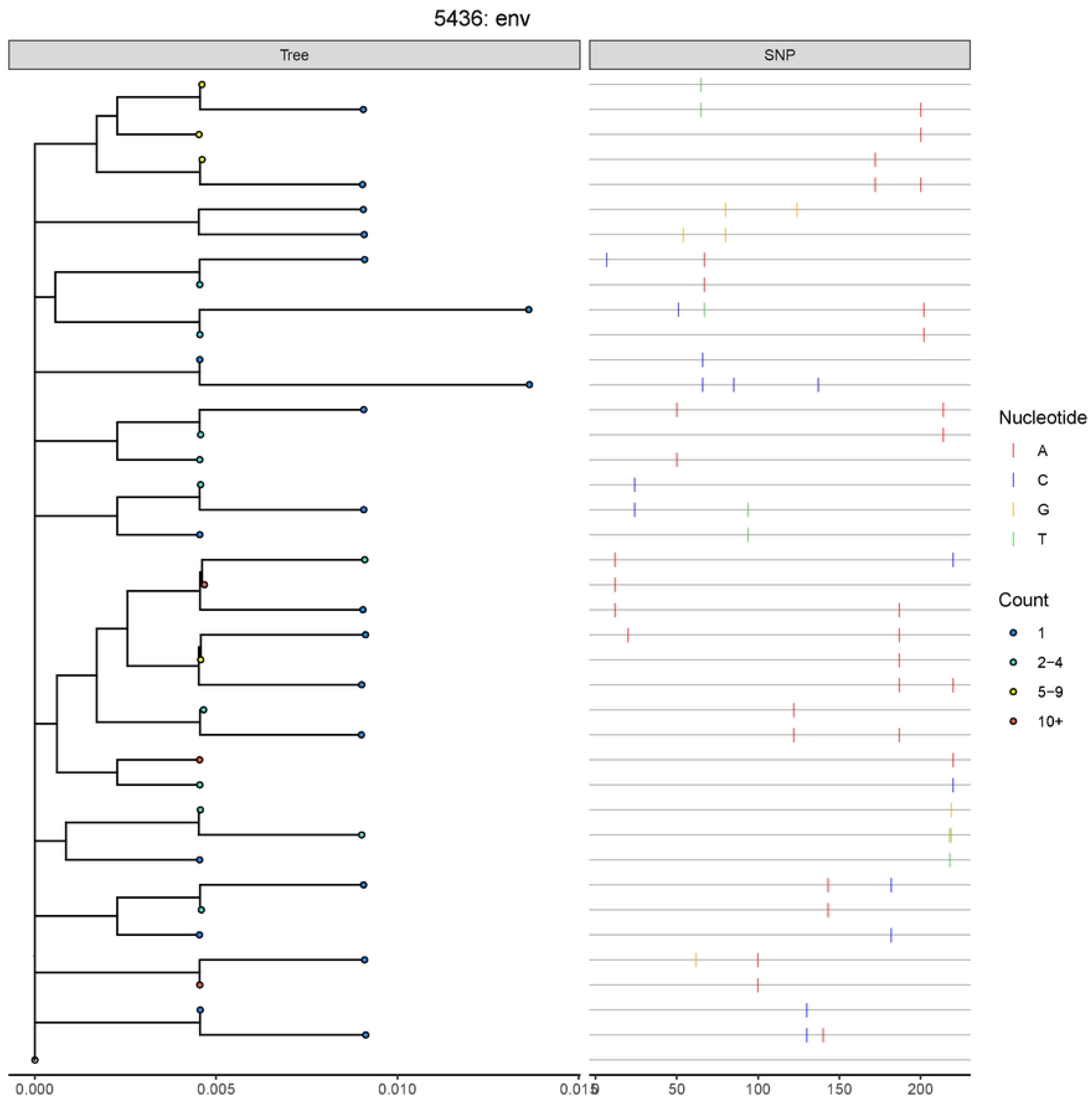
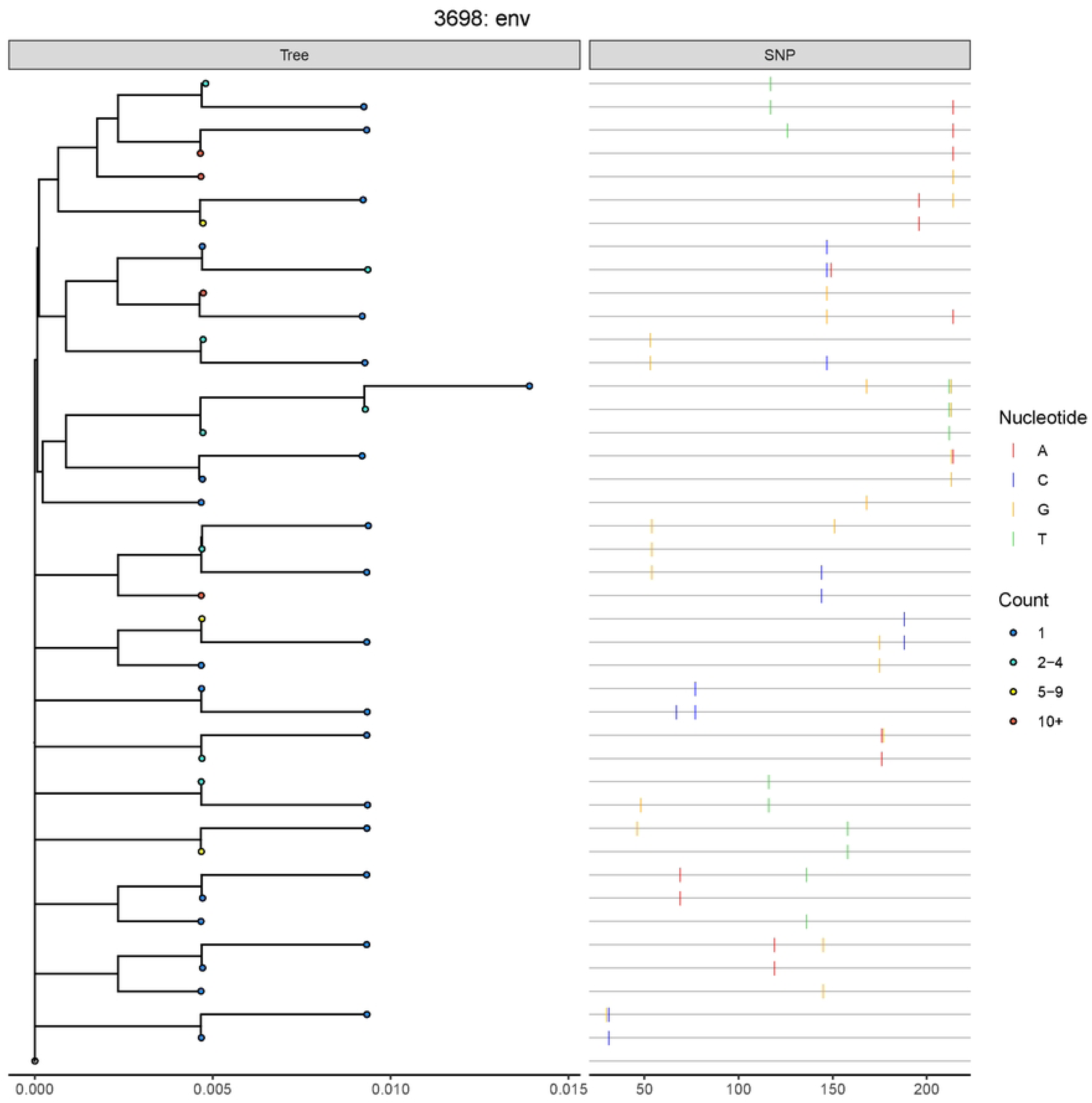
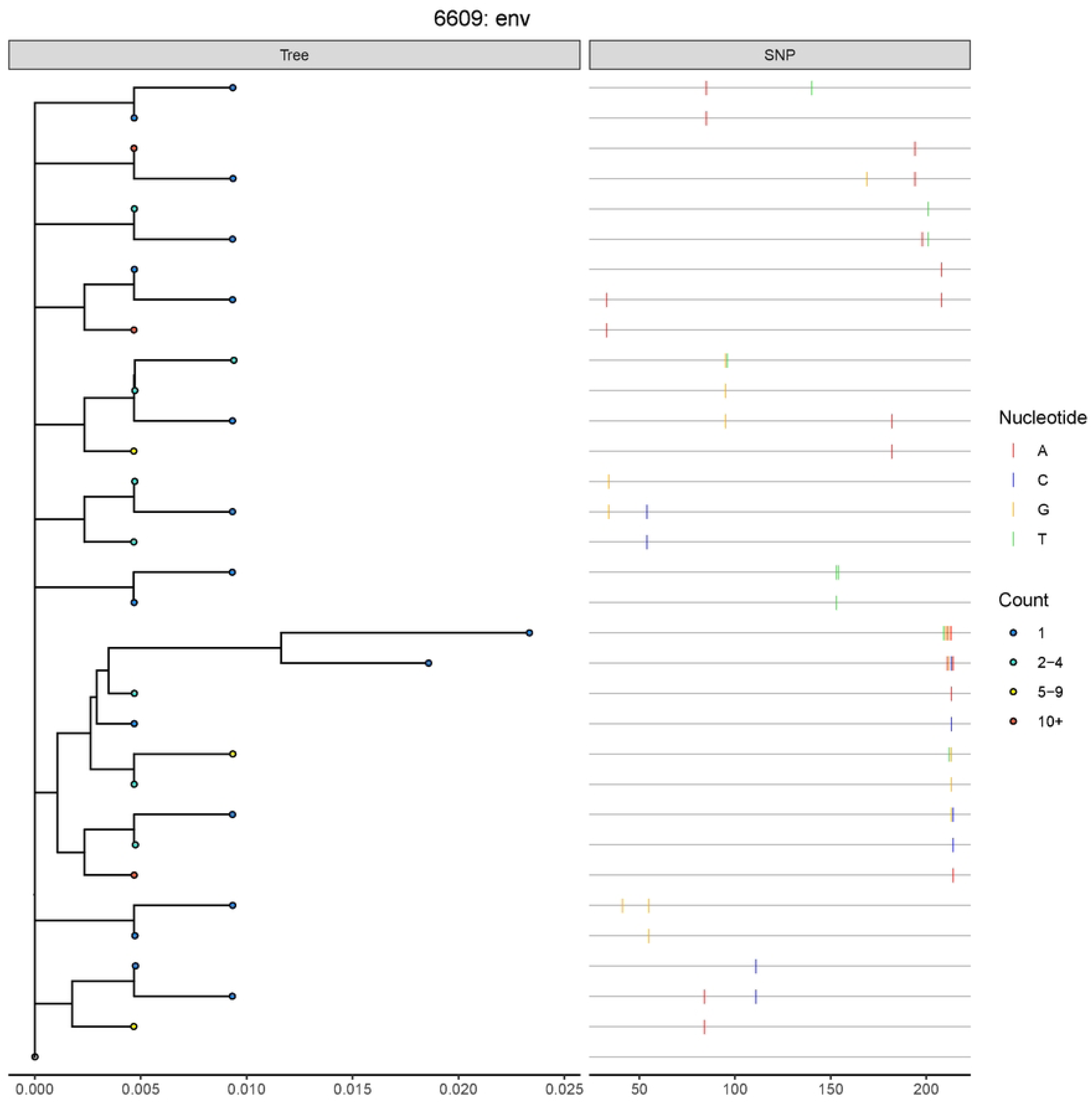
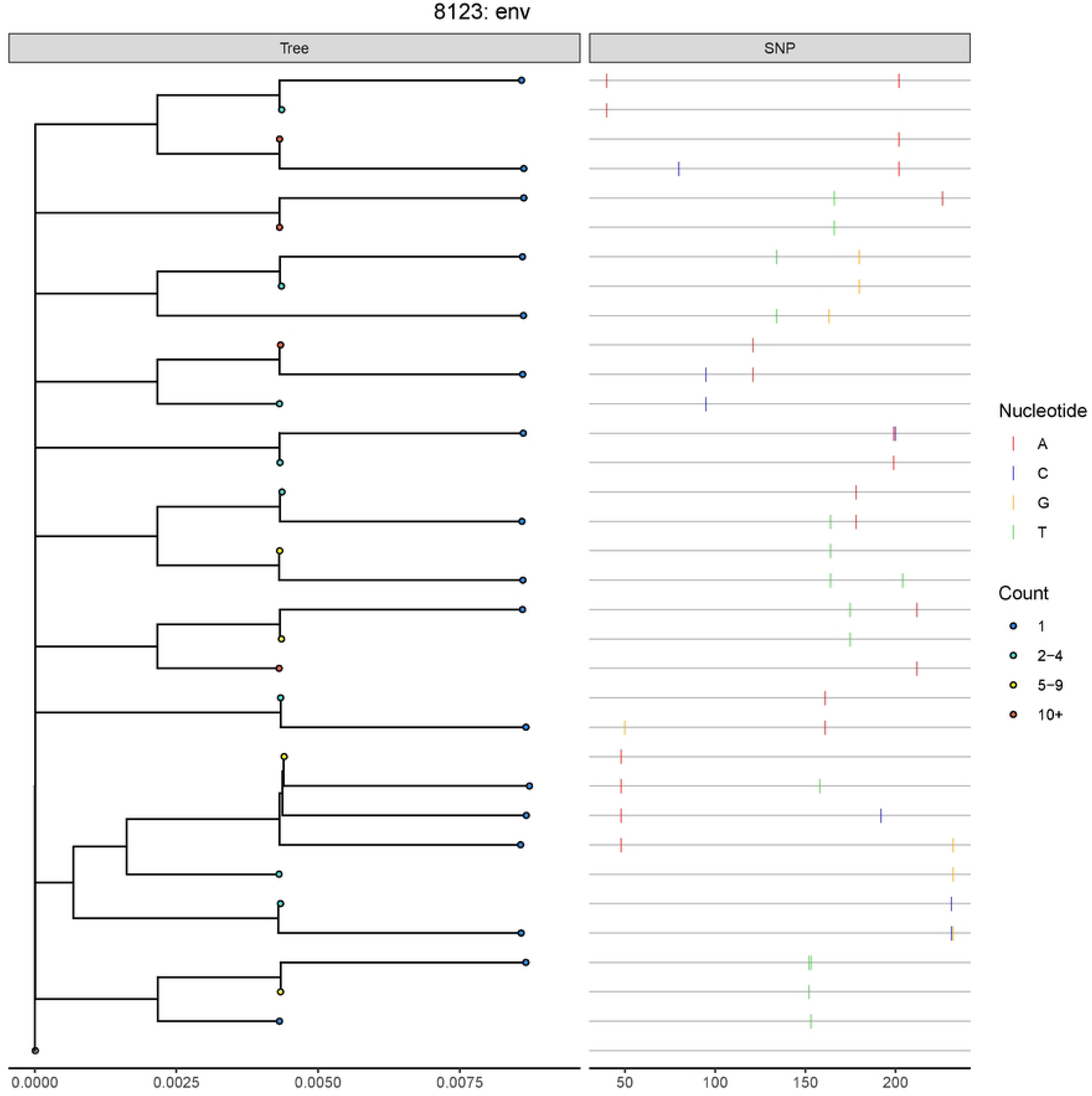
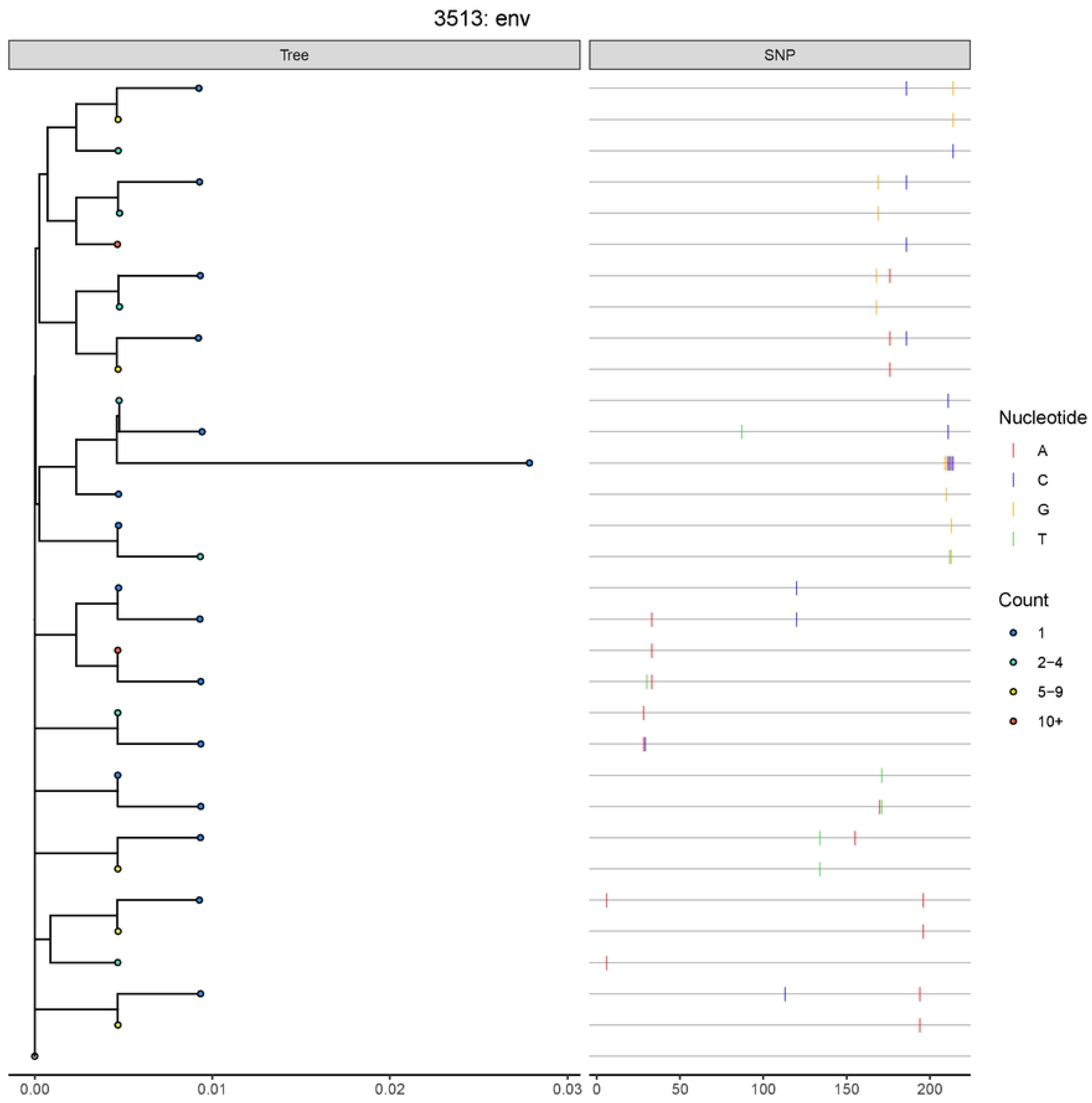
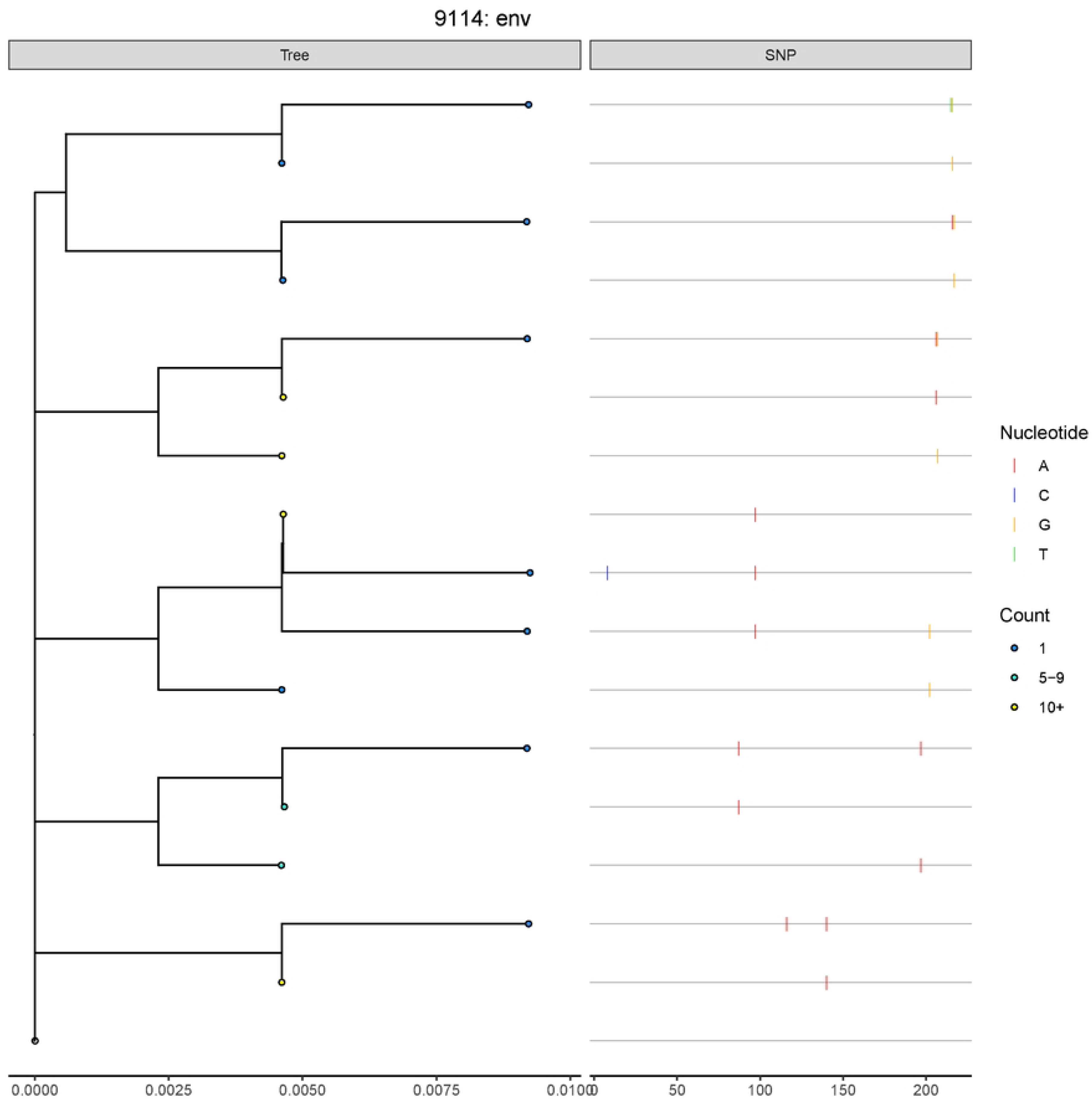
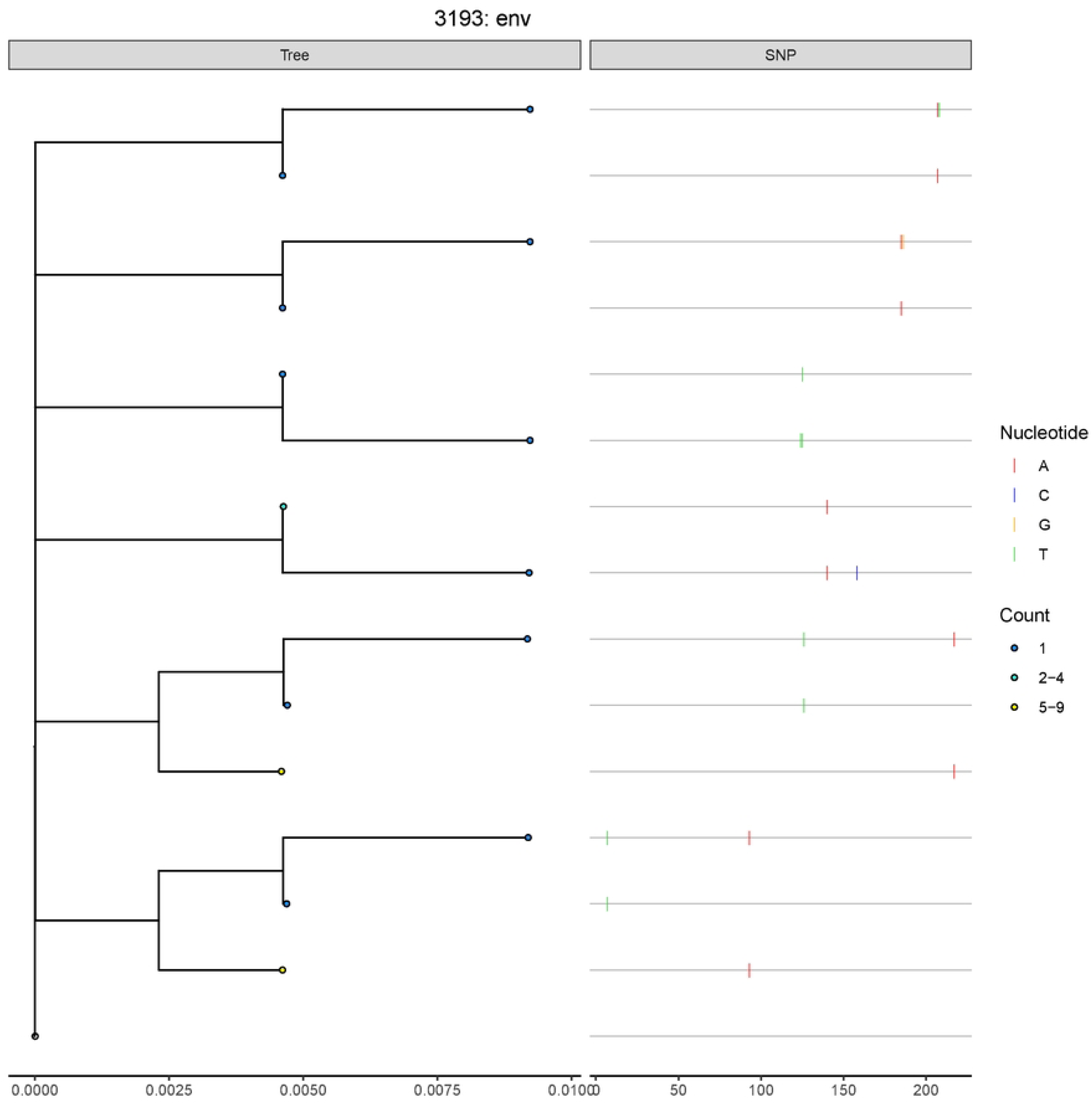
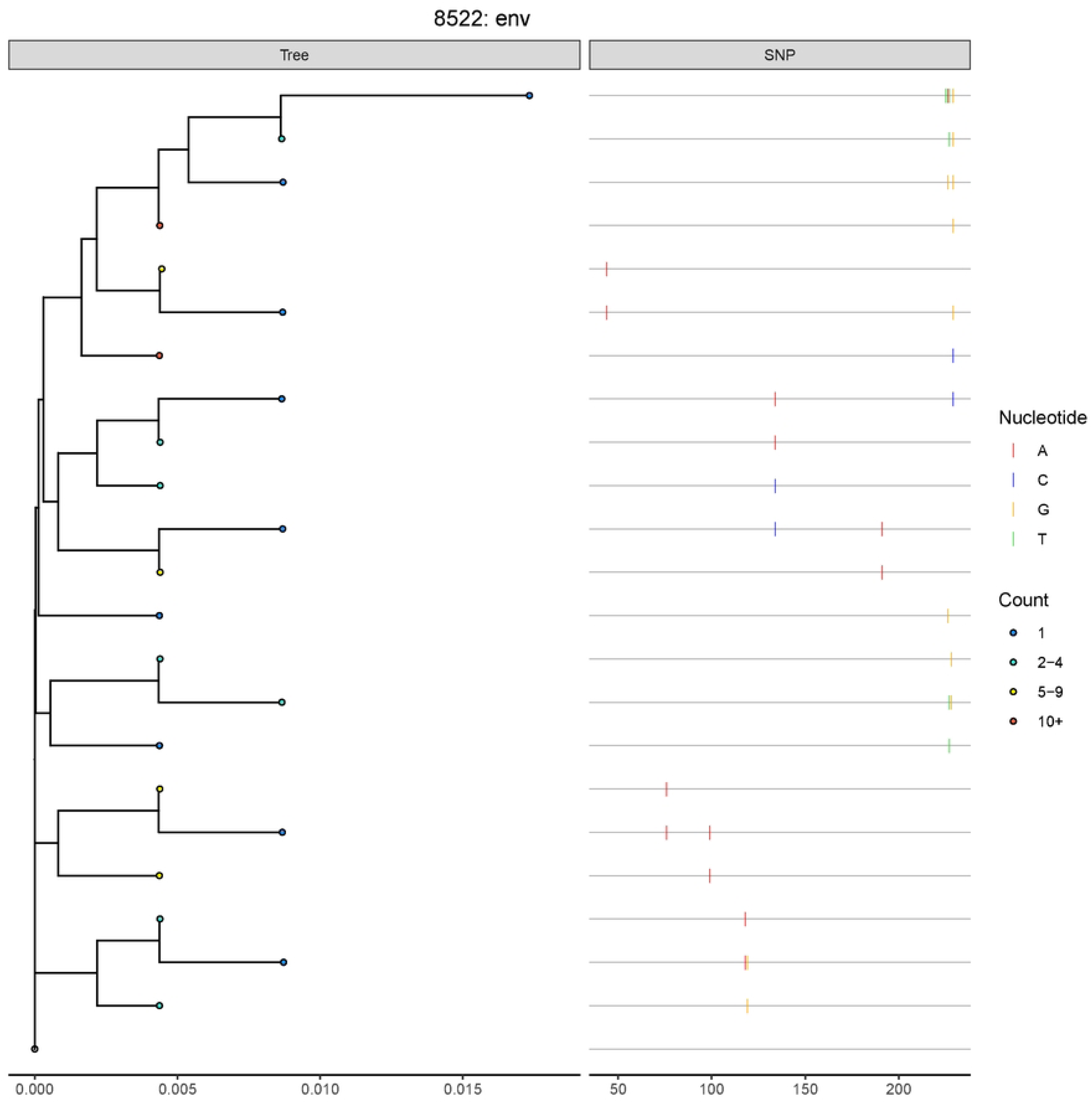
Phylogenetic reconstruction of TF viral quasi-species population during Fiebig stages II-III after filtering potential APOBEC3G/F mutations. **Trees were reconstructed from** viral genomes in the GR-AR filtered dataset where ≥2 genomes were included thereby excluding singlets (*i.e.*, viral genomes that differed from the TF by 1 nt and found only once). The tree was rooted on the TF for *pol* and *env*. Neighbor-joining phylogenetic trees were reconstructed and are plotted to the level of diversity (where 0.004 units ∼ 1nt) with respect to that participant’s inferred single TF virus (serving as the root). The nodes at the leaves are colored by the number of collapsed viral genomes with 1 (dark blue), 2-4 (light blue), 5-9 (yellow), and 10+ (red). In alignment to the phylogenies, a SNP matrix (highlighter plot) is provided where changes to the inferred single TF virus are noted as: A (red), G (orange), C (blue), and T (green). The nucleotide position is relative to the absolute sequenced region. At the top of each tree is the PID and region.

**Figure S7.**
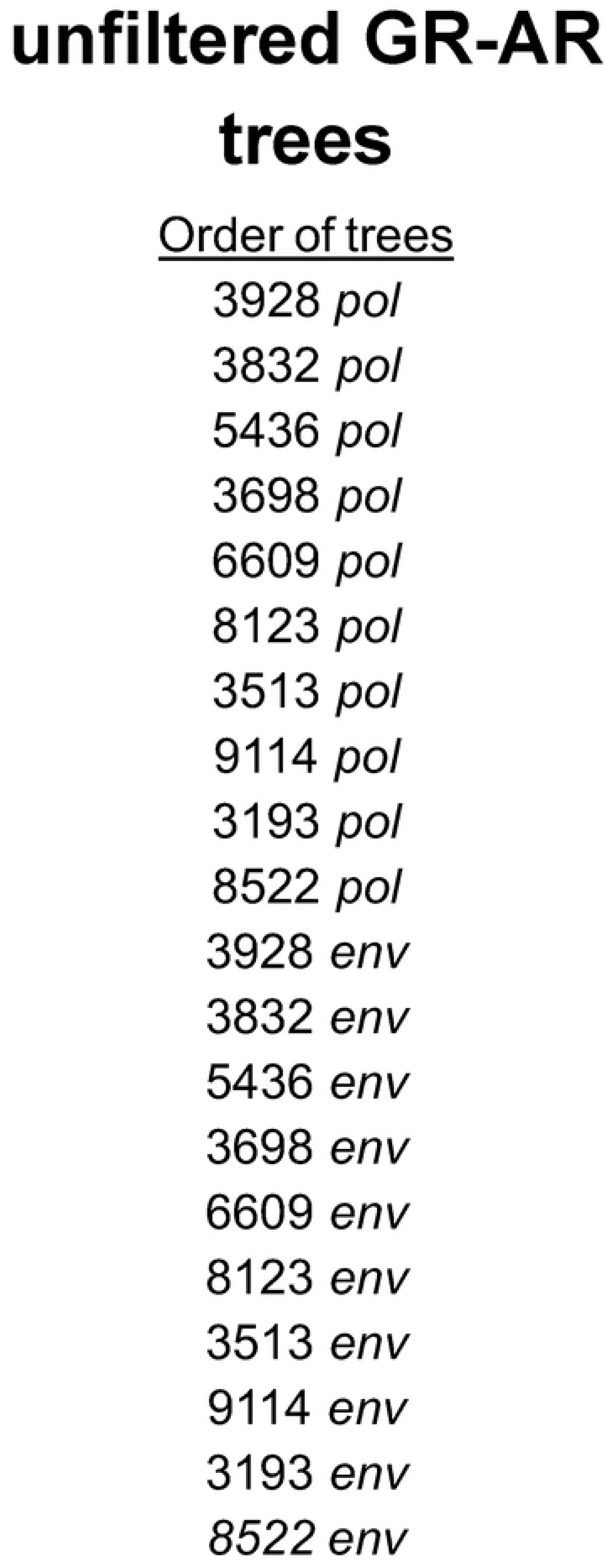
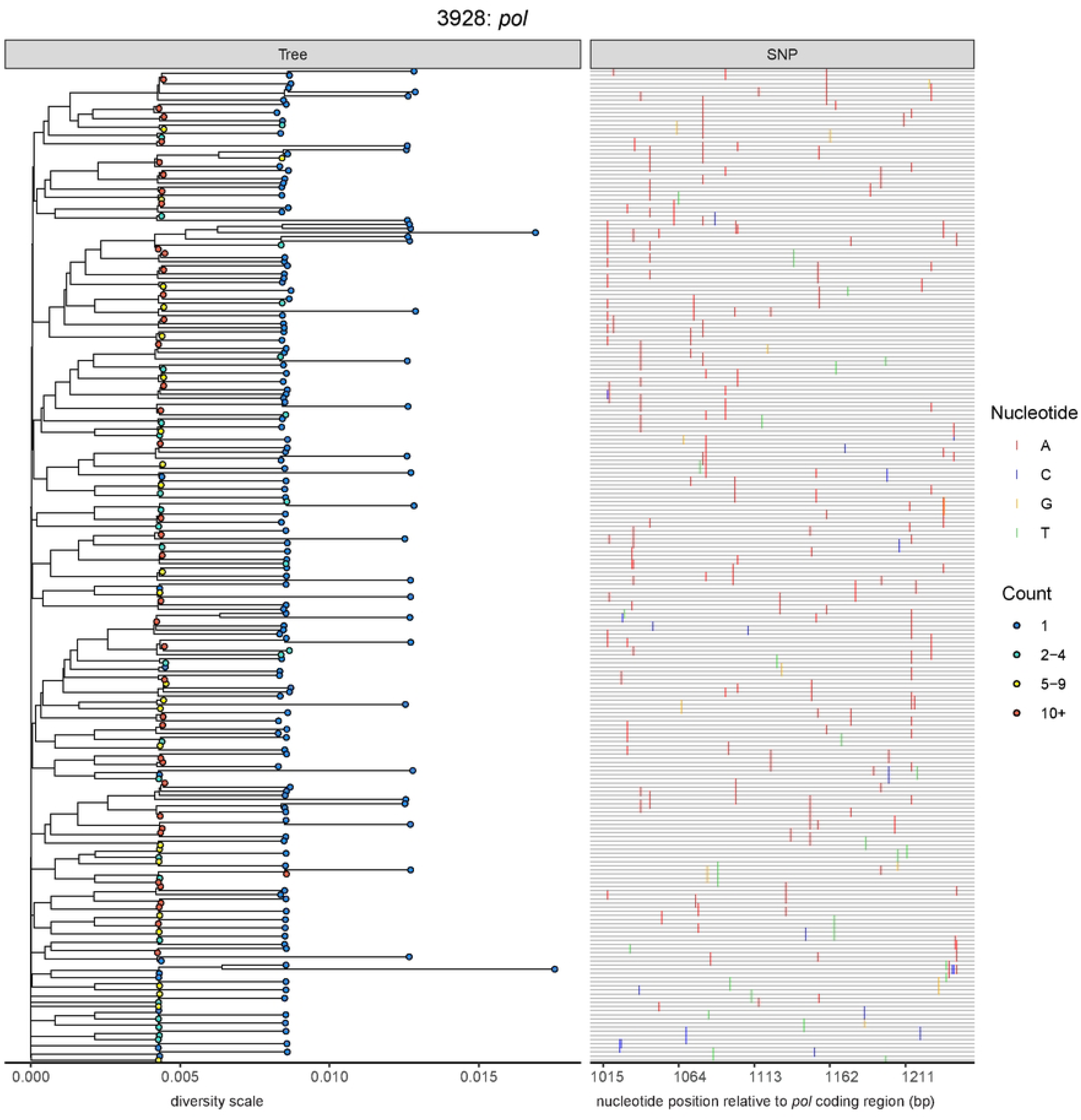
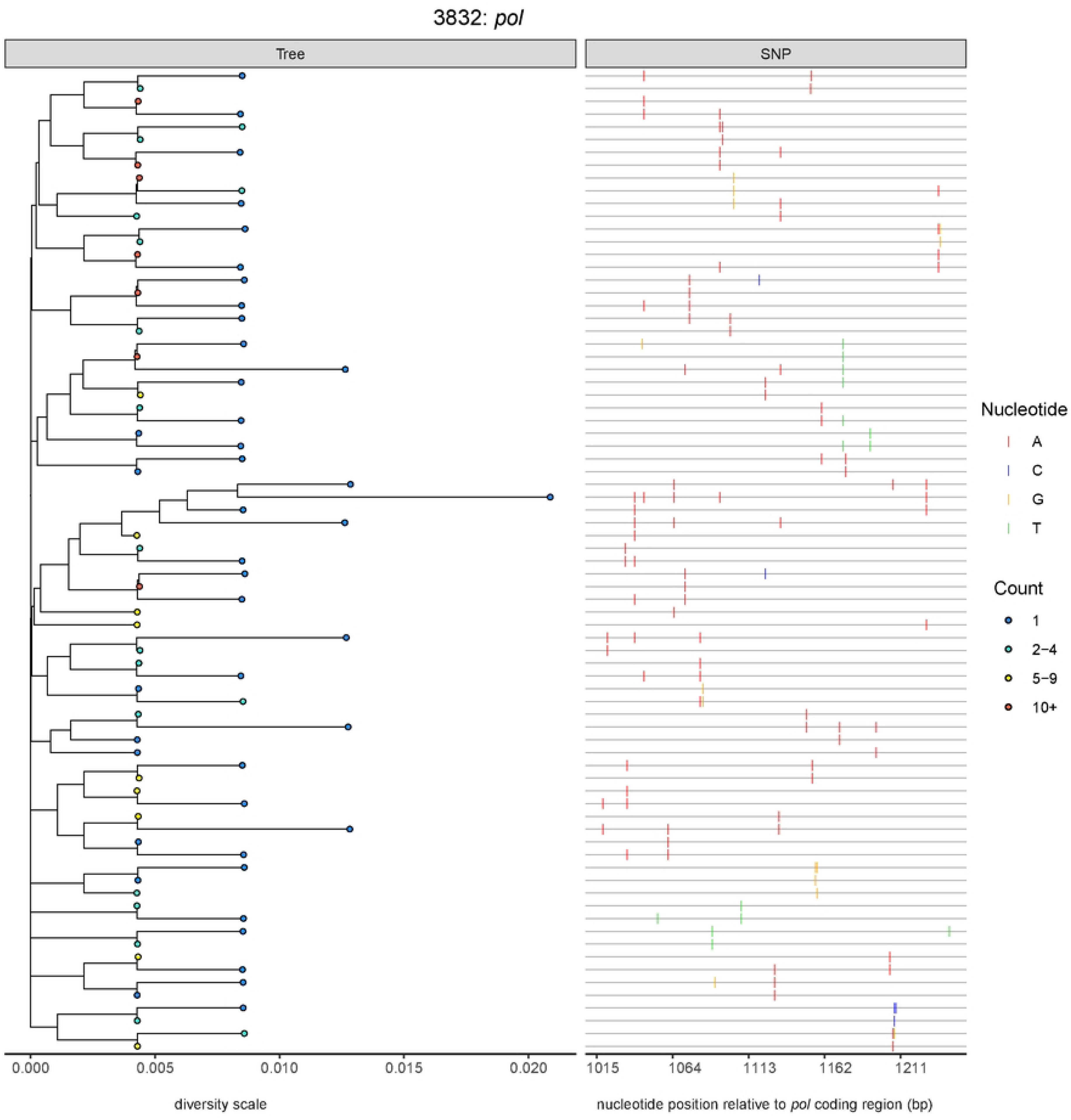
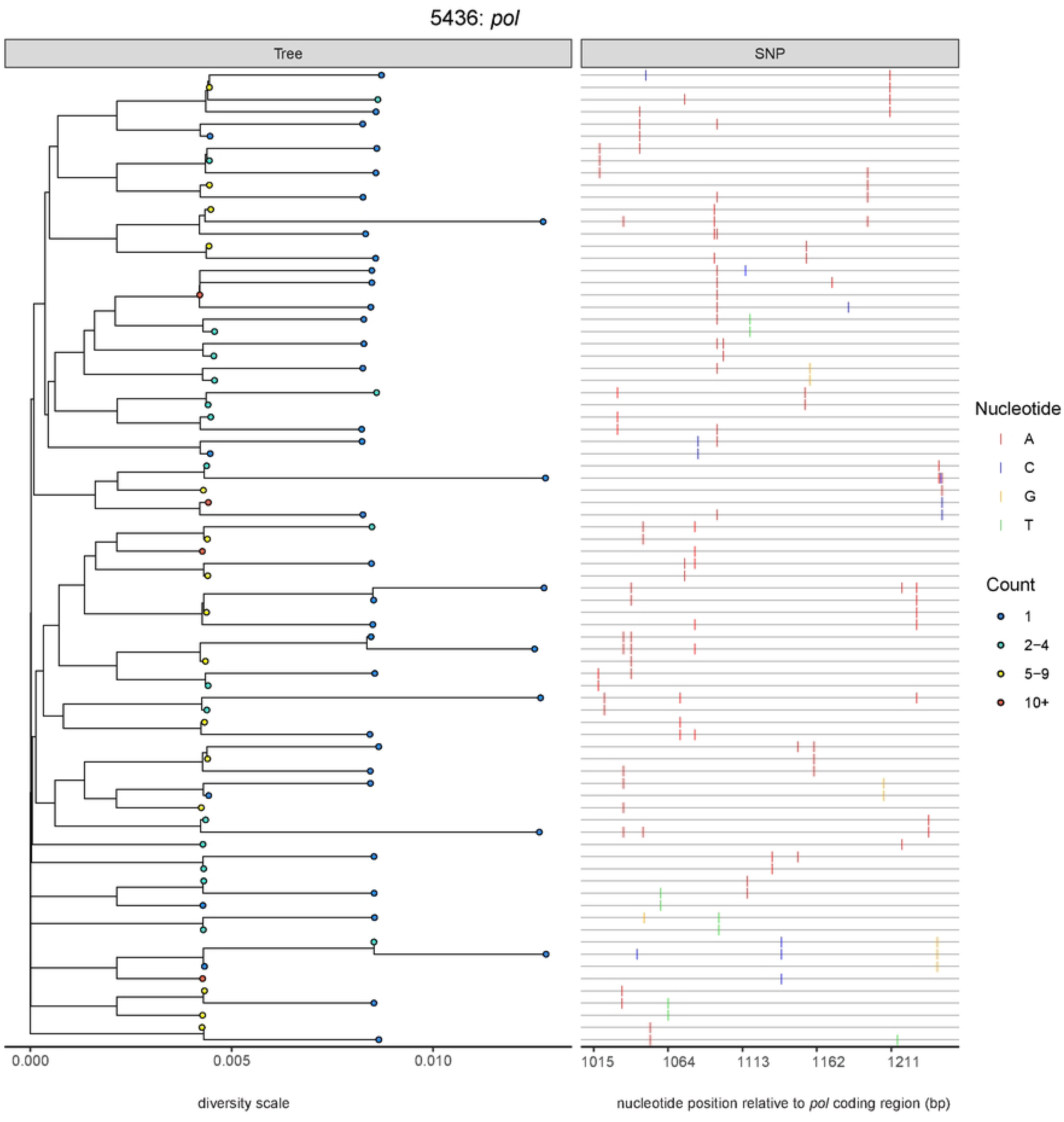
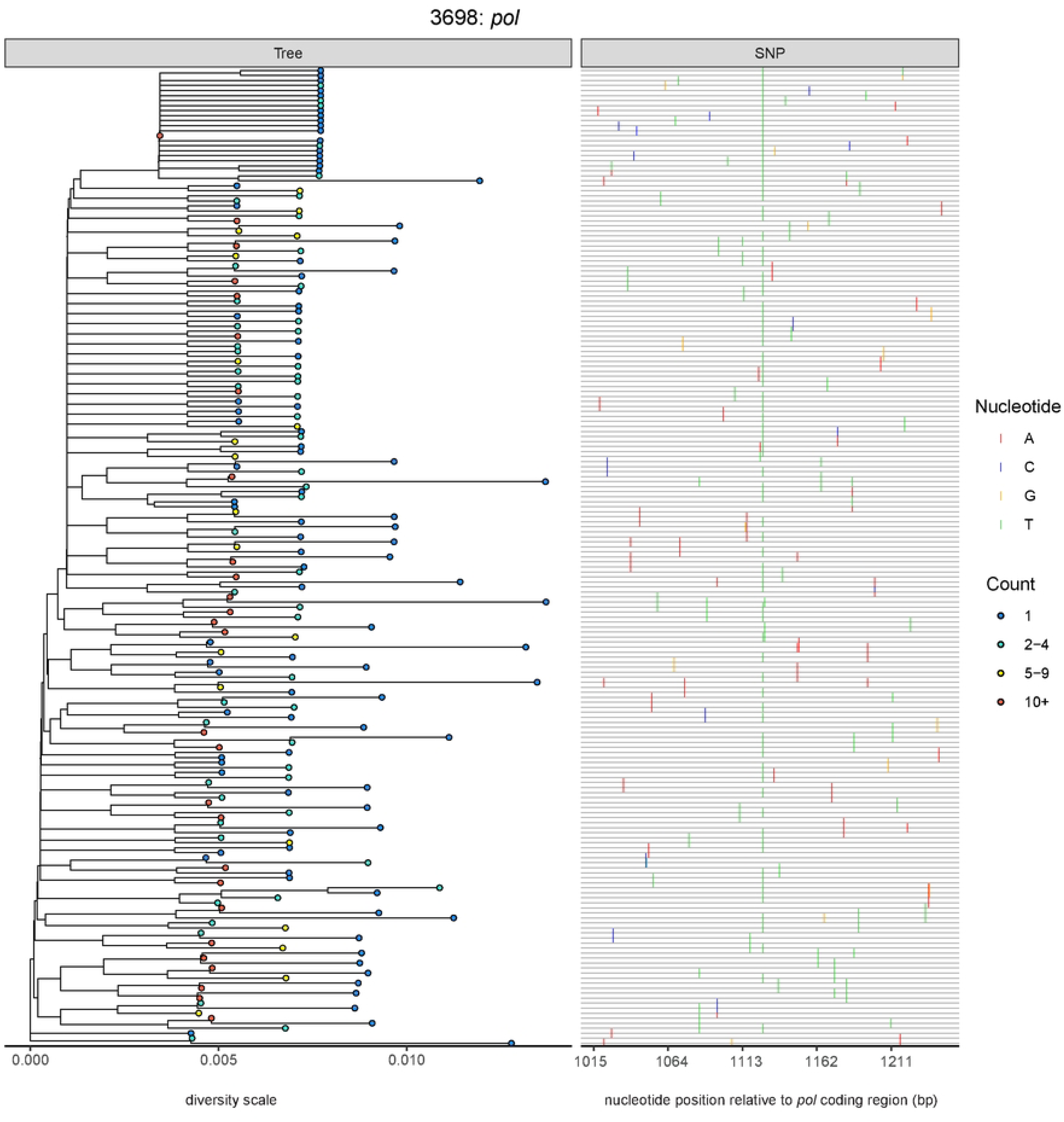
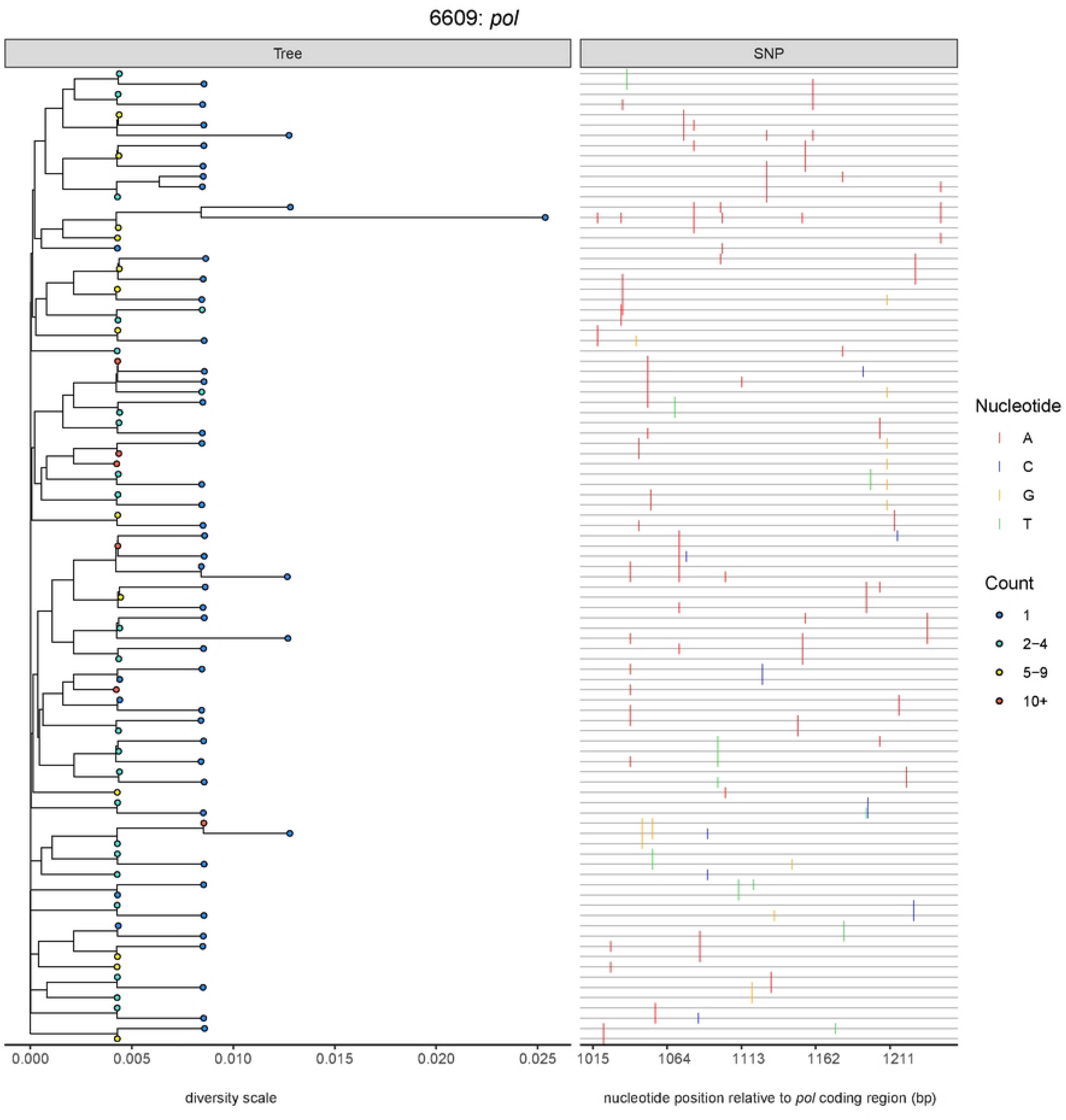
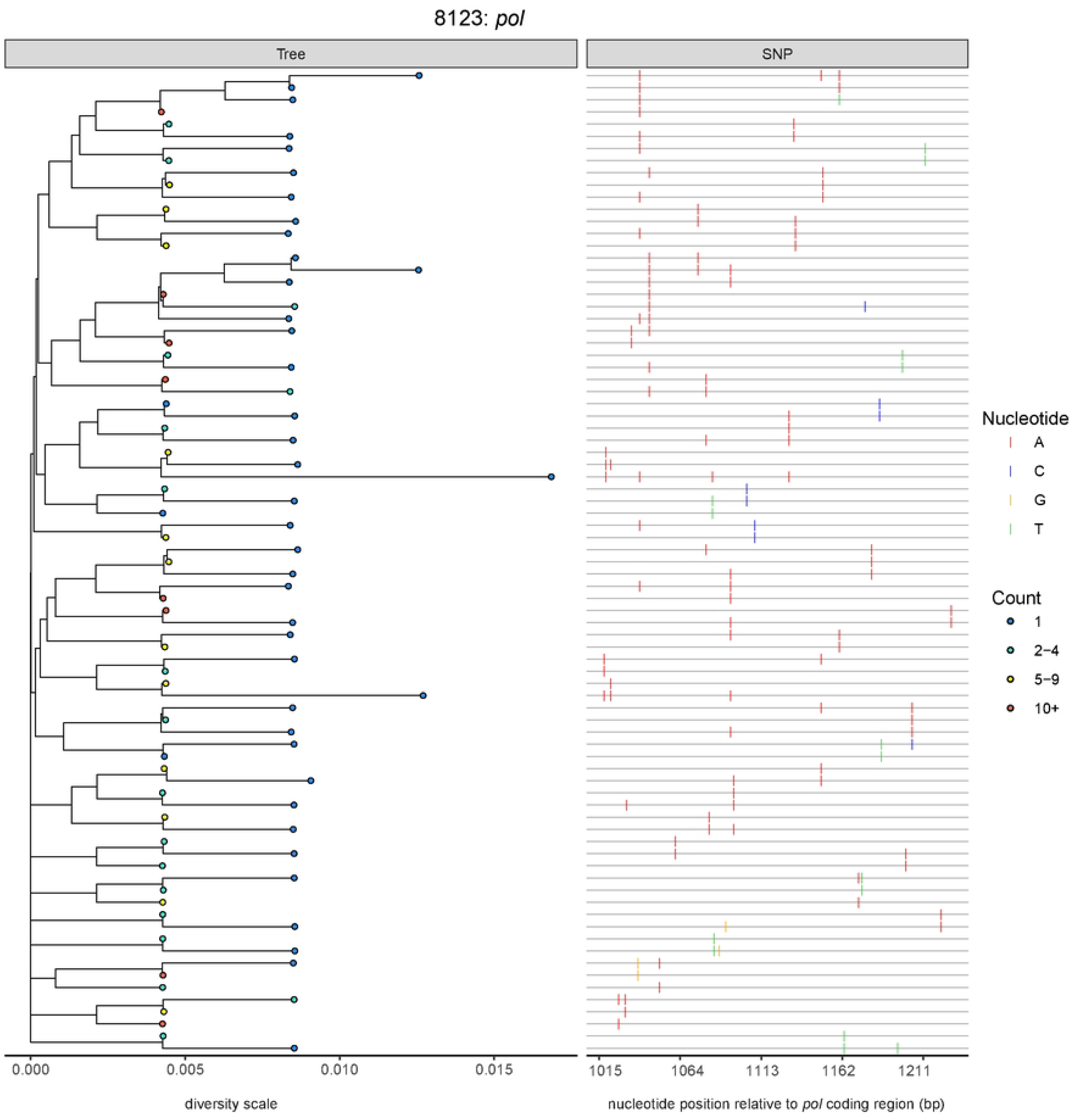
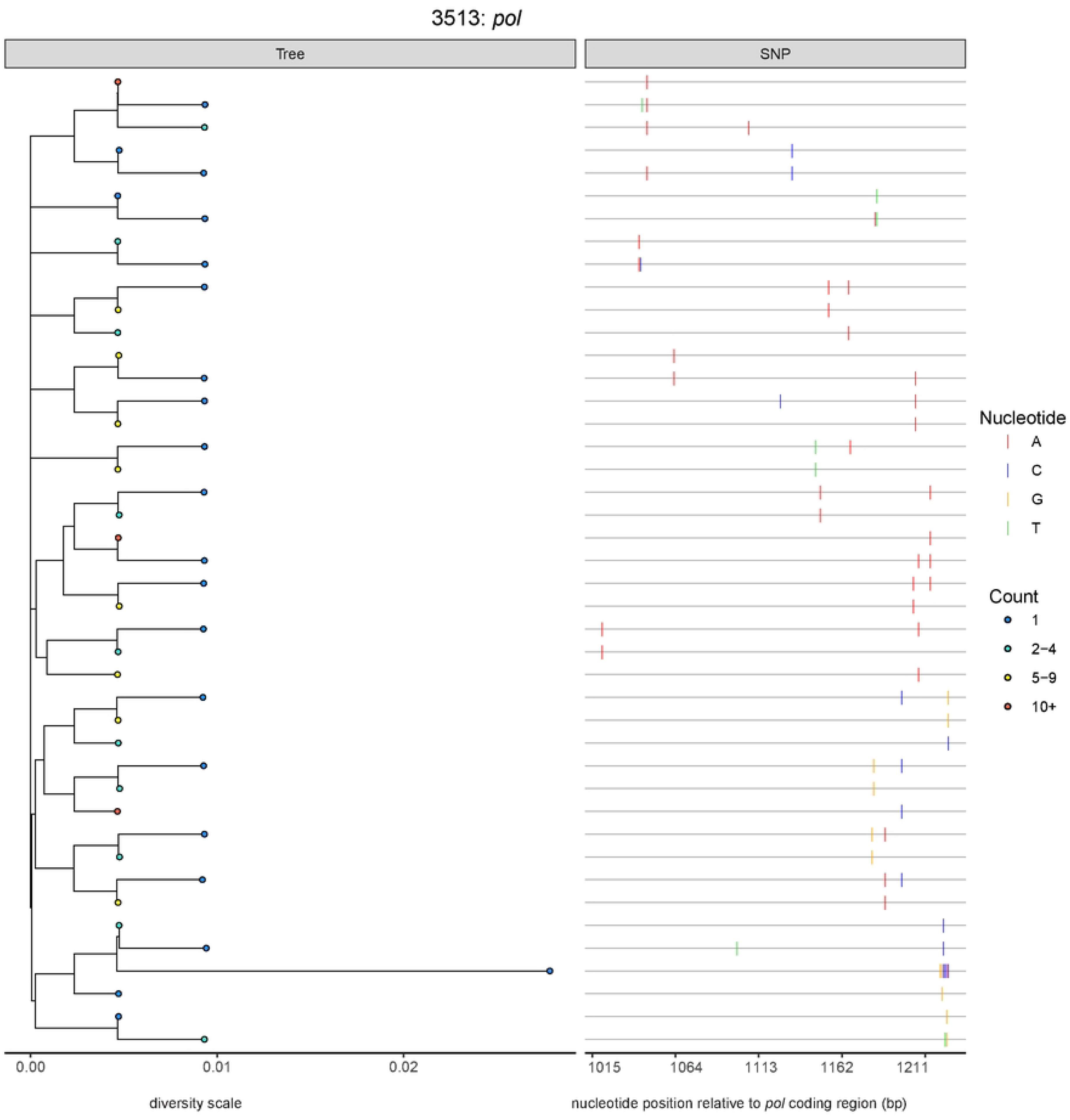
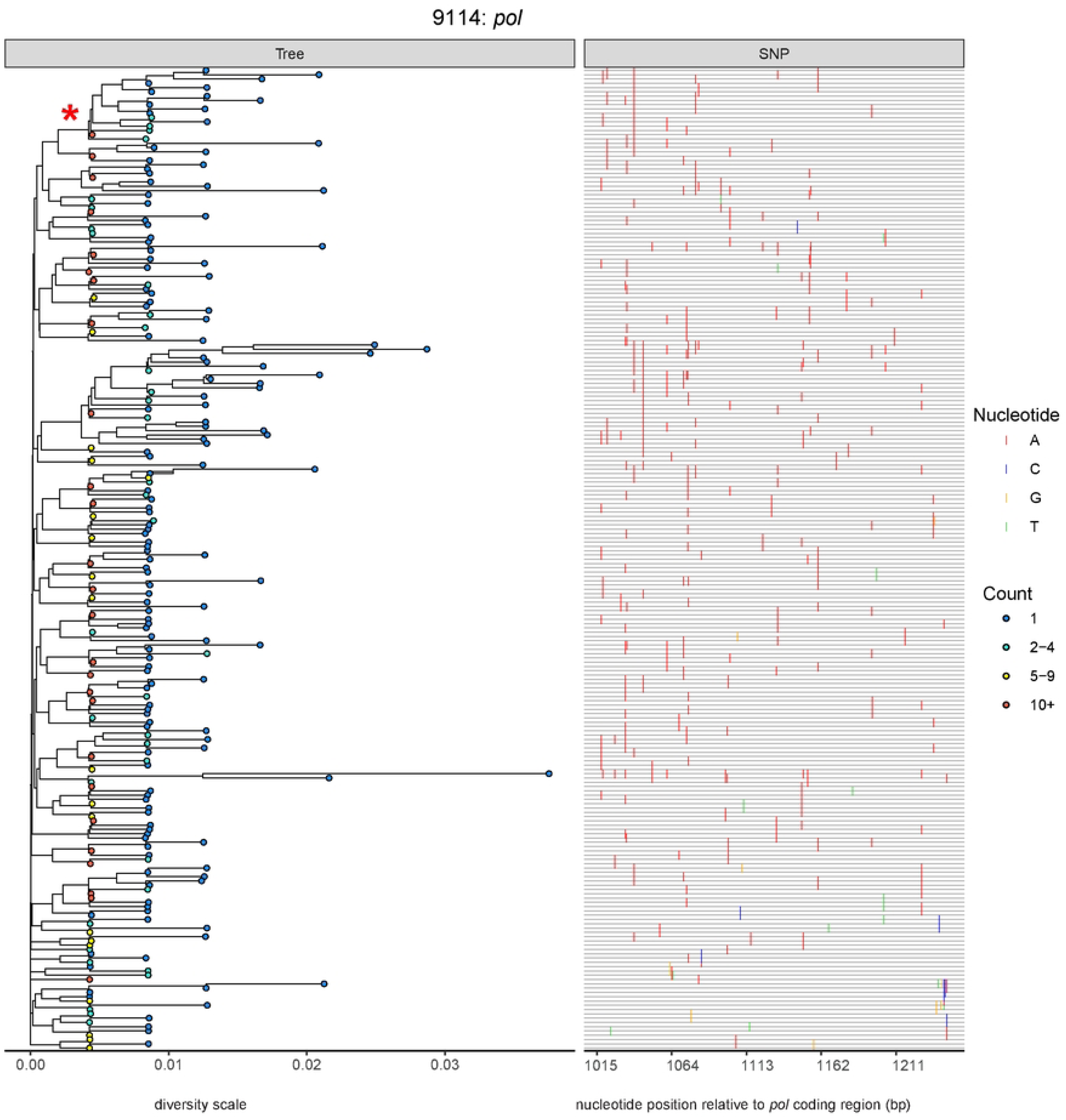
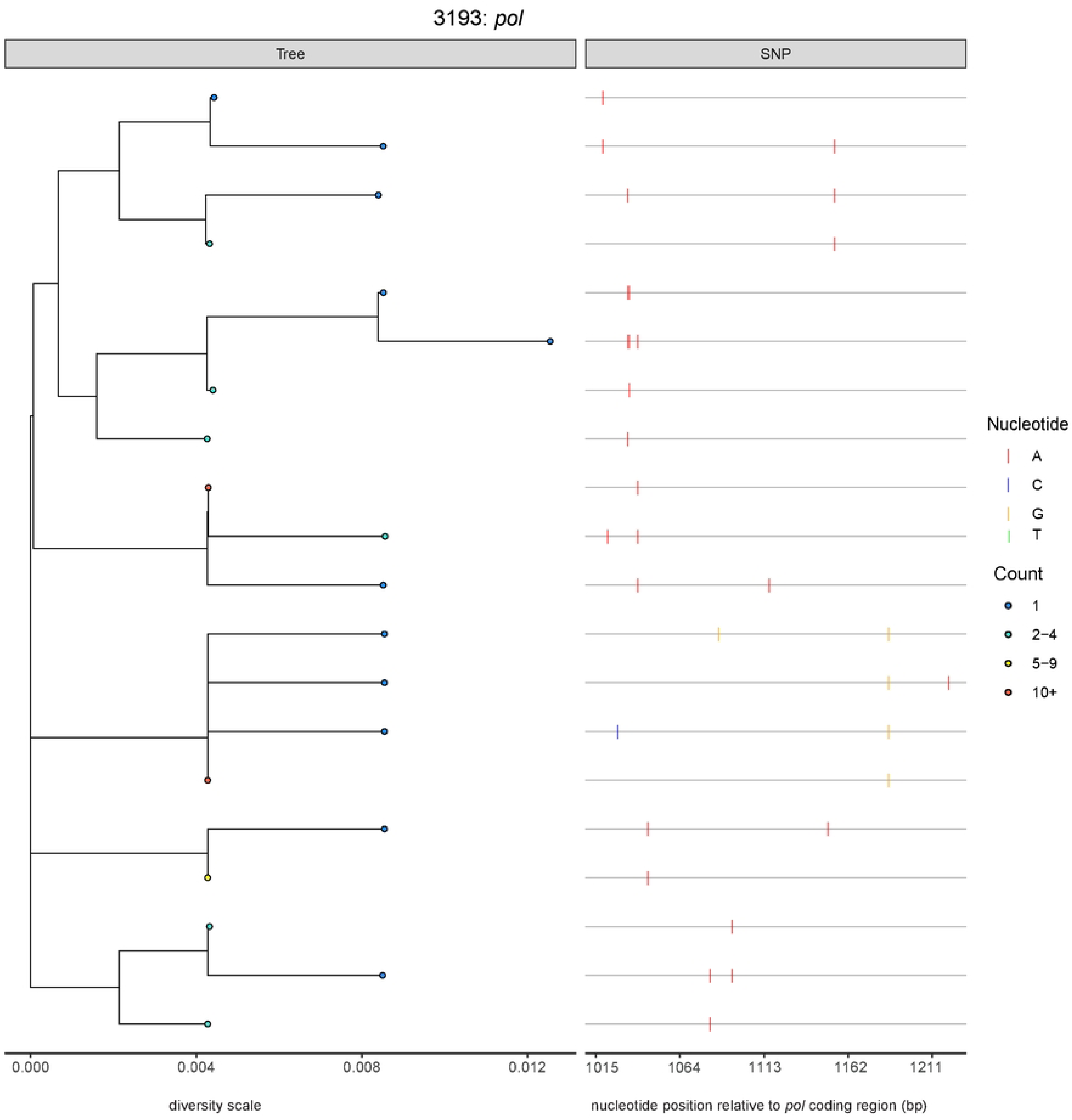
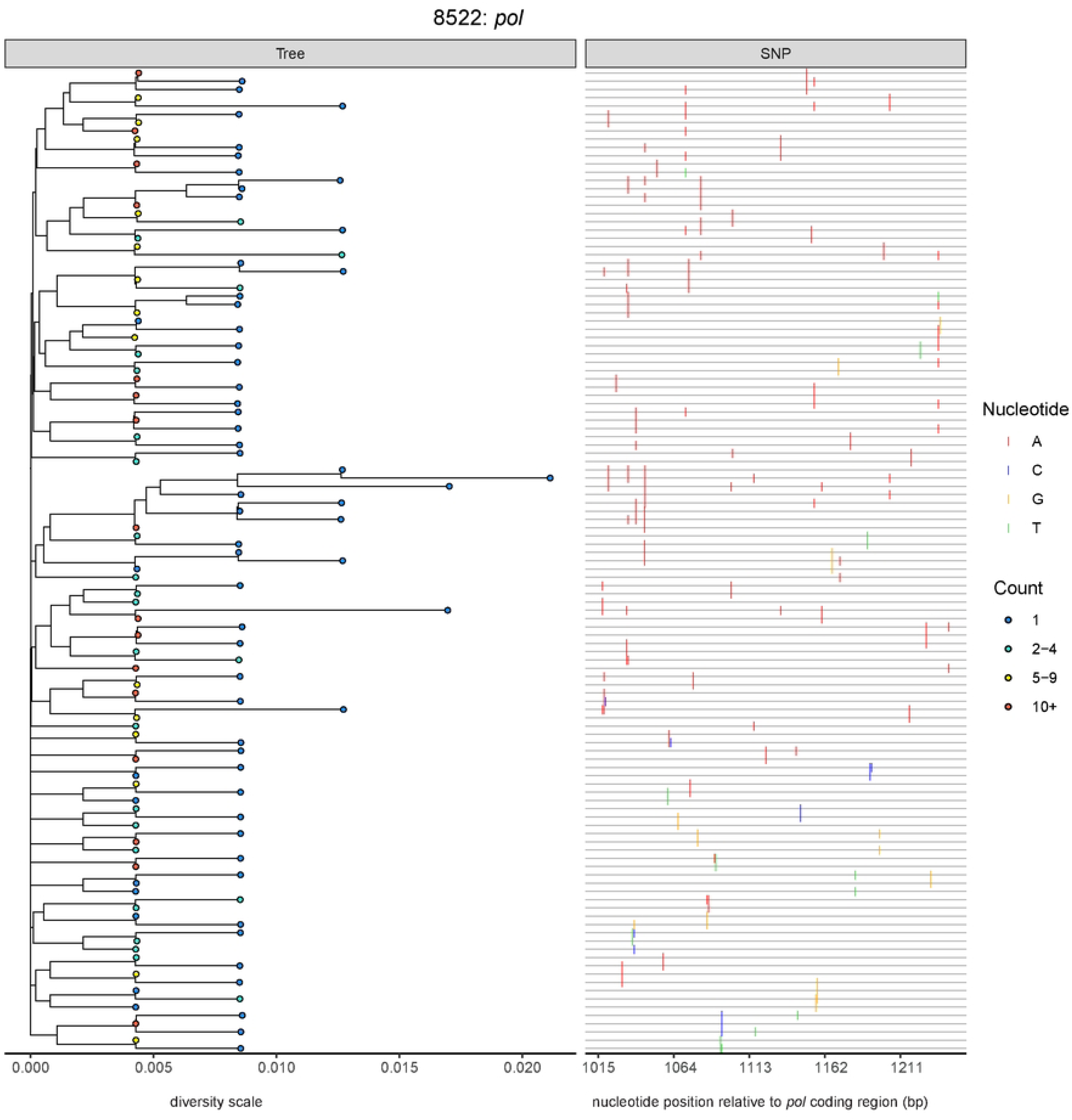
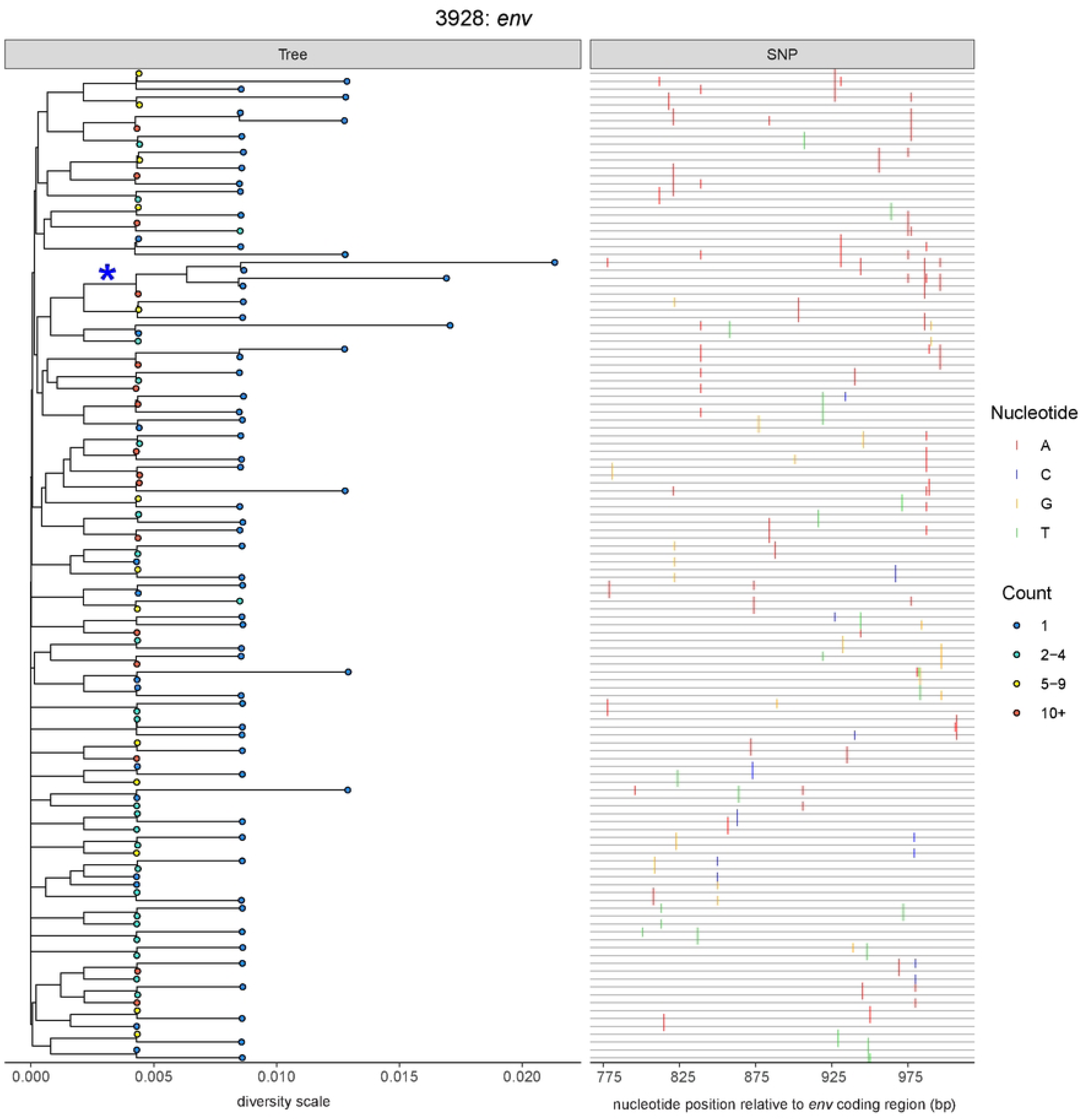
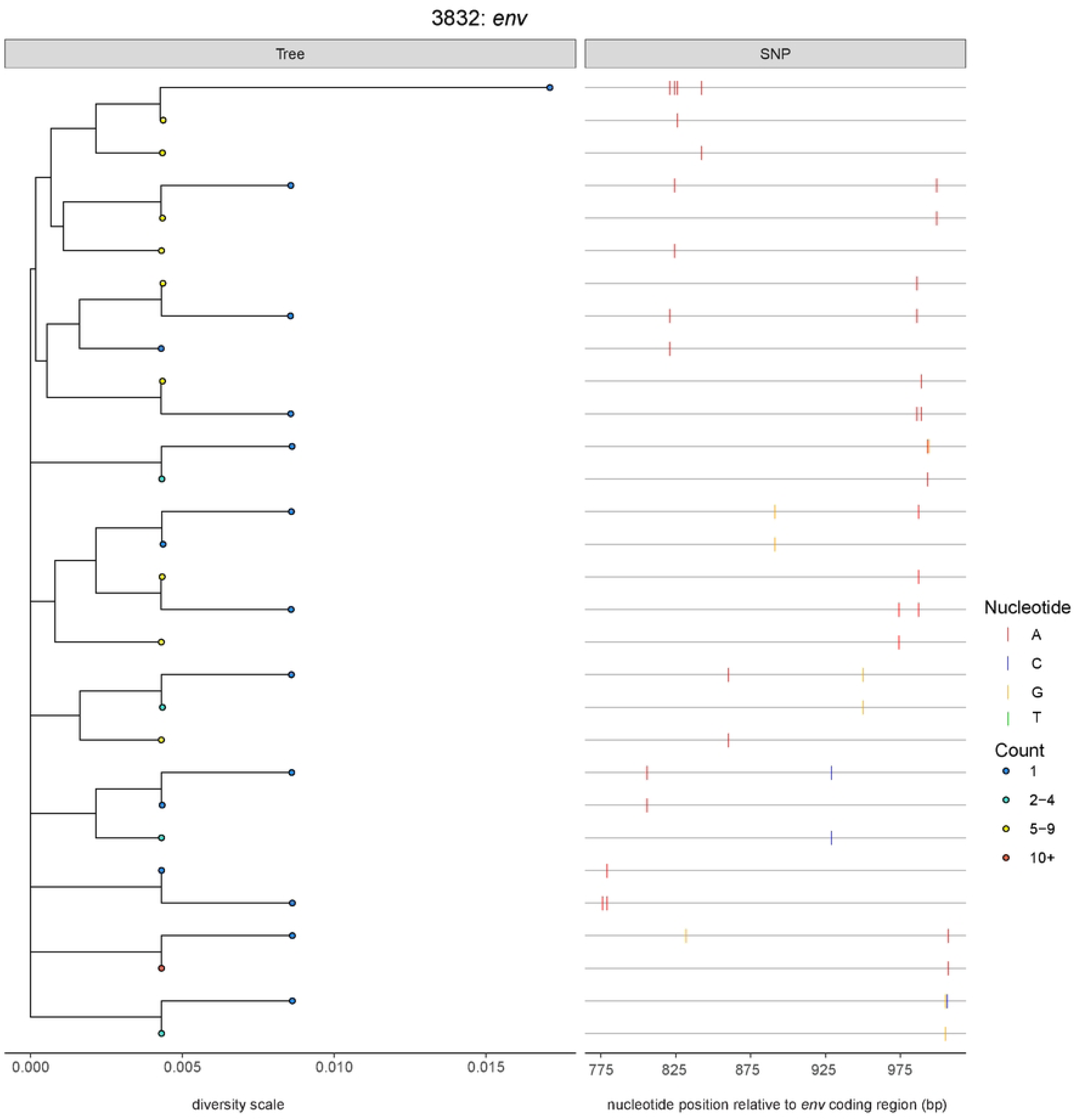
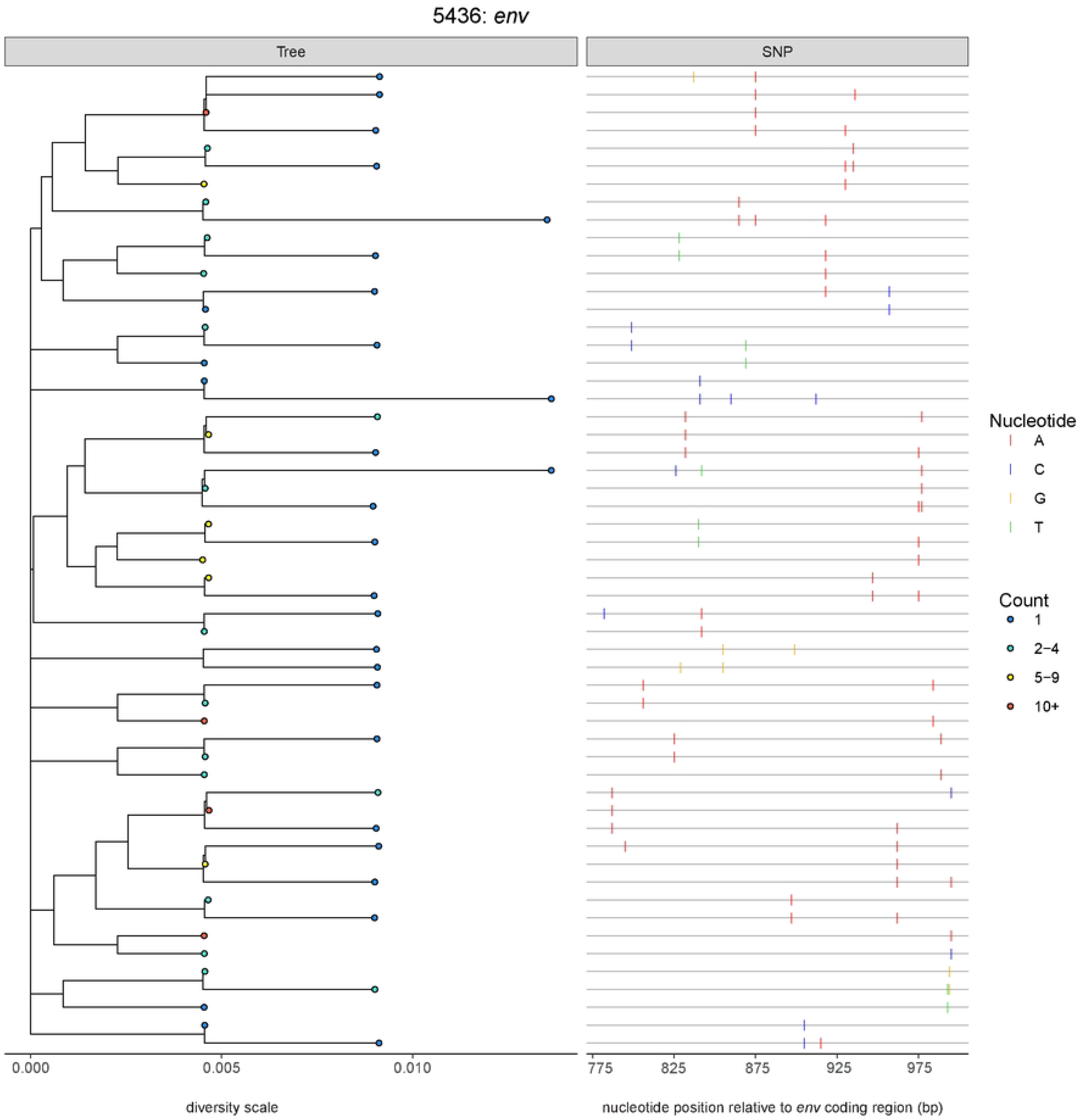
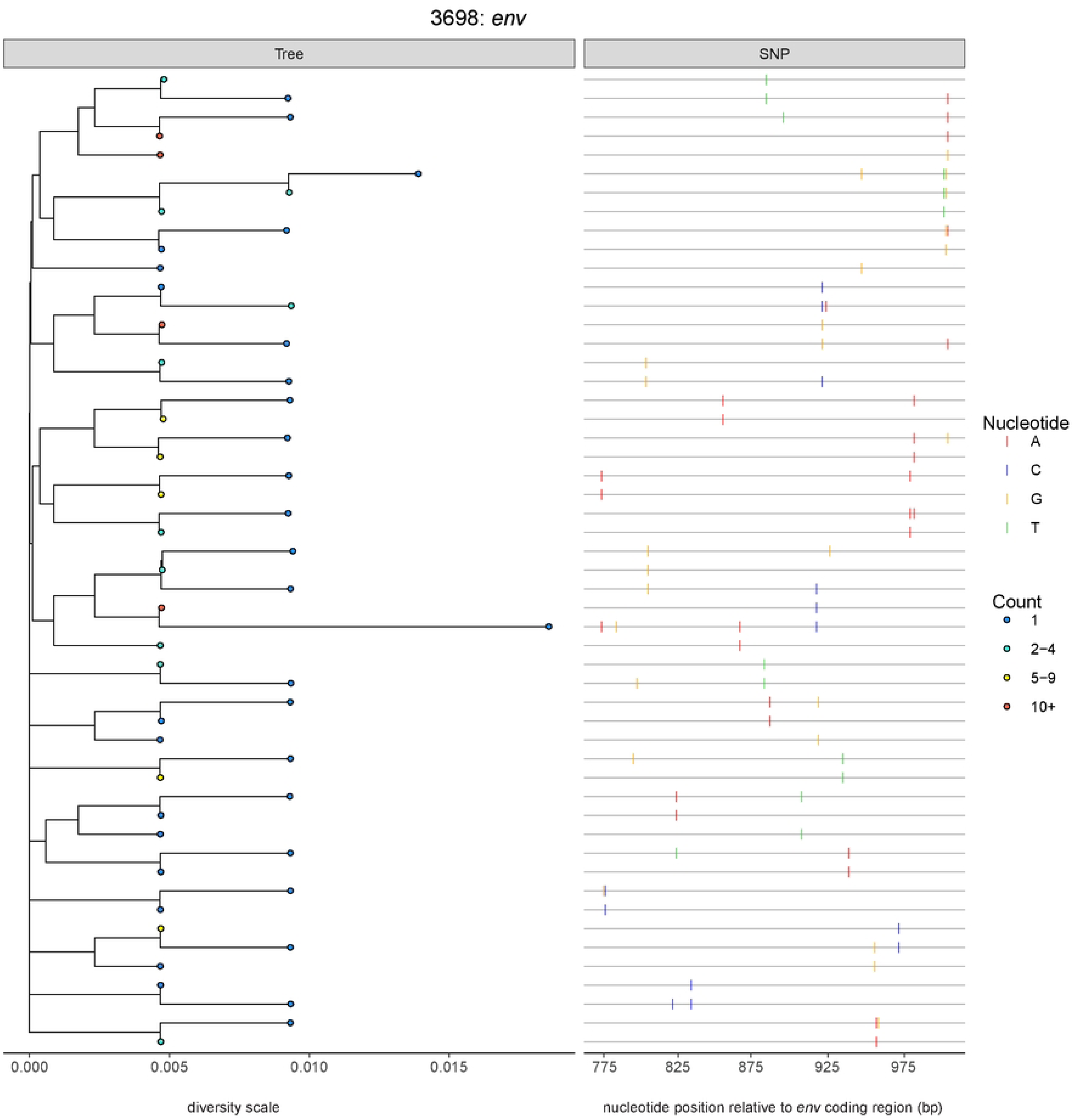
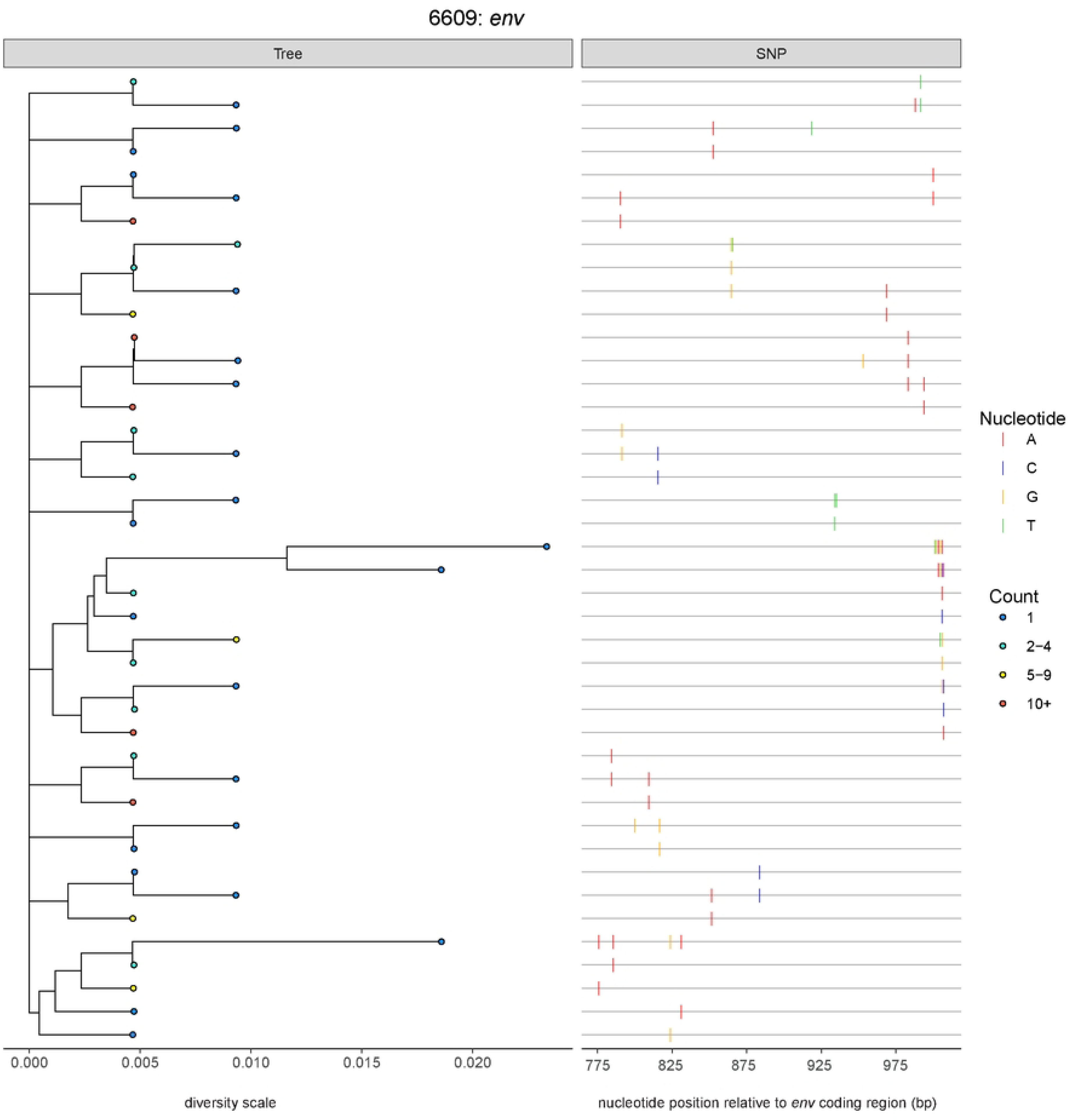
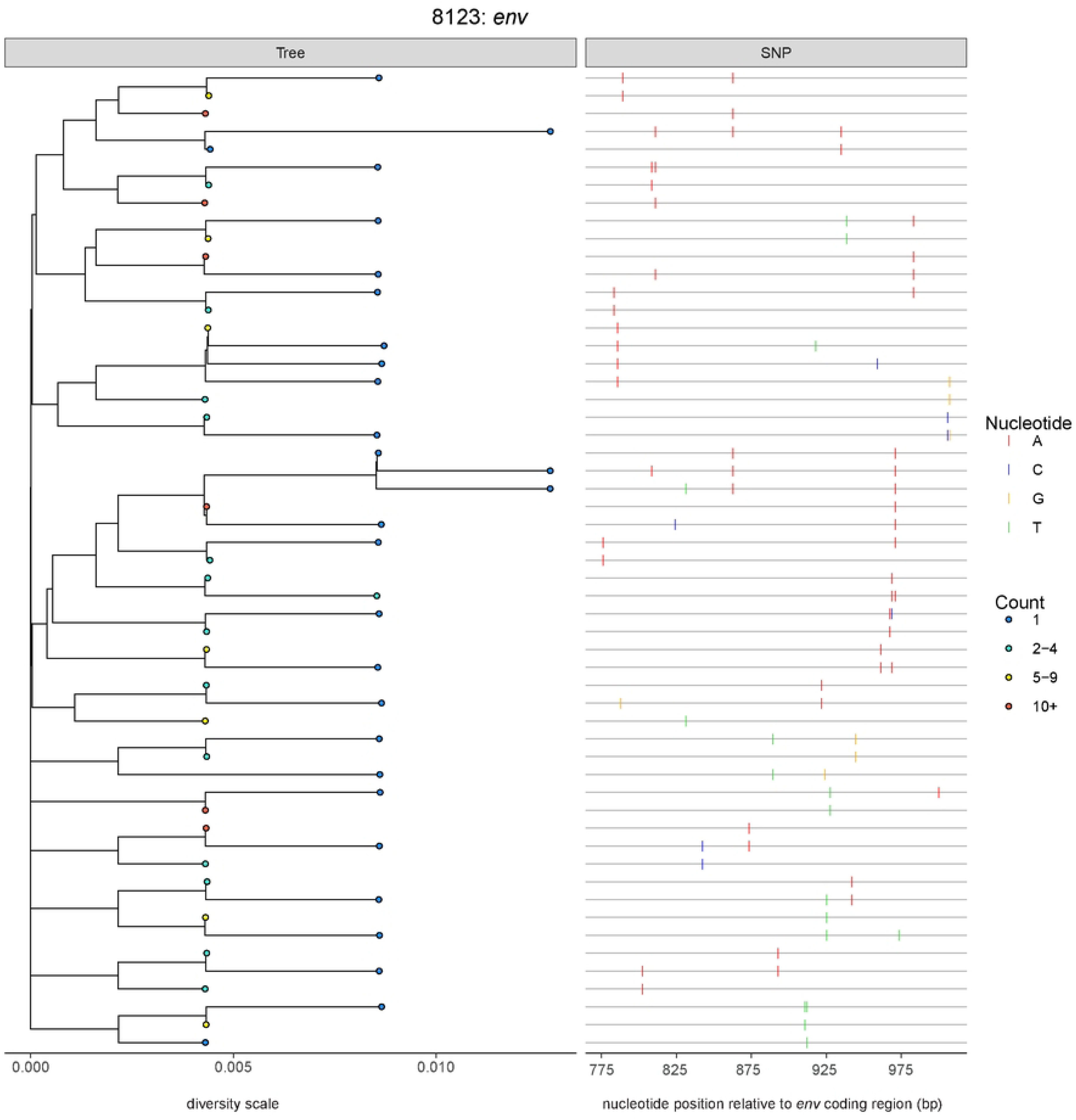
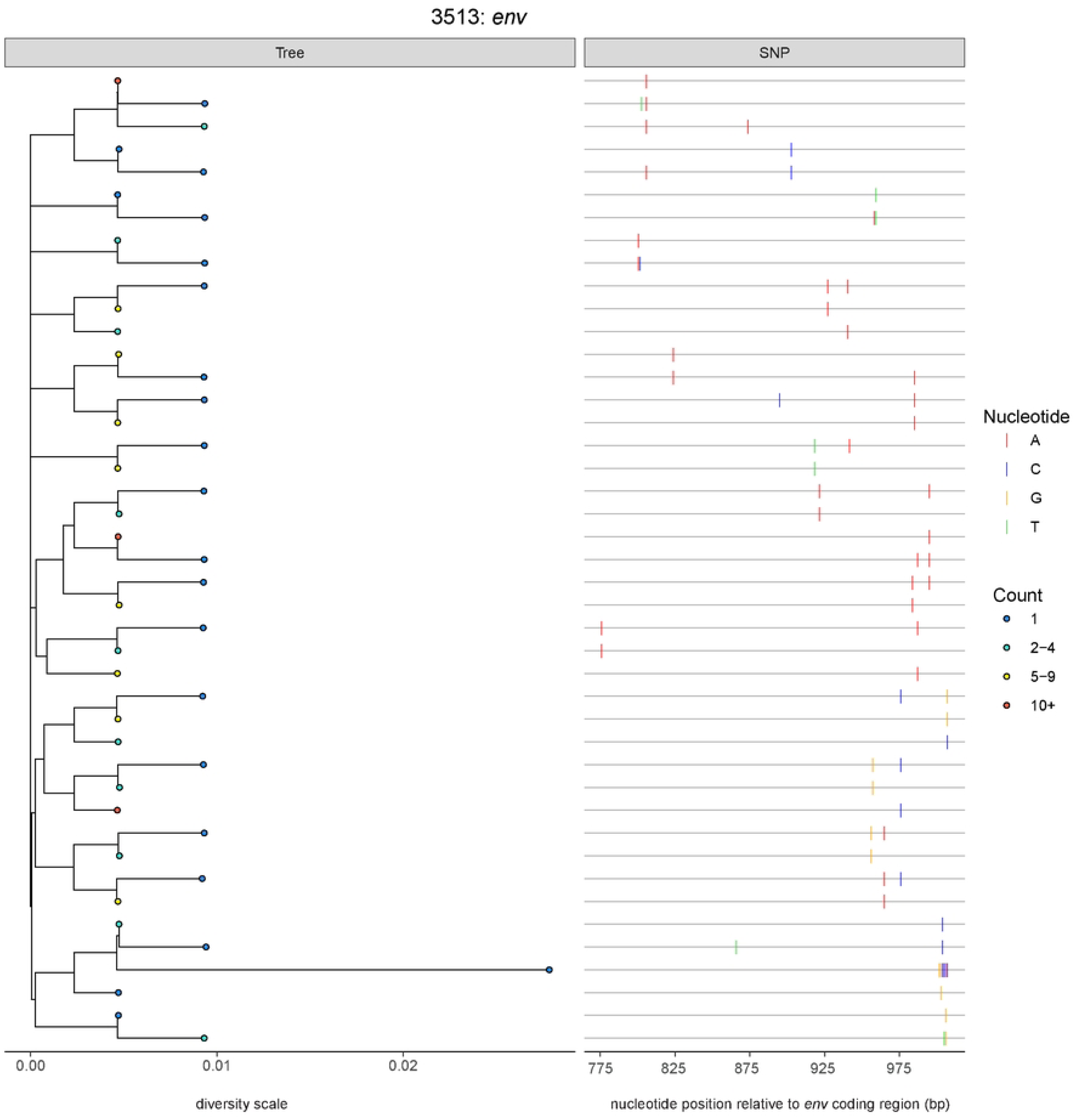
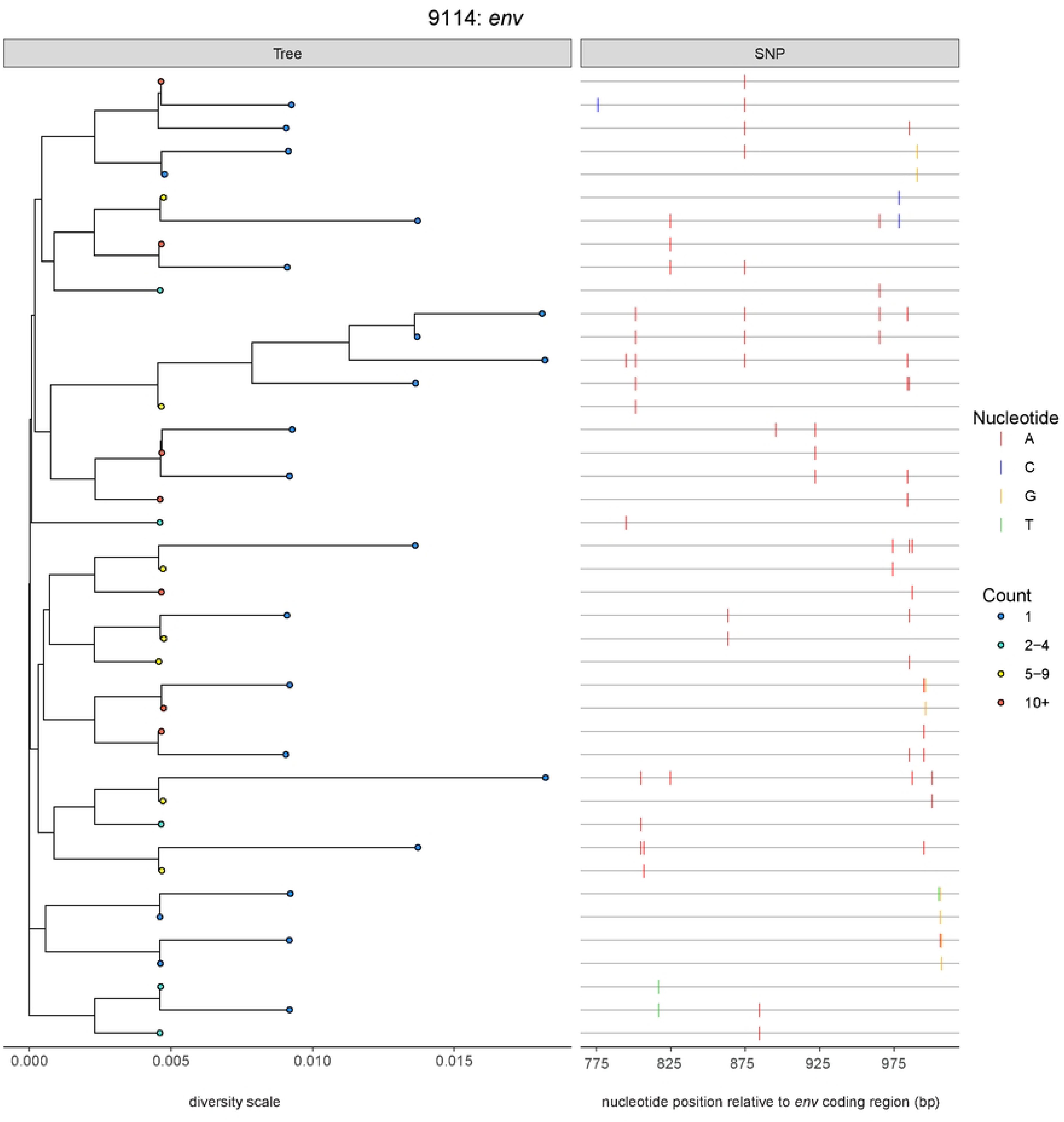
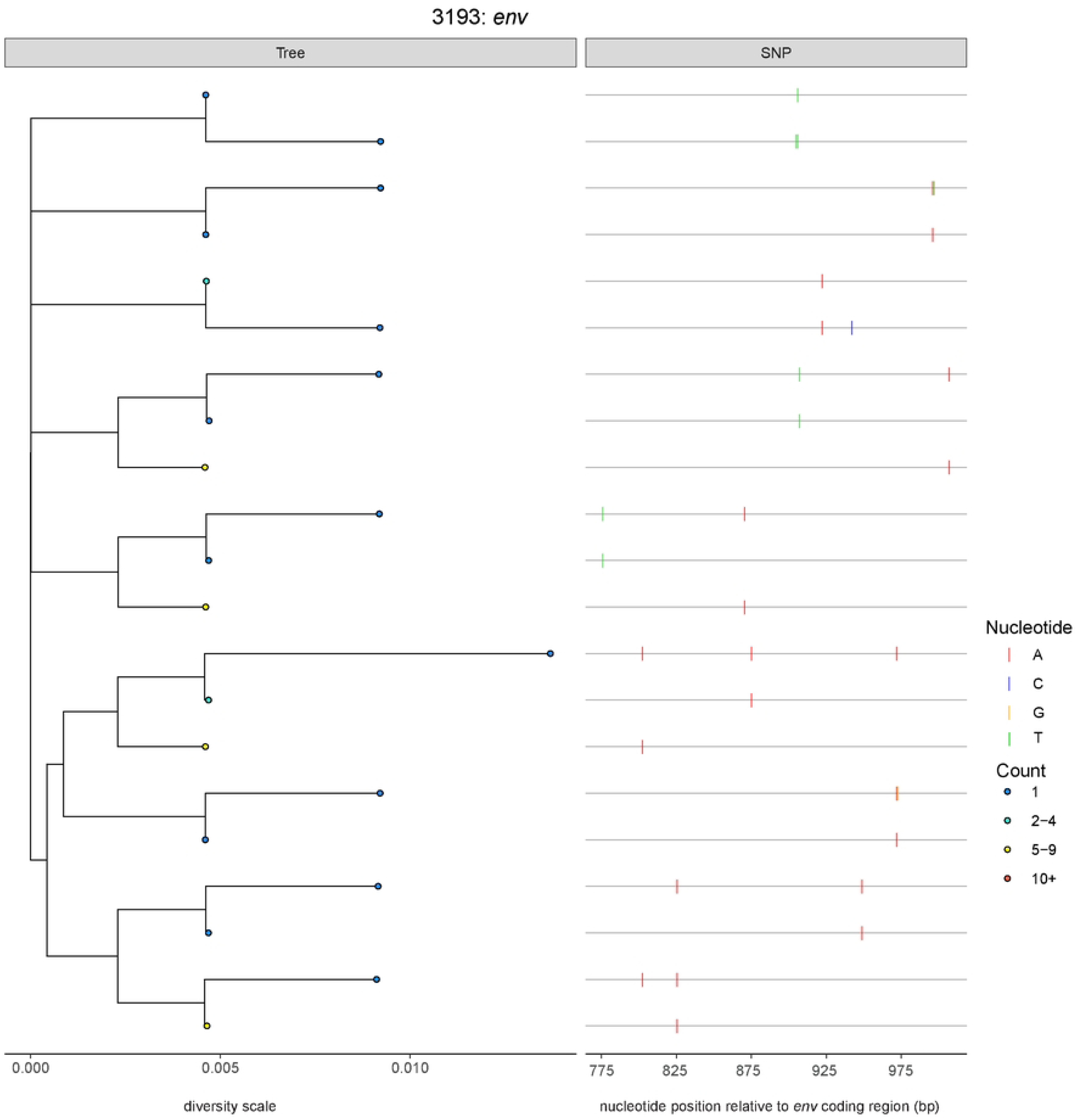
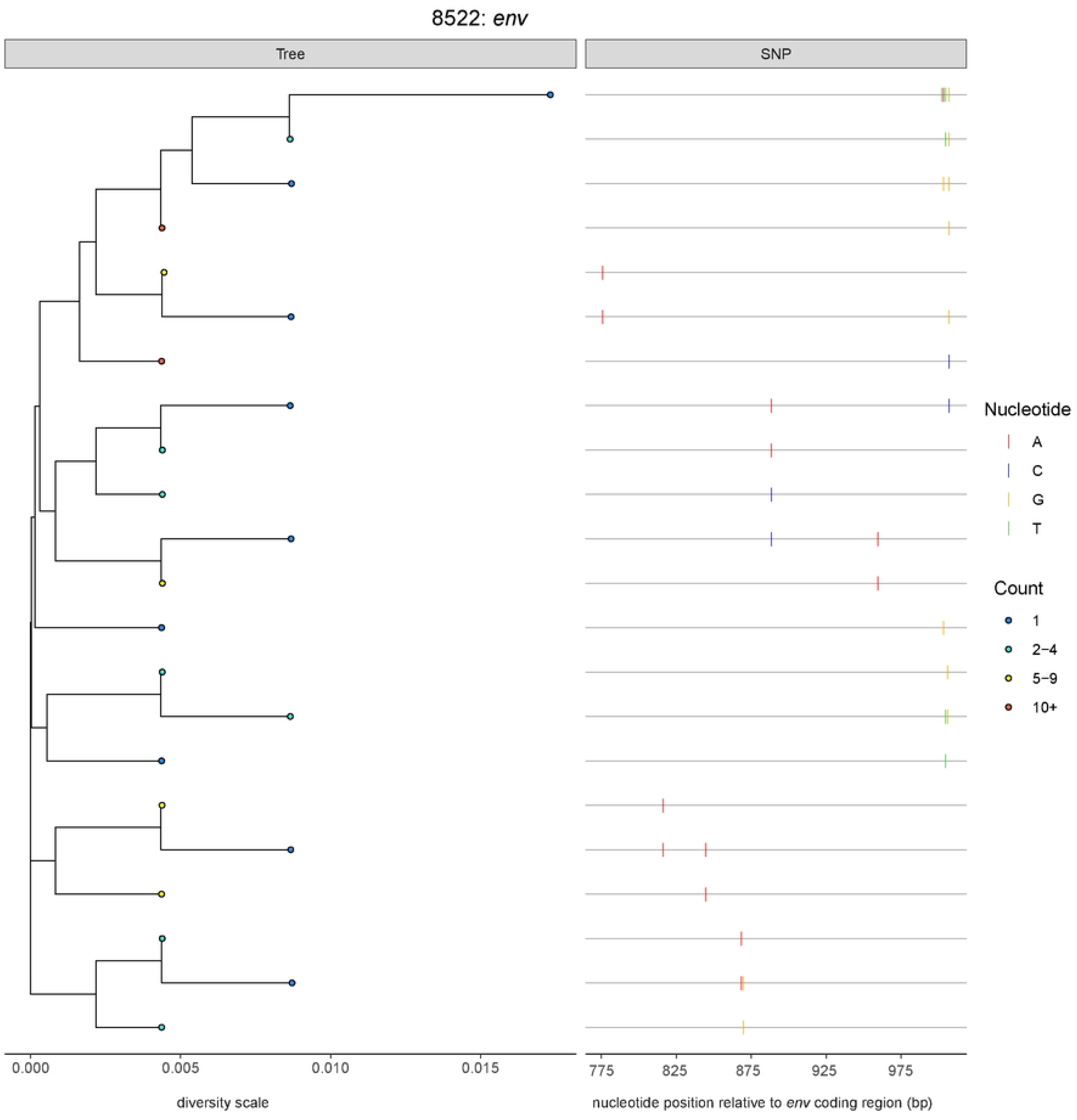
Phylogenetic reconstruction of TF viral quasi-species population during Fiebig stages II-III including APOBEC3F/G mutations. Viral genomes that were generated from ≥2 were included thereby excluding singlets (*i.e.*, viral genomes that differed from the transmitted/founder by 1nt and found only once) where the tree was rooted on the TF for *pol* and *env*. Neighbor-joining phylogenetic trees were reconstructed and are plotted to the level of diversity with respect to that participant’s inferred single TF virus (serving as the root). The nodes at the leaves are colored by the number of collapsed viral genomes with 1 (dark blue), 2-4 (light blue), 5-9 (yellow), and 10+ (red). In alignment to the phylogenies, a SNP matrix (highlighter plot) is provided where changes to the inferred single TF virus are noted as: A (red), G (orange), C (blue), and T (green). The nucleotide position is relative to the coding region sequenced in HXB2 coordinates.

**Figure S8.**
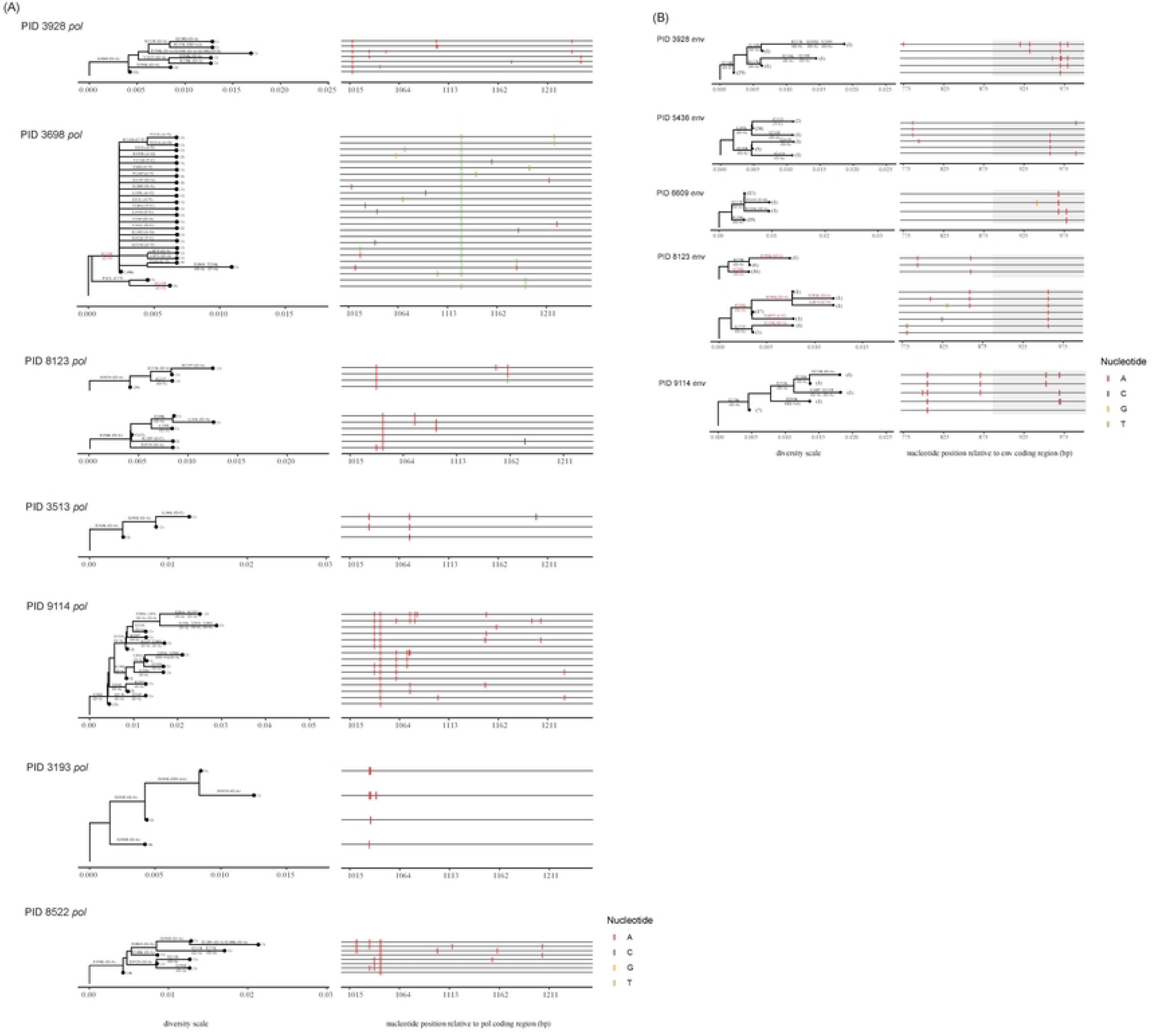
Extracted pseudo-trees of select branches that illustrate likely effects of APOBEC3G/F activity or ‘Founder Effects’. Selected inferred single TF virus participants for (**A**) *pol* and (**B**) *env*. Neighbor-joining phylogenetic trees were reconstructed and selected branches were excised for further analysis. The branches are likely artificial due to extensive APOBEC3G/F activity with the accumulation of multiple G>A mutations within GR>AR motifs. In select pseudo-trees, mutations in red indicate “reversions” to the consensus CRF01_AE. The excised branches are plotted according to the Hamming distance with respect to that participant’s inferred single TF virus (serving as the root). Labeled on branches are the amino acid changes (both synonymous and non-synonymous) reported with the associated nucleotide change. The leaves of the trees have the number of viral genomes in parentheses. In PID 3698 *pol*, synonymous C>T change that would be considered a reversion to the consensus CRF01_AE demonstrates a potential ‘Founder Effect”. In alignment to the phylogenies, an SNP matrix (highlighter plot) is provided where changes to the inferred single TF virus are noted as: A (red), G (orange), C (blue), and T (green). The nucleotide position is relative to the coding region sequenced in HXB2 coordinates. The listed mutations on the branches are relative to the amino acid change in HXB2 coordinates.

**Figure S9.**
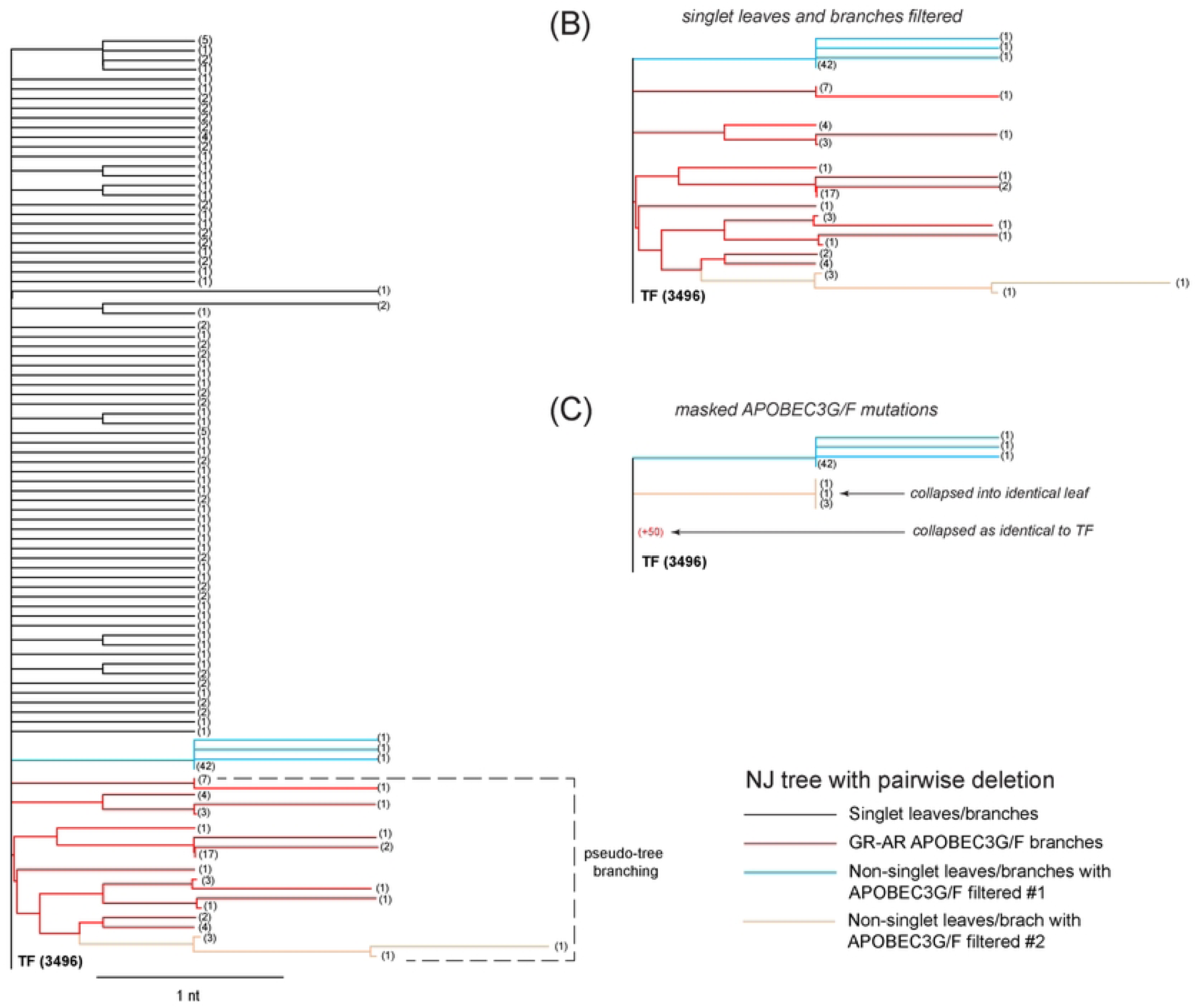
Representative tree illustrating pseudo-tree topology caused by APOBEC3G/F mutations. PID 3193 *pol* neighbor-joining tree with pairwise deletion for gap handling. (**A**) Complete tree with all unique viral genomes included. Majority of genomes are singlet leaves/branches (black). Pseudo-tree branching is highlighted within the overall topology (dotted bracket). (**B**) Tree topology after filtering singlet leaves/branches. Equivalent tree can be found in **Figure S7**. (**C**) Tree topology after masking all GR-AR mutations that could be due to APOBEC3G/F. Viral genomes that had GR-AR mutations collapsed as identical to the TF (red branch). An inner branching of the pseudo-tree (beige branch, #2) collapsed into a single leaf but remained 1nt different than the TF. Non-singlet branch population demonstrating an accumulation of mutations (sky blue branch, #1). The number in the parentheses indicate the number of viral genomes collapsed into the unique sequence.

**Figure S10.**
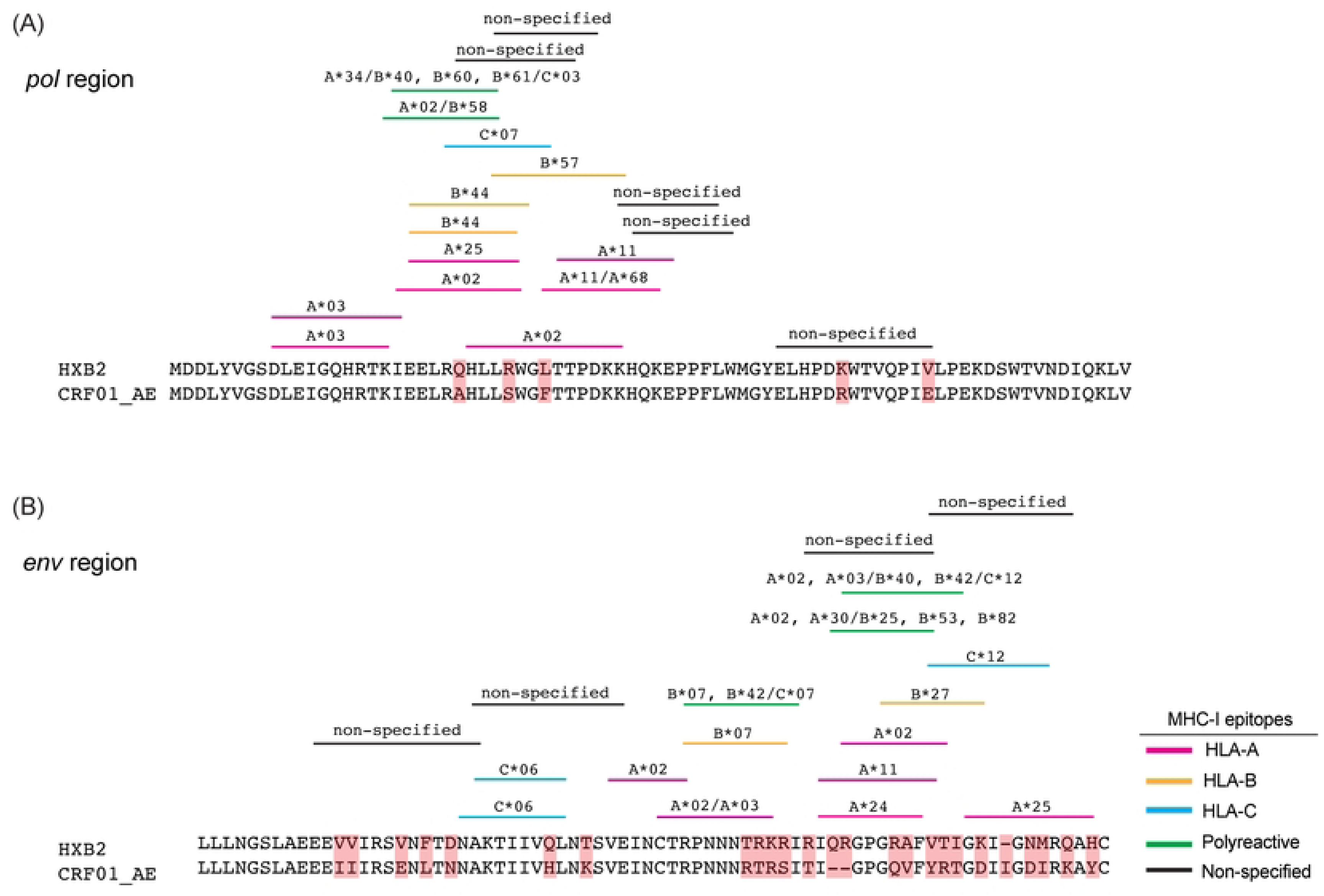
CTL epitope mapping for CRF01_AE. Mapping of CTL epitopes from HXB2 to CRF01_AE in (A) *pol* and (B) *env*. Epitopes are mapped to within a region of 14 amino acids or less. Epitopes were designated as either specific to HLA alleles (*e.g.*, A*02; A* pink/ B* orange/ C* blue bars), polyreactive (*e.g.*, A*02/B*58; green bars), or when no MHC presenting molecule is defined the host species is note (*e.g.*, human; black bars).

**Figure S11.**
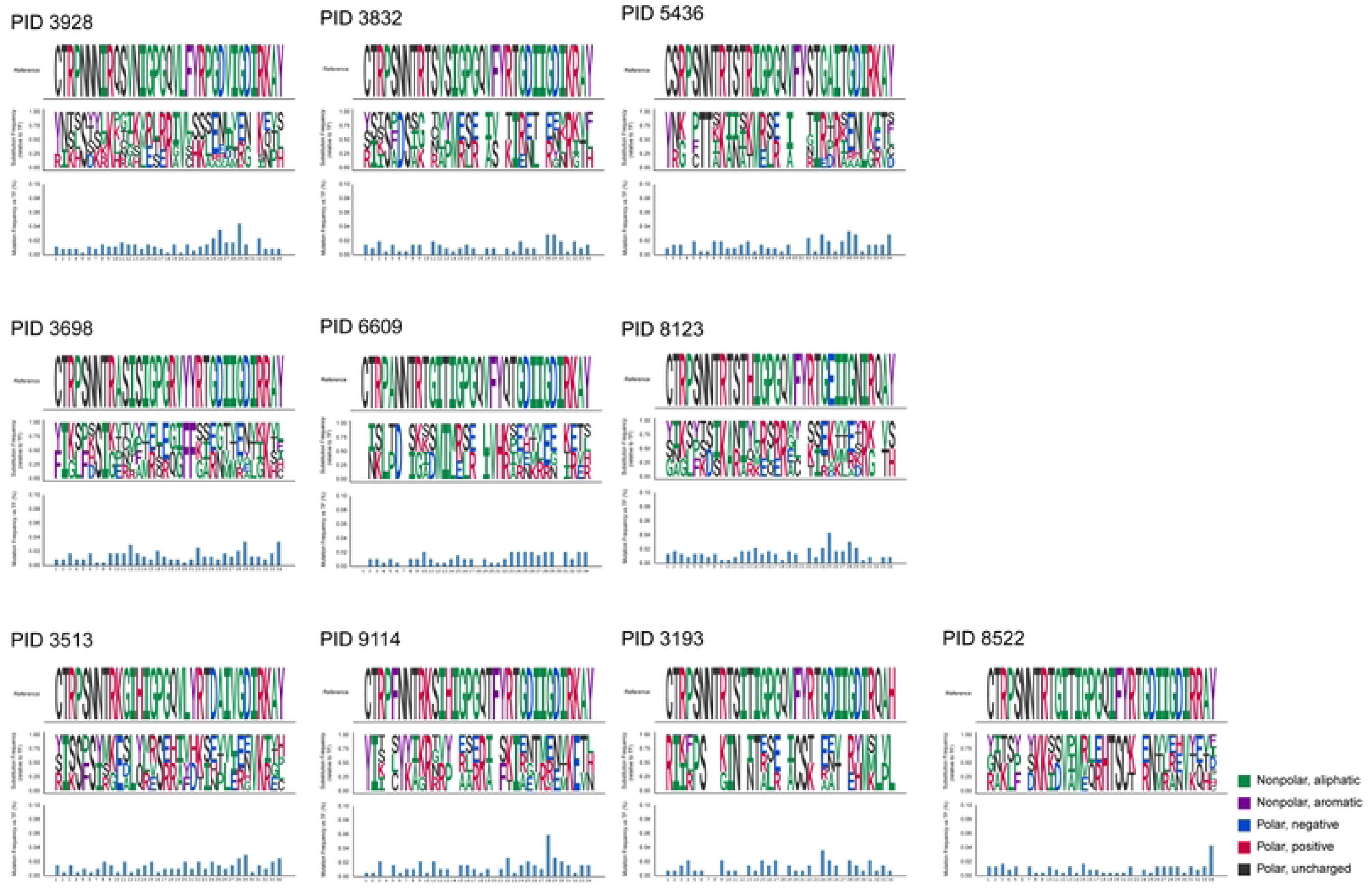
V3 loop mutations among single TF participants. Layer one (top) is the reference V3 loop to the TF for the PID shown. Layer two (middle) is the substitution frequency relative to the TF V3 loop sequence. Layer 3 (bottom) is the mutation frequency of the amino acid residues that differ from the transmitted/founder normalized to 100%. Where complete conservation is observed, no residue is shown in the bottom panel. Each participant ID is noted for each set of panels. Amino acid results were defined as nonpolar aliphatic (green: G (Gly), A (Ala), V (Val), L (Leu), I (Ile), M (Met), P (Pro)), nonpolar aromatic (purple: F (Phe), W (Trp), Y (Tyr)), polar uncharged (black: C (Cys), S (Ser), T (Thr), N (Asn), Q (Gln)), polar positivel charged (red: K (Lys), R (Arg), H (His)), and polar negativel charged (blue: D (Asp), E (Glu)).

**Figure S12.**
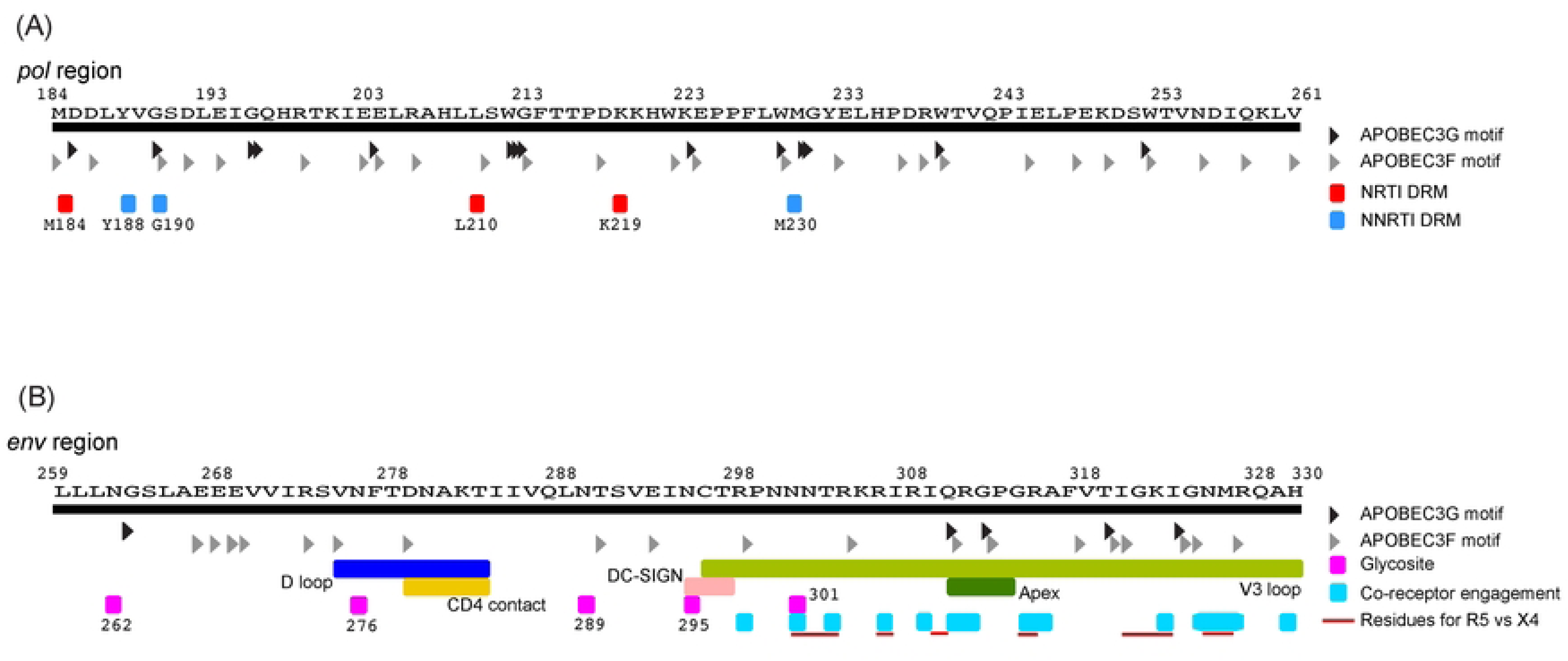
Illustrative map of sequenced subgenomic regions. (**A**) Amino acid sequence of *pol* from codon 184 to 261. Site of drug resistance mutations are denoted for NRTI (red) and NNRTI (bright blue). (**B**) Amino acid sequence of *env* from codon 259 to 330. The glycosite (magenta) and co-receptor engagement residues (cyan) are noted along with other features in this subgenomic regon of *env*. Residues found to be related to either R5 vs. X4 tropism are noted with a red line. APOBEC3G/F sites at the nucleotide motif level superimposed the amino acid sequence for APOBEC3G (GG>AG; black arrows) or APOBEC3F (GA>AA; grey arrows).

**Table S1.** Number of observed nucleotide mutations and inferred minimal single step rate.

| PID | *pol* (unfiltered dataset)<sup>a</sup> |  |  | *env* (unfiltered dataset)<sup>a</sup> |  |  |
| --- | --- | --- | --- | --- | --- | --- |
|  | # of genomes | Observed # of mutations (bp) | Inferred minimal single step rate (mut/site/day) | # of genomes | Observed # of mutations (bp) | Inferred minimal single step rate (mut/site/day) |
| 3928 | 19,251 | 2,921 | 2.82x10<sup>-5</sup> | 17,242 | 1,122 | 1.21x10<sup>-5</sup> |
| 3832 | 12,518 | 713 | 1.28x10<sup>-5</sup> | 12,993 | 508 | 8.79x10<sup>-6</sup> |
| 5436 | 7,766 | 763 | 1.83x10<sup>-5</sup> | 10,329 | 535 | 1.02x10<sup>-5</sup> |
| 3698<sup>c</sup> | 6,479 | 864 | 2.48x10<sup>-5</sup> |  |  |  |
| 3698<sup>d</sup> | 8,148 | 2,722 | 6.21x10<sup>-5</sup> | 10,120 | 590 | 1.18x10<sup>-5</sup> |
| 6609 | 10,386 | 912 | 1.63x10<sup>-5</sup> | 13,476 | 1,293 | 1.95x10<sup>-5</sup> |
| 8123 | 11,339 | 772 | 1.27x10<sup>-5</sup> | 13,867 | 1,011 | 1.37x10<sup>-5</sup> |
| 3513 | 12,913 | 590 | 8.49x10<sup>-6</sup> | 12,459 | 992 | 1.62x10<sup>-5</sup> |
| 9114 | 27,884 | 3,857 | 2.57x10<sup>-5</sup> | 10,957 | 501 | 9.16x10<sup>-6</sup> |
| 3193 | 3,704 | 224 | 1.12x10<sup>-5</sup> | 6,914 | 268 | 7.77x10<sup>-6</sup> |
| 8522 | 22,675 | 1,544 | 1.27x10<sup>-5</sup> | 12,602 | 622 | 9.17x10<sup>-6</sup> |
| **Median** | **11,929** | **308** | **1.46x10<sup>-5</sup>** | **12,531** | **606** | **1.10x10<sup>-5</sup>** |
| **[IQR]** | **[8,053–20,107]** | **[259–486]** | **[1.23x10<sup>-5</sup>–2.63x10<sup>-5</sup>]** | **[10,277–13,574]** | **[506–1,039]** | **[9.07x10<sup>-6</sup>–1.43x10<sup>-5</sup>]** |
| **p-value<sup>f</sup>** | **t(9)=5.00, p=0.0007** |  |  | **t(9)=10.47, p<0.0001** |  |  |
| **p-value<sup>g</sup>** | **t(9)=1.21 p=0.26** |  |  | **t(9)=2.25, p=0.051** |  |  |
| PID | *pol* (filtered dataset)<sup>b</sup> |  |  | *env* (filtered dataset)<sup>b</sup> |  |  |
|  | # of genomes | Observed # of mutations (bp) | Inferred minimal single step rate (mut/site/day) | # of genomes | Observed # of mutations (bp) | Inferred minimal single step rate (mut/site/day) |
| 3928 | 19,152 | 2,704 | 2.62x10<sup>-5</sup> | 17,220 | 1,065 | 1.15x10<sup>-5</sup> |
| 3832 | 12,496 | 660 | 1.19x10<sup>-5</sup> | 12,987 | 494 | 8.56x10<sup>-6</sup> |
| 5436 | 7,744 | 714 | 1.71x10<sup>-5</sup> | 10,319 | 514 | 9.84x10<sup>-6</sup> |
| 3698<sup>c</sup> | 6,624 | 1,154 | 3.24x10<sup>-5</sup> |  |  |  |
| 3698<sup>d</sup> | 8,137 | 1,209 | 2.76x10<sup>-5</sup> | 10,115 | 578 | 1.16x10<sup>-5</sup> |
| 6609 | 10,362 | 856 | 1.53x10<sup>-5</sup> | 13,473 | 1,285 | 1.94x10<sup>-5</sup> |
| 8123 | 11,313 | 715 | 1.17x10<sup>-5</sup> | 13,855 | 984 | 1.33x10<sup>-5</sup> |
| 3513 | 12,902 | 560 | 8.06x10<sup>-6</sup> | 12,452 | 978 | 1.60x10<sup>-5</sup> |
| 9114 | 27,723 | 3,438 | 2.30x10<sup>-5</sup> | 10,944 | 464 | 8.49x10<sup>-6</sup> |
| 3193 | 3,692 | 199 | 1.00x10<sup>-5</sup> | 6,911 | 261 | 7.57x10<sup>-6</sup> |
| 8522 | 22,636 | 1,448 | 1.19x10<sup>-5</sup> | 12,602 | 622 | 9.17x10<sup>-6</sup> |
| **Median<sup>e</sup>** | **11,905** | **715** | **1.36x10<sup>-5</sup>** | **12,527** | **600** | **1.07x10<sup>-5</sup>** |
| **[IQR]** | **[7,464–20,023]** | **[635–1,762]** | **[1.13x10<sup>-5</sup>–2.38x10<sup>-5</sup>]** | **[10,268–13,569]** | **[487–1,004]** | **[8.54x10<sup>-6</sup>–1.40x10<sup>-5</sup>]** |
| **p-value<sup>f</sup>** | **t(9)=4.76, p=0.001** |  |  | **t(9)=10.47, p<0.0001** |  |  |
| **p-value<sup>g</sup>** | **t(9)=0.14, p=0.89** |  |  | **t(9)=2.47, p=0.04** |  |  |

<sup>a</sup> Unfiltered dataset, where all viral genome sequences were analyzed<sup>b</sup> Filtered dataset, viral genome sequences with ≥2 GR-AR (APOBEC3G/F) were removed<sup>c,d</sup> Since PID 3698 has a potential 'Founder Effect', 3698<sup>c</sup> is where the C>T lineage sequences were removed, and 3698<sup>d</sup> maintains the C>T lineage viral genomes in analysis<sup>e</sup> The Median and IQR were calculated with the exclusion of PID 3698<sup>d</sup> where the C>T lineage was present for the uncollapsed dataset<sup>f</sup> PID 3698<sup>d</sup> dataset was excluded. Values were log<sub>10</sub>-transformed; One-sample t-test compared to HIV-1 RT single step rate of 3.0x10<sup>-5</sup> mut/site/day<sup>g</sup> PID 3698<sup>d</sup> dataset was excluded. Values were log<sub>10</sub>-transformed; One-sample t-test compared to HIV-1 RT single step rate of 1.4x10<sup>-5</sup> mut/site/day

**Table S2.** Inferred mutation rate based on the proportion of viral genomes identical to the TF.

| PID | *pol* subgenomic region |  |  | *env* subgenomic region |  |  |
| --- | --- | --- | --- | --- | --- | --- |
| | Proportion of TF<sup>a</sup> | Mean ($\lambda$)<sup>b</sup> | Inferred mutation rate (mut/site/day)<sup>c</sup> | Proportion of TF<sup>a</sup> | Mean ($\lambda$)<sup>b</sup> | Inferred mutation rate (mut/site/day)<sup>c</sup> |
| 3928 | 0.86 | 0.151 | 2.80x10<sup>-5</sup> | 0.94 | 0.062 | 1.15x10<sup>-5</sup> |
| 3832 | 0.95 | 0.051 | 1.15x10<sup>-5</sup> | 0.96 | 0.041 | 9.18x10<sup>-6</sup> |
| 5436 | 0.91 | 0.094 | 1.75x10<sup>-5</sup> | 0.85 | 0.163 | 3.02x10<sup>-5</sup> |
| 3698 | 0.87 | 0.139 | 2.59x10<sup>-5</sup> | 0.95 | 0.051 | 9.53x10<sup>-6</sup> |
| 6609 | 0.92 | 0.083 | 1.55x10<sup>-5</sup> | 0.91 | 0.094 | 1.75x10<sup>-5</sup> |
| 8123 | 0.94 | 0.062 | 1.15x10<sup>-5</sup> | 0.93 | 0.073 | 1.35x10<sup>-5</sup> |
| 3513 | 0.96 | 0.041 | 7.58x10<sup>-6</sup> | 0.92 | 0.083 | 1.55x10<sup>-5</sup> |
| 9114 | 0.87 | 0.139 | 2.59x10<sup>-5</sup> | 0.96 | 0.041 | 7.58x10<sup>-6</sup> |
| 3193 | 0.94 | 0.062 | 1.15x10<sup>-5</sup> | 0.96 | 0.041 | 7.58x10<sup>-6</sup> |
| 8522 | 0.94 | 0.062 | 1.15x10<sup>-5</sup> | 0.95 | 0.051 | 9.53x10<sup>-6</sup> |
| **Median** | **0.93** | **0.73** | **1.35x10<sup>-5</sup>** | **0.95** | **0.057** | **1.05x10<sup>-5</sup>** |
| **IQR** | **0.88–0.94** | **0.062–0.128** | **1.15x10<sup>-5</sup>–2.38x10<sup>-5</sup>** | **0.92–0.96** | **0.043–0.081** | **9.27x10<sup>-6</sup>–1.50x10<sup>-5</sup>** |
| ***p*-value<sup>d</sup>** | **t(9)=4.85, *p*=0.0009** |  |  | **t(9)=6.73, *p*<0.0001** |  |  |
| ***p*-value<sup>e</sup>** | **t(9)=0.61, *p*=0.57** |  |  | **t(9)=1.14, *p*=0.28** |  |  |

<sup>a</sup> Proportion of sequences that match the inferred single TF (Hamming distance of 0)<sup>b</sup> Mean of a Poisson distribution when $k=0$ (i.e., Hamming distance=0), $\lambda = -\ln(\Pr(0))$ <sup>c</sup> Median days corresponding to Fiebig stage used<sup>d</sup> Values were log<sub>10</sub>-transformed; One-sample t-test compared to HIV-1 RT single step rate of 3.0x10<sup>-5</sup> mut/site/day<sup>e</sup> Values were log<sub>10</sub>-transformed; One-sample t-test compared to HIV-1 RT single step rate of 1.4x10<sup>-5</sup> mut/site/day

**Table S3.** Summary of inferred minimum single step rate based on observed mutations, Poisson mean, and viral generation time sensitivity with assay error correction.

| Dataset type | Region | Inferred minimal single step rate based on Poisson mean ($\lambda$)<sup>a</sup> | Observed inferred minimal single step rate<sup>b</sup> | HIV-1 RT error rate with viral generation time sensitivity (assay error corrected) | | | | | |
| --- | --- | --- | --- | --- | --- | --- | --- | --- | --- |
| | | | | $\varepsilon = 3.0 \times 10^{-5}$ | | | $\varepsilon = 1.4 \times 10^{-5}$ | | |
| | | | | $\tau = 1.0^c$ | $\tau = 1.5^d$ | $\tau = 2.0^e$ | $\tau = 1.0^f$ | $\tau = 1.5^g$ | $\tau = 2.0^h$ |
| Unfiltered | *pol* | $1.35 \times 10^{-5}$<br>[$1.15 \times 10^{-5}$–$2.59 \times 10^{-5}$] | $1.46 \times 10^{-5}$<br>[$1.23 \times 10^{-5}$–$2.63 \times 10^{-5}$] | $4.30 \times 10^{-5}$ | $3.26 \times 10^{-5}$ | $\sim 2.80 \times 10^{-5}$ | $2.70 \times 10^{-5}$ | $2.22 \times 10^{-5}$ | $2.03 \times 10^{-5}$ |
| | *env* | $1.10 \times 10^{-5}$<br>[$8.99 \times 10^{-6}$–$1.75 \times 10^{-5}$] | $1.10 \times 10^{-5}$<br>[$9.07 \times 10^{-6}$–$1.43 \times 10^{-5}$] | | | | | | |
| Filtered | *pol* | $1.35 \times 10^{-5}$<br>[$1.15 \times 10^{-5}$–$2.59 \times 10^{-5}$] | $1.36 \times 10^{-5}$<br>[$1.13 \times 10^{-5}$–$2.38 \times 10^{-5}$] | | | | | | |
| | *env* | $1.10 \times 10^{-5}$<br>[$8.99 \times 10^{-6}$–$1.75 \times 10^{-5}$] | $1.07 \times 10^{-5}$<br>[$8.54 \times 10^{-6}$–$1.40 \times 10^{-5}$] | | | | | | |

All values are reported as the median [interquartile range] in mut/site/day
<sup>a</sup> Values from **Table S2**
<sup>b</sup> Values from **Table S1**
<sup>c</sup> Values from **Text S1 table 1**
<sup>d</sup> Values from **Text S1 table 3**
<sup>e</sup> Values from **Text S1 table 5**
<sup>f</sup> Values from **Text S1 table 2**
<sup>g</sup> Values from **Text S1 table 4**
<sup>h</sup> Values from **Text S1 table 6**

**Table S4.** The frequency of APOBEC3G/F-mediated activity among G>A mutations and all observed mutations.

| *pol* subgenomic regions |  |  |  |  |  |  |  |  |
| --- | --- | --- | --- | --- | --- | --- | --- | --- |
| PID | Total number of mutations | G→A mutations (%) | % G→A from APOBEC3G/F |  |  | % of all observed mutations |  |  |
|  |  |  | % A3G/F | % A3G | % A3F | % A3G/F | % A3G | % A3F |
| 3928 | 2,921 | 37.8 | 83.6 | 23.6 | 60.0 | 31.6 | 8.9 | 22.7 |
| 3832 | 713 | 42.2 | 86.6 | 12.4 | 74.2 | 36.5 | 5.2 | 31.3 |
| 5436 | 763 | 64.7 | 61.3 | 8.6 | 52.7 | 39.7 | 5.6 | 34.1 |
| 3698<sup>a</sup> | 864 | 19.2 | 78.5 | 17.8 | 60.7 | 15.1 | 3.4 | 11.7 |
| 3698<sup>b</sup> | 2,722 | 7.9 | 77.1 | 18.0 | 59.0 | 6.1 | 1.4 | 4.6 |
| 6609 | 912 | 49.5 | 84.5 | 33.9 | 50.6 | 41.8 | 16.8 | 25.0 |
| 8123 | 772 | 51.8 | 81.2 | 11.1 | 70.2 | 42.1 | 5.7 | 36.4 |
| 3513 | 590 | 49.5 | 75.9 | 18.2 | 57.7 | 37.6 | 9.0 | 28.6 |
| 9114 | 3,857 | 33.7 | 91.9 | 30.0 | 62.0 | 31.0 | 10.1 | 20.9 |
| 3193 | 224 | 46.6 | 89.2 | 18.6 | 70.6 | 41.6 | 8.7 | 32.9 |
| 8522 | 1,544 | 34.7 | 71.8 | 16.4 | 55.4 | 24.9 | 5.7 | 19.2 |
| **Median<sup>c</sup>** | **818** | **44.4** | **82.4** | **18.0** | **60.4** | **37.1** | **7.2** | **26.8** |
| **IQR<sup>c</sup>** | **682–1,888** | **34.5–50.1** | **74.9–87.3** | **12.0–25.2** | **54.7–70.3** | **29.5–41.6** | **5.5–9.3** | **20.5–33.2** |
| *env* subgenomic regions |  |  |  |  |  |  |  |  |
| PID | Total number of mutations | G→A mutations (%) | % G→A from APOBEC3G/F |  |  | % of all observed mutations |  |  |
|  |  |  | % A3G/F | % A3G | % A3F | % A3G/F | % A3G | % A3F |
| 3928 | 1,122 | 41.6 | 77.6 | 15.4 | 62.2 | 32.3 | 6.4 | 25.8 |
| 3832 | 508 | 43.3 | 57.3 | 12.3 | 45.0 | 24.8 | 5.3 | 19.5 |
| 5436 | 535 | 44.9 | 64.6 | 11.5 | 53.1 | 29.0 | 5.2 | 23.9 |
| 3698 | 590 | 25.4 | 78.7 | 12.5 | 66.2 | 20.0 | 3.2 | 16.8 |
| 6609 | 1,293 | 20.7 | 75.7 | 16.2 | 59.5 | 15.6 | 3.3 | 12.3 |
| 8123 | 1,011 | 57.9 | 94.3 | 66.4 | 27.8 | 54.6 | 38.5 | 16.1 |
| 3513 | 992 | 21.0 | 76.5 | 14.2 | 62.3 | 16.1 | 3.0 | 13.1 |
| 9114 | 501 | 46.1 | 87.2 | 11.9 | 75.3 | 40.2 | 5.5 | 34.7 |
| 3193 | 268 | 36.0 | 80.9 | 18.0 | 62.9 | 29.1 | 6.5 | 22.6 |
| 8522 | 622 | 29.0 | 78.6 | 19.3 | 59.3 | 22.7 | 5.6 | 17.2 |
| **Median** | **606** | **38.8** | **78.1** | **14.8** | **60.8** | **26.9** | **5.4** | **18.3** |
| **IQR** | **506–1,039** | **24.3–45.2** | **72.8–82.5** | **12.2–18.3** | **51.1–63.7** | **19.0–34.2** | **3.3–6.4** | **15.4–24.4** |
| % G→A from APOBEC3G/F: % A3G vs. % A3F *p*-value<sup>d</sup> |  |  |  |  |  |  | *pol* | 0.001 |
|  |  |  |  |  |  |  | *env* | 0.003 |
| % observed mutations: % A3G vs. % A3F *p*-value<sup>d</sup> |  |  |  |  |  |  | *pol* | 0.002 |
|  |  |  |  |  |  |  | *env* | 0.06 |

<sup>a,b</sup> Since PID 3698 has a potential 'Founder Effect', 3698<sup>a</sup> is where the C>T lineage sequences were removed, and 3698<sup>b</sup> maintains the C>T lineage viral genomes in analysis
<sup>c</sup> The Median and IQR were calculated with the exclusion of PID 3698<sup>b</sup> where the C>T lineage was present for the uncollapsed dataset
<sup>d</sup> Wilcoxon matched-pairs signed rank test

**Table S5.** Predicted coreceptor tropism of inferred single transmitted/founder virus by genetic model training of known CRF01_AE genotypes.

| Sequence/<br>Participant ID | V3 loop<sup>a,b</sup> | Net charge | Training model<br>*Population % total (% unique)* |  |  |
| --- | --- | --- | --- | --- | --- |
|  |  |  | TF tropism | R5 | X4 |
| Con. CRF01_AE | CT**R**PSNNT**R**T**S**ITIGPGQV-FY**R**T**G****D**I**I****G****D**I**R**KAY C | +3 | R5 | - | - |
| 3928 | CT**R**PNNNI**R**Q**S**VNIGPGQVL**F**Y**R**P**G****D**V**I****G****D**I**R**KAY (C) | +3 | R5 | 99.10 (89.7) | 0.90 (10.3) |
| 3832 | CT**R**PSNNT**R**T**S**V**S**IGPGQV-FY**R**T**G****D**I**I****G****D**I**K**RAY (C) | +3 | R5 | 99.99 (99.5) | 0.01 (0.5) |
| 5436 | CS**R**PSNNT**R**T**S**T**R**IGPGQV-FYST**G**A**I**T**G****D**I**R**KAY (C) | +4 | R5 | 99.94 (98.1) | 0.06 (1.9) |
| 3698 | CT**R**PSNNT**R**A**S**ISIGPG**R**V-YY**R**T**G****D**I**I****G****D**I**R**RAY (C) | +4 | R5 | 99.62 (91.6) | 0.38 (8.4) |
| 6609 | CT**R**PANNT**R**T**G**ITIGPGQV-FY**Q**T**G****D**I**I****G****D**I**R**KAY (C) | +2 | R5 | 100.00 (100) | 0.00 (0) |
| 8123 | CT**R**PSNNT**R**T**S**T**H**IGPGQV-FY**R**T**G****E**I**I****G**N**I**RQAY (C) | +4 | R5 | 99.50 (88.7) | 0.50 (11.3) |
| 3513 | CT**R**PSNNT**R**K**G**I**H**IGPGQVL-Y**R**T**D**A**I**V**G****D**I**R**KAY (C) | +5 | R5 | 99.57 (88.1) | 0.43 (11.9) |
| 9114 | CT**R**PFNNT**R**K**S**I**H**IGPGQT-FY**R**T**G****D**I**I****G****D**I**R**KAY (C) | +5 | X4 | 0.31 (8.1) | 99.69 (91.9) |
| 3193 | CT**R**PSNNT**R**T**S**ITIGPGQV-FY**R**T**G****D**I**I****G****D**I**R**QAH (C) | +3 | R5 | 99.99 (99.3) | 0.01 (0.7) |
| 8522 | CT**R**PSNNT**R**T**G**ITIGPGQ**I**-FY**R**T**G****D**I**I****G****D**I**R**RAY (C) | +3 | R5 | 99.98 (99.6) | 0.02 (0.4) |

<sup>a</sup> V3 loop residues are denoted as negative charged (blue), positive charged (red), non-charged (black). HXB2 codons 254 to 331 of the amplicon sequenced. The final residue of the V3 loop (Cys, C) was not captured, a placeholder (C) is included
<sup>b</sup> The 11<sup>th</sup> and 25<sup>th</sup> positions are in bold

**Table S6.** Frequency of detected drug resistance across viral genomes that possess at least one drug resistance mutations in inferred single TF virus participants.

| PID | % Frequency observed among viral genomes with at least one drug resistance mutation |  |  |  |  |  |  |  |  |  |
| --- | --- | --- | --- | --- | --- | --- | --- | --- | --- | --- |
|  | NRTI |  |  |  |  | NNRTI |  |  |  |  |
|  | M184I<sup>a</sup><br>ATG→ATA | M184V<br>ATG→GTG | L210W<br>TTG→TGG | K219E<br>AAA→GAA | K219Q<br>AAA→CAA | Y188C<br>TAT→TGT | Y188H<br>TAT→CAT | G190A<br>GGA→GCA | G190E<sup>b</sup><br>GGA→GAA | M230L<br>ATG→YTG |
| 3928 | 4.03 | - | 0.21 | 0.21 | - | 0.21 | 0.21 | - | 1.69 | 0.21 |
| 3832 | 0.46 | - | - | - | - | - | - | - | 1.83 | - |
| 5436 | 0.97 | - | - | - | - | - | - | 0.49 | 3.40 | - |
| 3698 | 0.35 | - | - | - | - | - | 0.69 | - | 0.69 | - |
| 6609 | 1.12 | - | - | - | 0.37 | - | - | - | 1.49 | - |
| 8123 | 1.26 | 0.42 | - | - | - | 0.42 | 0.42 | - | 0.84 | 0.42 |
| 3513 | 1.74 | - | - | 0.43 | - | 0.43 | - | 0.43 | 1.74 | - |
| 9114 | 3.42 | - | - | - | - | - | 0.23 | 0.23 | 3.65 | - |
| 3193 | - | 1.04 | - | - | - | - | 1.04 | - | 1.04 | - |
| 8522 | 1.14 | - | - | - | - | 0.28 | - | - | 2.85 | 0.58 |

<sup>a</sup> Very common APOBEC3G mutation<sup>b</sup> Frequent APOBEC3F mutation

